# Differentiation latency, cell division symmetry, and dormancy signatures define fetal liver HSCs at single cell resolution

**DOI:** 10.1101/2023.06.01.543314

**Authors:** Takashi Ishida, Juliane Mercoli, Adam M. Heck, Ian Phelps, Barbara Varnum-Finney, Stacey Dozono, Cynthia Nourigat-McKay, Katie Kraskouskas, Rachel Wellington, Olivia Waltner, Dana L Jackson, Colleen Delaney, Shahin Rafii, Irwin D. Bernstein, Kimberly A. Aldinger, Birth Defects Research Laboratory (BDRL), Cole Trapnell, Helong G. Zhao, Brandon Hadland

## Abstract

Decoding the gene regulatory mechanisms and signaling interactions that orchestrate the self-renewal of hematopoietic stem cells (HSCs) during their expansion in the fetal liver (FL) could unlock novel therapeutic strategies to expand transplantable HSCs, a long-standing challenge. Here, to explore intrinsic and extrinsic regulation of FL-HSC self-renewal at the single cell level, we engineered a culture platform designed to recapitulate the FL endothelial niche, which supports the ex vivo amplification of serially engraftable HSCs. Leveraging this platform in combination with single cell index flow cytometry, live imaging, serial transplantation assays, and single cell RNA-sequencing, we uncovered previously unrecognized heterogeneity within immunophenotypically defined FL-HSCs. Specifically, we demonstrated that differentiation latency, symmetric cell divisions, and transcriptional signatures of biosynthetic dormancy and lipid metabolism are distinguishing properties of rare FL-HSCs capable of serial, long-term multilineage hematopoietic reconstitution. Our findings support a paradigm in which intrinsic programs and extrinsic signals combinatorially facilitate the symmetric self-renewal and expansion of nascent HSCs in the FL niche while delaying their active participation in hematopoiesis. Additionally, our study provides a valuable resource for future investigations into the intrinsic and niche-derived signaling pathways that govern FL-HSC self-renewal.

## Introduction

Hematopoietic stem cells (HSCs) have the distinguishing properties of life-long self-renewal and the ability to reconstitute multilineage hematopoiesis upon transplantation. Understanding the biology of HSC development is essential to unlocking methods for de novo HSC generation and expansion to facilitate advances in hematopoietic stem cell transplantation, gene therapies, and disease modeling for blood and immune disorders.

Following their initial emergence in embryonic arteries, HSCs seed the fetal liver (FL), where they have conventionally been thought to both rapidly expand (generating sufficient HSCs to sustain life-long hematopoiesis in the adult) and differentiate (generating mature blood cells essential to support the developing embryo). To facilitate these processes, FL-HSCs have been postulated to exist in a state of proliferative and biosynthetic activation during prenatal development, followed by a transition to a state of biosynthetic dormancy and cell cycle quiescence upon their migration to the bone marrow (BM) perinatally.^1,2^

Several elegant studies using methodologies to track the fate of nascent HSCs and progenitors through development have recently called for a revision of this classical paradigm of developmental hematopoiesis. These studies suggest that distinct waves of HSC-independent progenitors also seed the fetal liver and serve as the primary source of mature blood cells throughout prenatal development, with minimal contribution from HSCs, which are instead reserved to contribute primarily to adult hematopoiesis.^3^ Further supporting this novel paradigm, we and others have uncovered distinct ontogenies for oligopotent and even multipotent hematopoietic progenitors that precede HSCs during various waves of endothelial to hematopoietic transition in embryonic blood vessels.^3–5^ While some HSC-independent progenitors possess properties that overlap with HSCs, such as multilineage hematopoietic activity, they are generally lacking in durable self-renewal capacity assayed by serial transplantation, the gold standard methodology for defining functional HSCs of relevance for translational applications.

Recent studies also suggest that the number of precursors emerging in the embryo that contribute to hematopoiesis in the adult when traced in vivo is substantially larger than the number of engraftable HSCs detected by direct transplantation assays^6,7^ and that this compartment of long-lived hematopoietic precursors undergoes only limited expansion in the fetal liver.^8^ Lineage barcoding studies in vivo suggest significant heterogeneity in this pool of long-lived precursors in the FL, identifying a distinct population of embryonic multipotent progenitors (eMPPs) independent of HSC origin. Remarkably, eMPPs were found to contribute to mature blood cells in the adult when traced in situ, but lack long-term engraftment properties in transplantation assays essential to classify them as bona fide HSCs.^9^ In contrast, another study identified a subset of lymphoid-biased “developmentally-restricted HSCs” in the FL possessing long-term engraftment upon transplantation, yet failing to contribute to adult hematopoiesis in situ,^10^ in line with previous work identifying populations of FL-HSCs with distinct engraftment properties.^11^

Altogether, these reports highlight substantial heterogeneity of the HSCs and HSCs-independent progenitors that seed the fetal liver in early gestation and contribute to various aspects of hematopoiesis during prenatal development and into adulthood. Though prior studies have identified surface markers that enable substantial enrichment of engraftable FL-HSCs by FACS to study their properties at a population level, no set of markers has been shown to achieve functional HSC purity. Thus, uncovering the unique phenotypic, functional, and molecular properties of the rare subset with long-term, multilineage engraftment capacity requires studies at a single cell level. Furthermore, a better understanding of how functionally engraftable HSCs undergo maturation and self-renewal in the fetal liver niche could provide insight into engineering more robust protocols for generation of engraftable HSCs in vitro.

To address these issues at single cell level, we developed a serum-free culture platform that mimics the FL vascular niche, which enabled ex vivo amplification of serially engrafting HSCs from individual immunophenotypic FL-HSCs isolated by FACS at early and mid-gestation in the mouse embryo. We leveraged this platform to study the heterogeneity of the FL-HSC compartment by integrating index flow cytometry, live imaging, single cell RNA-sequencing (scRNAseq), and transplantation assays, revealing the distinguishing properties of serially engraftable FL-HSCs and niche signals that contribute to their self-renewal. Altogether, these studies provide a valuable resource for ongoing efforts to understand the ontogeny of functionally transplantable HSCs and recapitulate their development and expansion ex vivo.

## Results

### Establishment of a clonal HSC amplification platform reveals unique properties of serially engrafting FL-HSCs

Aiming to study FL-HSCs at single cell resolution, we first sought to define an immunophenotype sufficient to highly enrich for long-term engrafting HSC activity in both the early (embryonic day (E) 13.5) and mid-gestation (E15.5/16.5) FL at the population level. Using stringent gating for known FL-HSC markers, Endothelial Cell Protein C Receptor (EPCR) and Stem Cell Antigen-1 (SCA1),^12–15^ we found that CD45^+^DAPI^-^GR1^-^F4/80^-^SCA1^high^EPCR^high^ immunophenotype (hereafter denoted SE^hi^) efficiently excluded cells expressing other lineage markers as well as CD48, allowing us to simplify the sorting methodology for immunophenotypic FL-HSCs (Figures S1A and B). Since CD150 is not a specific marker of embryonic HSCs until E14.5,^16–18^ we excluded CD150 from these initial studies encompassing the early FL. As expected, the SE^hi^ population from both the early and mid-gestation FL provides long-term, multilineage hematopoietic engraftment following transplantation into congenic strain adult mice (Figure S1C).

Next, to evaluate functional heterogeneity within the immunophenotypically defined SE^hi^ FL-HSC fraction at the single cell level, we established a coculture platform recapitulating the FL endothelial niche (Figure 1A). Briefly, FL-derived endothelial cells (ECs) were transduced with a lentivirus encoding constitutively active AKT1 (myristoylated AKT1; MyrAKT) (henceforth “FL-AKT-ECs”), which enables EC propagation in serum-free culture while maintaining their endogenous properties, as previously described.^19,20^ We first tested immunophenotypically defined HSCs isolated from E15.5/16.5 FL, a developmental stage when engrafting HSCs are most abundant in the FL and are further enriched based on CD150 expression.^16,21^ SE^hi^CD150^+^ FL-HSCs were individually index sorted into one well of a 96 well-plate containing FL-AKT-ECs in serum-free media with hematopoietic cytokines (SCF and TPO). In initial experiments, the formation of hematopoietic colonies in coculture was monitored visually over time by microscopy, and a portion of each colony at the time it was initially observed (day 7, 12, or 14) was harvested by pipetting for flow cytometric analysis. Following coculture, most colonies contained cells retaining co-expression of HSC markers EPCR and SCA1 (Figure 1B), suggesting maintenance and expansion of HSCs from single SE^hi^ FL cells during culture.^13,14,22^ Interestingly, a subset of late-emerging colonies (detected on day 12 and beyond) consisted nearly exclusively of HSC-like SCA1^+^EPCR^+^ cells, suggesting slower cell cycle kinetics, symmetric expansion, and differentiation latency of this subset during EC coculture (Figure 1B, left panel; Figure S1D). In contrast, most colonies detectable at day 12 and earlier consisted of a mixed population of SCA1^+^EPCR^+^ HSC-like cells and cells lacking SCA1 and/or EPCR consistent with differentiation to hematopoietic progenitors (Figure 1B, right panel; Figure S1D). These studies reveal significant heterogeneity of immunophenotypic CD150^+^SE^hi^ FL-HSC behavior when assayed at the single cell level, suggesting intrinsic differences in proliferative potential and propensity for self-renewal verses differentiation during FL-AKT-EC coculture. So as not to exclude late-emerging colonies, colonies were harvested between day 12 and 15 for subsequent experiments.

**Figure 1:**
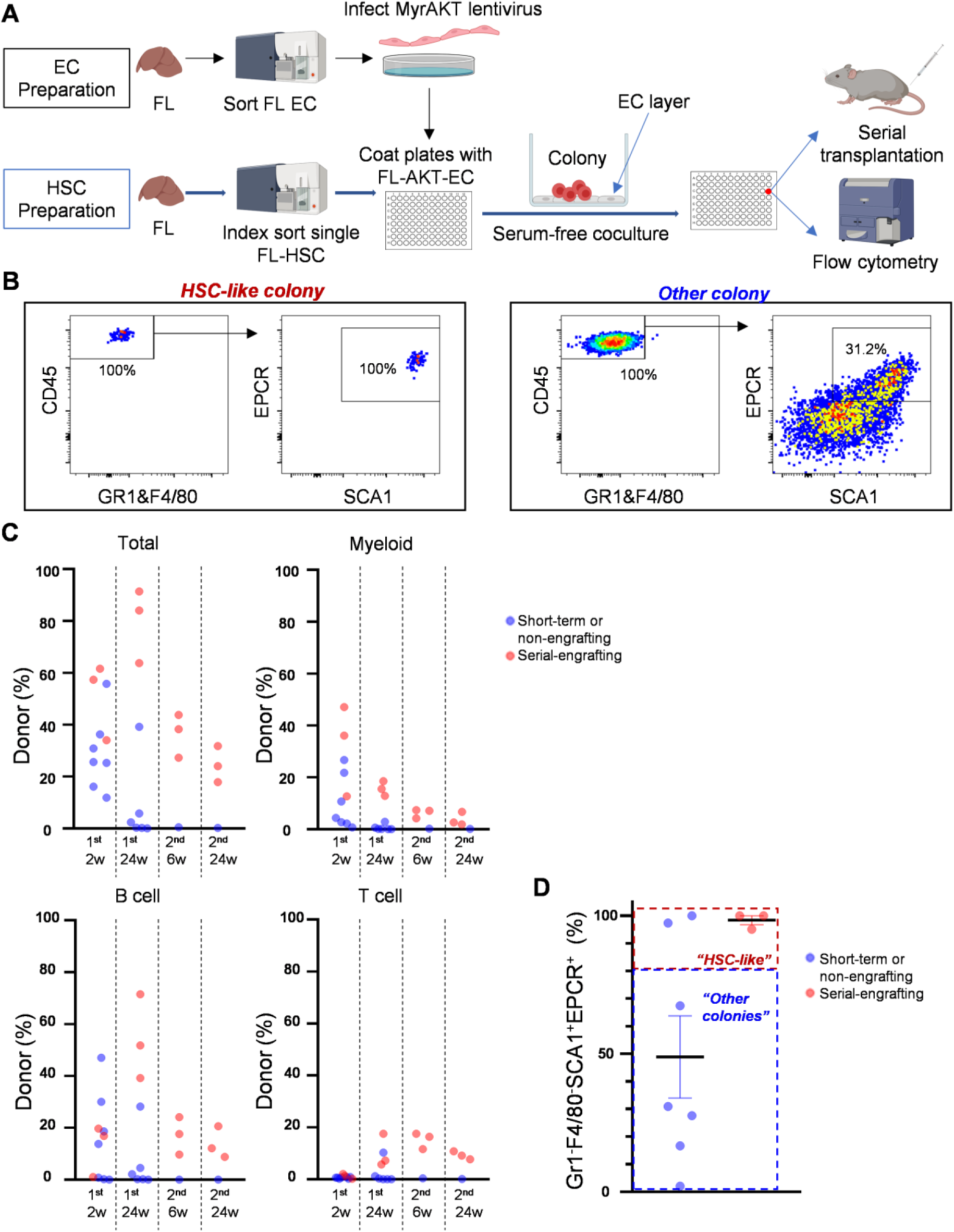
Establishment of a FL vascular niche platform supporting ex vivo amplification of clonal FL-HSCs. **(A)** Overview of methodology. Sorted FL-derived ECs were infected with lentivirus encoding myristoylated AKT (myrAKT) to generate FL-AKT-ECs. Single FL-HSCs were sorted into each well of a 96 well-plate coated with FL-AKT-EC in serum free media with hematopoietic cytokines (SCF, TPO). Following a period of coculture, emerging colonies were assessed by flow cytometry for immunophenotype and by serial transplantation to measure long-term hematopoietic engraftment (See Materials & Methods for details). **(B)** Representative immunophenotypes of colonies emerging from single E16.5 SE^hi^CD150^+^ FL-HSC following coculture with FL-AKT-EC. Colonies were analyzed on day 15 by flow cytometry. Shown is expression of EPCR and SCA1 within cells gated as CD45^+^GR1^-^F4/80^-^, after exclusion of dead cells (DAPI^+^) and VE-cadherin^+^ FL-AKT-EC. **(C)** Donor chimerism (total, myeloid, B cell, and T cell) in peripheral blood of primary and secondary recipients transplanted with the progeny of single E16.5 FL-HSCs following coculture with FL-AKT-ECs. 50% of cells from each colony were analyzed by flow cytometry and a portion of the remaining cells (25-50%) was used for transplantation (n=10). **(D)** Frequency of GR1^-^F4/80^-^SCA1^+^EPCR^+^ cells (amongst total viable CD45^+^ cells) in individual colonies possessing serial multilineage engraftment (red) (n=3) or lacking serial multilineage engraftment (blue) (n=7). “HSC-like” colonies were defined as colonies with frequency of >80% GR1^-^F4/80^-^SCA1^+^EPCR^+^ amongst total viable CD45^+^ cells (defined to encompass all colonies possessing serial multilineage engraftment).

Next, the functional engraftment properties of the progeny of single E15.5/16.5 SE^hi^CD150^+^ FL-HSCs following coculture was assessed by correlating their immunophenotype (using 50% of cells) with serial transplantation (using the remaining 50% of cells). Although at least short-term engraftment was observed from most colonies tested, colonies providing serial engraftment were exclusively from the subset consisting of nearly homogenous SCA1^+^EPCR^+^ HSC-like cells (Figure 1C-D). Similar results were observed when assessing single SE^hi^ cells from the early E13.5 FL in FL-AKT-EC coculture (Figure 2A), with serial engraftment detected only from a subset of colonies consisting predominantly (>80%) of SCA1^+^EPCR^+^ cells (Figure 2B), which we hereafter refer to as “HSC-like colonies.” At both developmental stages, HSC-like colonies contained nearly 100-fold lower average number of total hematopoietic (CD45^+^) cells compared with other colony types (Figure 2C), consistent with relatively lower cell division kinetics. Notably, HSC-like colonies were less frequently generated from SE^hi^ FL cells at E13.5 (mean 17.8% of total SE^hi^ cells sorted) compared with E15.5/16.5 (mean 37.9% of total SE^hi^ cells sorted) (Figure 2D). Unlike at E15.5/16.5, E13.5 FL SE^hi^ cells with HSC-like colony potential did not consistently express CD150 (Figures 2E, S1E-F). ESAM (endothelial cell adhesion molecule), another HSC marker,^23^ was expressed ubiquitously in FL SE^hi^ cells and thus failed to enrich for HSC-like colony potential at either E13.5 or E15.5/E16.5 (Figures 2E, S1E-F).

**Figure 2:**
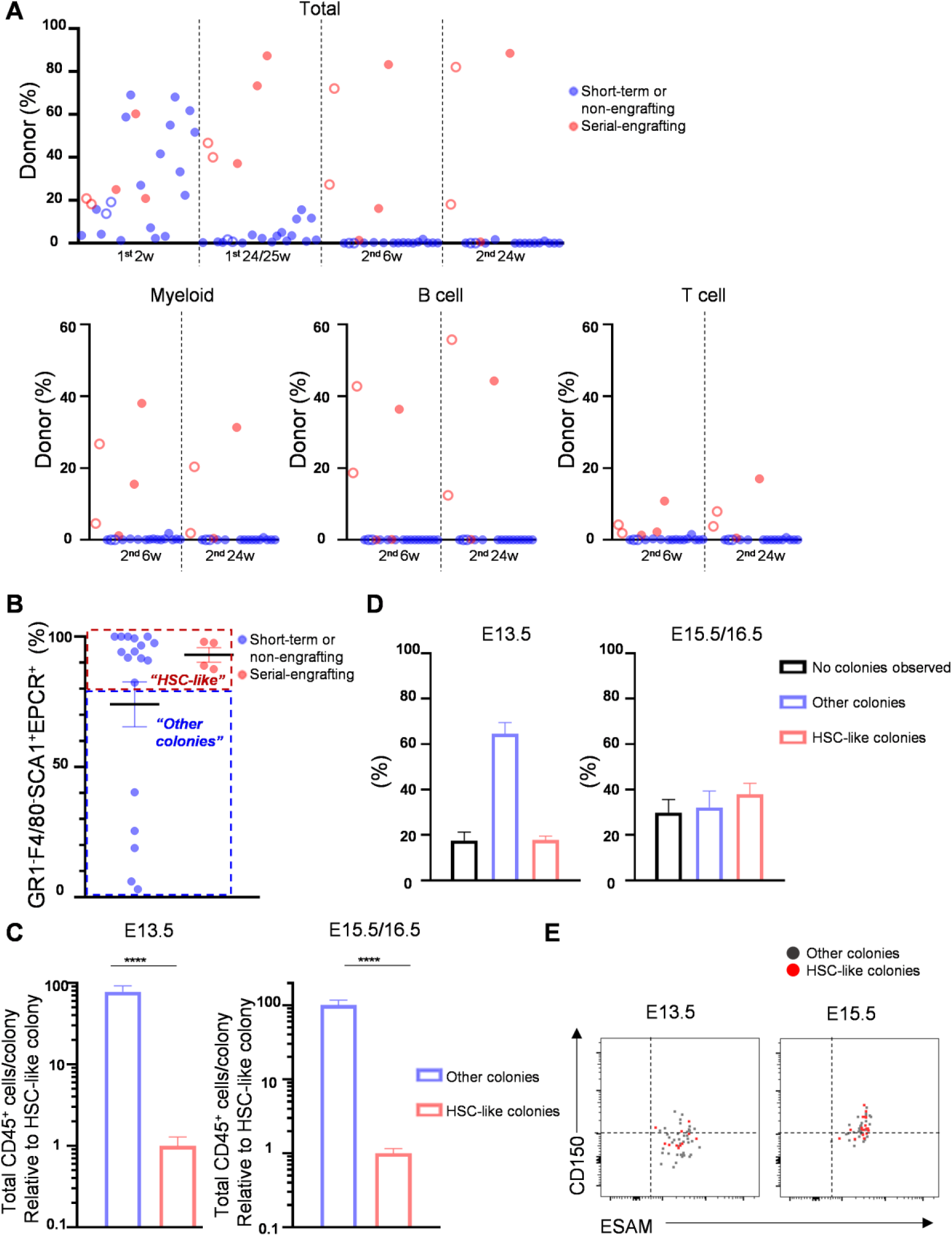
SE^hi^ FL-HSCs demonstrate heterogeneity in functional engraftment, proliferation, and immunophenotype following FL-AKT-EC coculture. **(A)** Donor chimerism (total, myeloid, B cell, and T cell) in peripheral blood of primary and secondary recipients transplanted with the progeny of single E13.5 SE^hi^ FL-HSCs following coculture with FL-AKT-ECs. 50% of cells from each colony were analyzed by flow cytometry and the remaining cells were used for transplantation (n=22 total colonies transplanted to n=24 total recipients; results pooled from 3 independent experiments). Two colonies (open circles) were each transplanted to 2 recipients. **(B)** Frequency of GR1^-^F4/80^-^SCA1^+^EPCR^+^ cells (amongst total viable CD45^+^ cells) in individual colonies possessing serial multilineage engraftment (red) (n=4) or lacking serial multilineage engraftment (blue) (n=18). “HSC-like” colonies were defined as colonies with frequency of >80% GR1^-^F4/80^-^SCA1^+^EPCR^+^ amongst total viable CD45^+^ cells (defined to encompass all colonies possessing serial multilineage engraftment). **(C)** Total viable CD45+ cells per colony determined by flow cytometry following coculture of single E13.5 or E15.5/16.5 SE^hi^ FL-HSCs with FL-AKT-ECs. Cell numbers were normalized to HSC-like colonies for each experiment. (E13.5: n=32 HSC-like colonies, n=119 other colonies, from 3 independent experiments; E15.5/16.5: n=95 HSC colonies, n=80 other colonies, from 5 independent experiments). ****P<0.0001 (Mann-Whitney test). Data are expressed as mean±SEM. **(D)** Frequency of colony types observed following coculture of single E13.5 (3 independent experiments) or E15.5/16.5 SE^hi^ (5 independent experiments) FL-HSCs with FL-AKT-ECs. Data are expressed as mean±SEM. **(E)** Expression of ESAM and CD150 by index analysis of E13.5 (left) and E15.5 (right) SE^hi^ FL-HSCs giving rise to immunophenotypic HSC-like colonies (red) or other colonies (gray) following FL-AKT-EC coculture.

Across multiple independent experiments, the average frequencies of HSC-like colonies generated was: 1 in 5.7 index sorted SE^hi^ cells at E13.5 (range 1 in 4.7 to 1 in 6.5, n=3 experiments) and 1 in 2.3 index sorted SE^hi^CD150^+^ cells at E15.5/16.5 (range 1 in 1.7 to 1 in 4.3, n=5 experiments). Notably, this latter frequency is similar to a previous study, which reported an HSC frequency of 1 in 2.7 CD150^+^CD48^+^Sca1^+^Lineage^-^Mac-1^+^ cells in the E14.5 FL, based on detection of multilineage engraftment at 16 weeks after limiting dilution transplantation in primary recipients.^16^ By more rigorously testing for self-renewing HSC potential by serial transplantation, our current studies suggest the true frequency of bona fide HSCs is significantly lower than that observed by assessing primary transplantation only, which may include short-term engrafting populations. Measuring HSC potential based on serial engraftment, we predict FL-HSC frequency to be approximately 1 in 17 SE^hi^ cells at E13.5 and 1 in 5.8 SE^hi^CD150^+^ FL cells at E15.5/16.5. Together, these studies suggest significant heterogeneity in functional engraftment capacity in even the most stringently immunophenotypically defined FL-HSC populations.

### Maintenance of ESAM expression identifies HSC-like colonies with serial engraftment potential

Since the SCA1^+^EPCR^+^ HSC-like colonies remain heterogeneous in their engraftment properties (Figure 1D, 2B), we next sought to identify additional surface proteins that may serve as predictive markers of their serial engraftment capacity. Although ESAM was ubiquitously expressed on fresh SE^hi^ FL cells independent of HSC potential (Figure 2E, Figures S1E-F), we hypothesized that maintenance of ESAM expression following in vitro culture might identify SCA1^+^EPCR^+^ HSC-like colonies with serial engraftment based on prior studies suggesting ESAM as a marker of HSC activity in vitro.^13^ To test this, we sorted single E15.5 SE^hi^CD150^+^ FL-HSCs for coculture with FL-AKT-EC (Figure 3A). Following coculture, we observed heterogeneity of ESAM expression amongst identified SCA1^+^EPCR^+^ HSC-like colonies (Figure 3B). When we transplanted the remaining cells from each of these HSC-like colonies, we observed long-term, serial multilineage engraftment in HSC-like colonies containing predominantly (>60%) ESAM^+^ cells (hereafter referred to as ESAM^+^ HSC colonies), but not from HSC-like colonies containing predominantly ESAM^-^ cells (Figures 3B and 3C). In an independent experiment from E15.5 SE^hi^CD150^+^ FL-HSCs, ESAM expression in HSC-like colonies also predicted high level donor engraftment of immunophenotypic HSCs (Lin^-^SCA1^+^KIT^+^CD150^+^CD48^-^) in bone marrow at 24 weeks post-transplant (Figure S2). Notably, when cells from an individual ESAM^+^ HSC colony were transplanted at limiting dilutions, secondary engraftment was observed in multiple recipients, indicating the FL-AKT-EC niche supported expansion of functional long-term ESAM^+^ HSCs from single FL-HSCs (approximately 15 serial-engrafting HSCs generated, range 5-43) (Figure S3).

**Figure 3:**
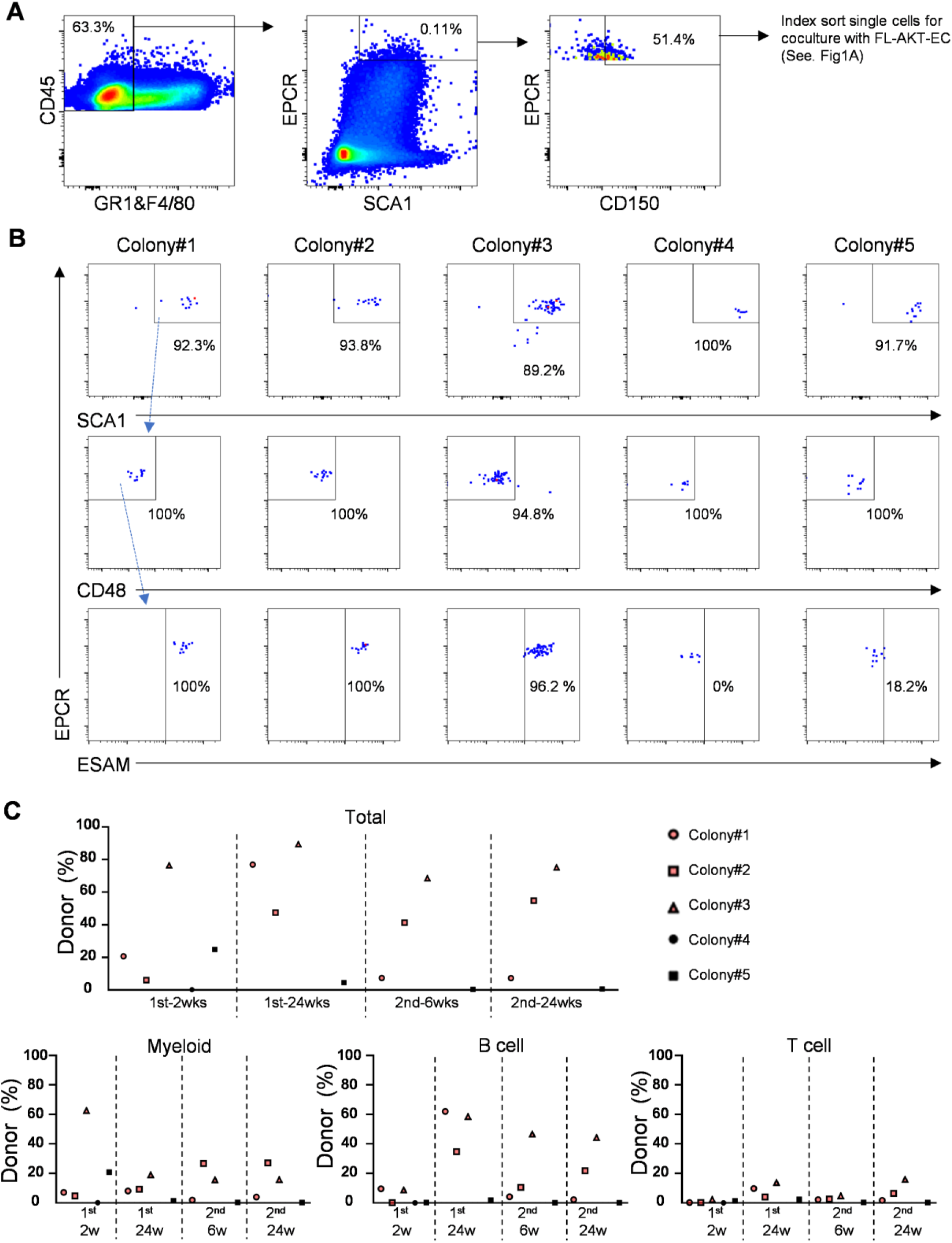
ESAM-expressing HSC colonies exhibit serial hematopoietic reconstitution capacity. **(A)** Single cell sorting strategy of E15.5 SE^hi^CD150^+^ FL-HSCs for coculture with FL-AKT-EC. **(B)** Expression of CD48 and ESAM in 5 HSC-like colonies analyzed by flow cytometry following 15 days of coculture. 50% of the cells from each colony were used for flow cytometry and the remaining cells were used for transplantation. (HSC-like colonies were defined as containing >80% CD48^-^SCA1^+^EPCR^+^ cells amongst total viable CD45^+^ cells). **(C)** Donor chimerism (total, myeloid, B cell, and T cell) in peripheral blood of primary and secondary recipients transplanted with cells from each of the HSC-like colonies shown in B (n=5).

Altogether, these studies establish a novel assay to assess the immunophenotypic properties and engraftment potential of FL-HSCs at single cell resolution in a developmentally relevant ex vivo niche. Specifically, this approach revealed functional heterogeneity in both the immunophenotypic E13.5 SE^hi^ and E15.5/16.5 SE^hi^CD150^+^ FL-HSC populations at the single cell level based on their differential capacity to generate serially engrafting ESAM^+^ HSC colonies during FL-AKT-EC coculture. Based on homogenous ESAM^+^SCA1^+^EPCR^+^ immunophenotype and clonal expansion of serially engrafting HSCs during FL-AKT-EC coculture, these results further suggest that long-term engrafting FL-HSCs are distinctly characterized by latency to differentiate toward progenitors and propensity for symmetric expansion in our ex vivo assay. Remarkably, this differentiation latency and self-renewal behavior of FL-HSCs in vitro is consistent with recent studies that have tracked native FL-HSC fates in the FL niche in vivo.^3,9^

### Single cell RNA sequencing uncovers transcriptional heterogeneity in immunophenotypically defined FL-HSCs

In light of the functional heterogeneity of SE^hi^ FL-HSCs revealed by our single cell assays above, we next sought to explore the corresponding transcriptional heterogeneity of the immunophenotypic SE^hi^ FL-HSC population in the early-stage FL, which may provide insight into the gene regulatory programs responsible for observed differences in self-renewal and engraftment potential of cells in this immunophenotypically homogenous population. To this end, we sorted SE^hi^ cells from freshly isolated FL at E13.5 for scRNAseq (Figures 4A and S4A). After filtering out poor quality cells, we obtained genome-wide transcriptome data for 338 cells. Using the Monocle3 toolkit ^24^, we performed dimensionality reduction by uniform manifold approximation and projection (UMAP) and unsupervised clustering by the Louvain algorithm to identify distinct cell clusters (Figure 4B). E13.5 SE^hi^ cells globally express *Procr* (EPCR), as expected, as well as other genes associated with HSCs, such as *Hlf, Esam, Mecom*, *Fgd5*, *H19*, and *Cdkn1c,* and mostly lack expression of genes associated with differentiating progenitors and mature blood cells (Figures S4B and S4C). However, the expression levels of HSC-associated transcripts varied among cell clusters across UMAP space, suggesting underlying heterogeneity in the SE^hi^ population. Moreover, although this population is negative for CD48 by flow cytometry (Figure S1A), we observed heterogenous expression of *Cd48* transcript (Figure S4C). *Flt3* expression was also observed heterogeneously (Figure S4C), which may imply presence of previously described eMPP^9^ or developmentally-restricted HSCs^10^ (known to express *Flt3*) in the SE^hi^ population. Furthermore, while FL-HSCs are generally thought to be proliferative,^1,25^ we observed substantial variability in the expressions of genes associated with cell cycle activity within the immunophenotypically defined SE^hi^ FL-HSCs, including a distinct population with low expression of cell cycle genes and low proliferation index, a transcriptional measure of cell cycle activity (Figure S4D).

**Figure 4:**
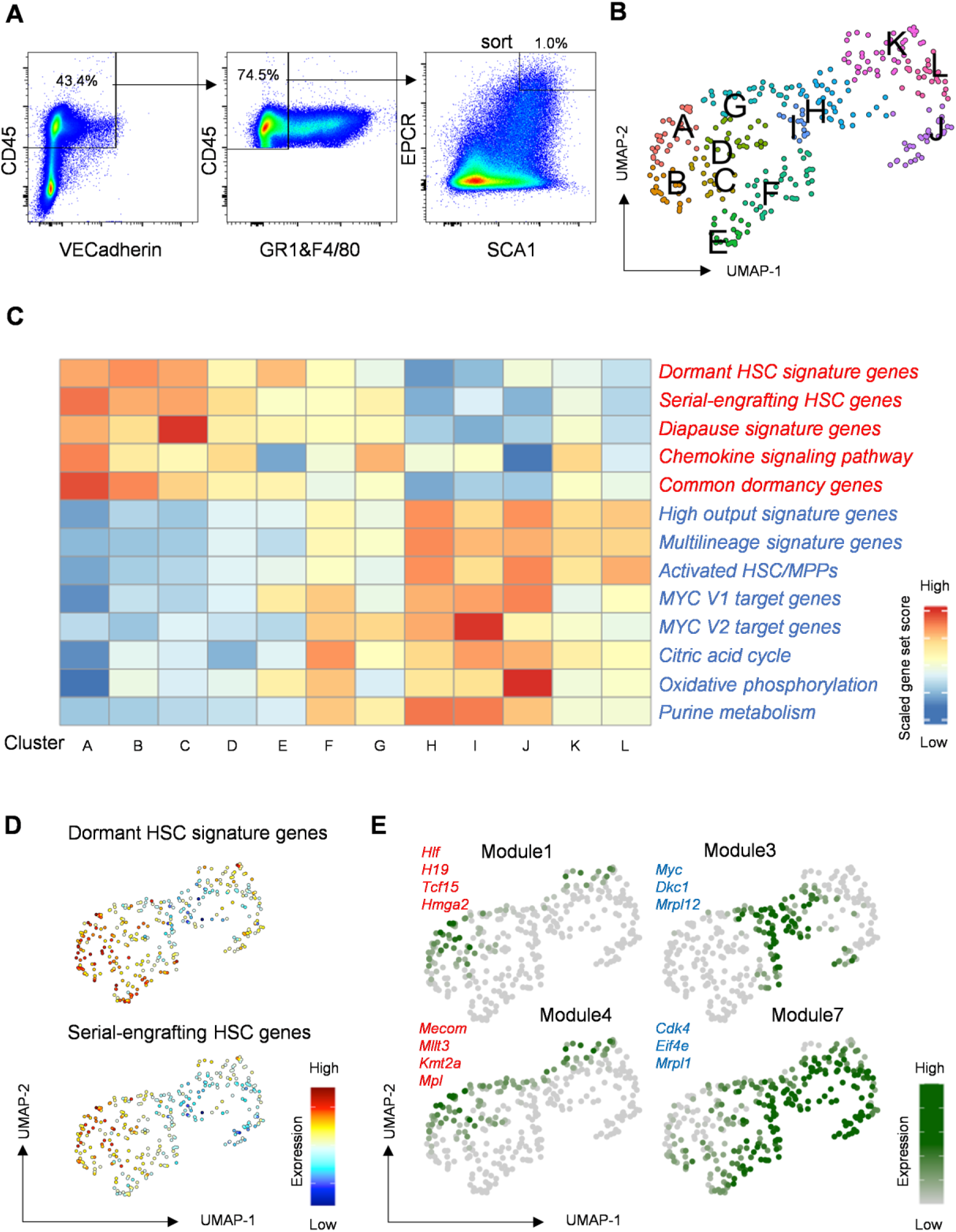
Single cell RNA sequencing reveals transcriptional heterogeneity amongst immunophenotypically-defined SE^hi^ FL-HSCs. **(A)** Sorting strategy for SE^hi^ FL-HSCs from freshly isolated E13.5 FL used for scRNAseq. **(B)** Unsupervised clustering of E13.5 SE^hi^ FL-HSC scRNAseq data in UMAP. **(C)** Heatmap of gene-set scores by cluster for genes associated with adult HSC dormancy,^27^ serial-engrafting HSCs,^28^ diapause,^29,31^ chemokine signaling (WP_CHEMOKINE_SIGNALING_PATHWAY), and a common stem cell dormancy state associated with lipid metabolism,^32^ or genes associated with activated HSC/MPP states including high output and multilineage signatures,^27,28,84^ Myc pathway activation (Hallmark Myc Target Genes V1, V2), and metabolic activity (WP_TCA_CYCLE, HALLMARK_OXIDATIVE_PHOSPHORYLATION, WP_PURINE_METABOLISM). **(D)** Gene-set expression heatmaps for dormant HSC signature genes and serial-engrafting HSC signature genes.^27,28^ **(E)** Expression heatmaps for modules of co-regulated genes determined using Louvain community analysis in Monocle3. Gene modules 1, 3, 4 and 7 are shown with representative genes identified in each module.

We next examined expression of gene-sets that characterize dormant and serial-engrafting HSCs in adult BM across cell clusters in the SE^hi^ FL-HSC scRNAseq data (Figure 4C, Table S1).^26–29^ We observed a marked variation in the expression levels of the adult HSC and dormancy-related gene-sets across the population of SE^hi^ cells, with clusters exhibiting the highest dormancy-associated gene-set scores predominantly localized to regions of UMAP with the lowest expression of genes associated with active cell cycle state and low proliferation index (Figures 4C, 4D and S4D). In contrast, genes characterizing activated HSCs/MPPs in adult BM and MYC target genes that promote biosynthetic and cell cycle activation were lower in this subset (Figure 4C, Table S1).^27,28,30^ Interestingly, the expression pattern of genes associated with diapause,^29,31^ a reversible embryonic state of dormancy unrelated to hematopoiesis, showed a similar pattern to the HSC dormancy gene-set, as did genes associated with chemokine signaling characteristic of dormant HSCs in adult BM^27^ and a gene signature associated with lipid metabolism and dormancy common to embryonic diapause and multiple adult stem cell types^32^ (Figure 4C, Table S1).

As an orthogonal validation, we carried out an analysis of the same dataset using a Bayesian approach of latent variable modeling (LVM) and feature extraction (MOFA-based framework, MuVI).^33–35^ Using the aforementioned gene signatures as the Bayesian priors in LVM, SE^hi^ FL-HSCs were embedded as two distinct islands of cells in latent space (Figure S5A). The isolated minor island (cluster 2, 5, 7) was composed of cells with strong association with serial-engrafting and dormant HSC gene-set features, but negatively correlated with oxidative phosphorylation and MYC-V1 target gene-set features (Figure S5B), supporting the heterogeneity of SE^hi^ FL-HSC observed in our prior analysis.

To further characterize transcriptional heterogeneity of the immunophenotypic population of FL-HSCs in an unbiased manner, we performed gene module analysis across cell clusters to determine co-regulated genes (Figures 4E, S4E and Table S2). We identified two modules (#1 and #4) of genes with cluster-specific expression patterns that are similar to dormant and serial-engrafting HSC and diapause gene-sets (compare Figure S4E and Figure 4C). Consistent with this observation, genes expressed in these modules include known HSC-specific genes (e.g. *Hlf, Mecom, Mllt3*) (Table S2). Gene Ontology (GO) analysis of genes in these two modules identified terms associated with dormancy in adult HSCs, including stem cell population maintenance, lysosomal transport, protein targeting to lysosome, and positive regulation of catabolic process.^25,36,37^ In contrast, GO analysis of genes in module 3 and module 7, which have non-overlapping expression patterns to modules 1 and 4, identified terms associated with activated metabolic and cell cycle states, such as DNA replication initiation,^38,39^ ATP biosynthetic process,^40,41^ oxidative phosphorylation,^42–44^ and translation^45,46^ (Table S2). Furthermore, gene-sets specifically related to the cellular metabolism of activated HSCs and MPPs^27,47,48^ exhibit similar cluster-specific expression patterns to gene modules 3 and 7 (compare Figure S4E and Figure 4C). To independently verify observed transcriptional heterogeneity in the early fetal liver HSC compartment, we also analyzed published scRNAseq data from E12.5 FL.^49^ Extracting HSCs as described in the original article (Figures S6A and S6B), we observed heterogeneity in proliferation index (Figure S6C), gene modules (Figures S6D-F and Table S3) and gene-set scores for HSC self-renewal and dormancy-related signatures (Figure S6G) analogous to that identified in our scRNAseq data. We also performed scRNAseq on HSCs sorted from human FL samples using stringent immunophenotypic markers (CD34^+^CD38^-^CD45RA^-^CD90^+^EPCR^+^) and observed heterogeneous clusters of human FL-HSCs characterized by differences in dormancy-related gene signatures and lipid metabolism, similar to that observed in murine FL-HSCs (Figure S7, Table S4).

Collectively, we here observed that immunophenotypically defined FL-HSCs are transcriptionally heterogeneous in their biosynthetic and cell cycle activation states, and we identified a subset exhibiting gene signatures overlapping with dormant adult HSCs, embryonic diapause^29,31^ and other dormant stem cell states,^32^ particularly those associated with lipid metabolism and lysosomal activity. These results are consistent with our observation that the same immunophenotypic SE^hi^ FL-HSCs exhibit heterogeneous self-renewal, proliferation, differentiation, and engraftment properties revealed by single cell coculture in the FL-AKT-EC niche. Furthermore, the distinct in vitro behavior of the subset of SE^hi^ FL-HSCs with serial engraftment potential (lower proliferation, latency to differentiate, and relatively homogenous ESAM^+^SCA1^+^EPCR^+^ HSC immunophenotype) suggests that this subset corresponds to cells in our scRNAseq data characterized by a transcriptional state of relative biosynthetic dormancy and decreased proliferation index.

### Integrating single cell transcriptomics and transplantation assays from clonal progeny of FL-HSCs identifies transcriptional signatures of self-renewing HSCs with serial multilineage engraftment

Based on our single cell transcriptomic analysis of freshly isolated SE^hi^ FL-HSCs above, we hypothesized that the subset of HSC-like colonies arising from SE^hi^ FL-HSCs possessing serial engraftment potential could also be distinguished by transcriptional signatures of biosynthetic dormancy. To test this hypothesis, we integrated scRNAseq into our ex vivo platform for clonal analysis of FL-HSCs (Figure 5A). Briefly, we index sorted single E13.5 FL SE^hi^ cells into a 96-well plate containing FL-AKT-EC. Following coculture, a portion of each colony (15%) was used for immunophenotyping by flow cytometry to identify HSC-like colonies for further analysis (Figure S8A). A portion of the remaining cells from each of these colonies was used for transplantation (10% per recipient, to three recipients per colony) and for scRNAseq (55%). To minimize batch effects between colonies, we individually labelled each colony with oligonucleotide-barcode-conjugated antibodies to ubiquitous murine cell surface antigens such that cells from each colony could be pooled for downstream steps of scRNAseq.^50^ This approach allows us to correlate the functional engraftment properties of each HSC-like colony to the unique transcriptomes of cells in the colony, as well as to the prospectively collected immunophenotype of the individually sorted E13.5 SE^hi^ FL-HSCs from which each HSC-like colony was derived.

**Figure 5:**
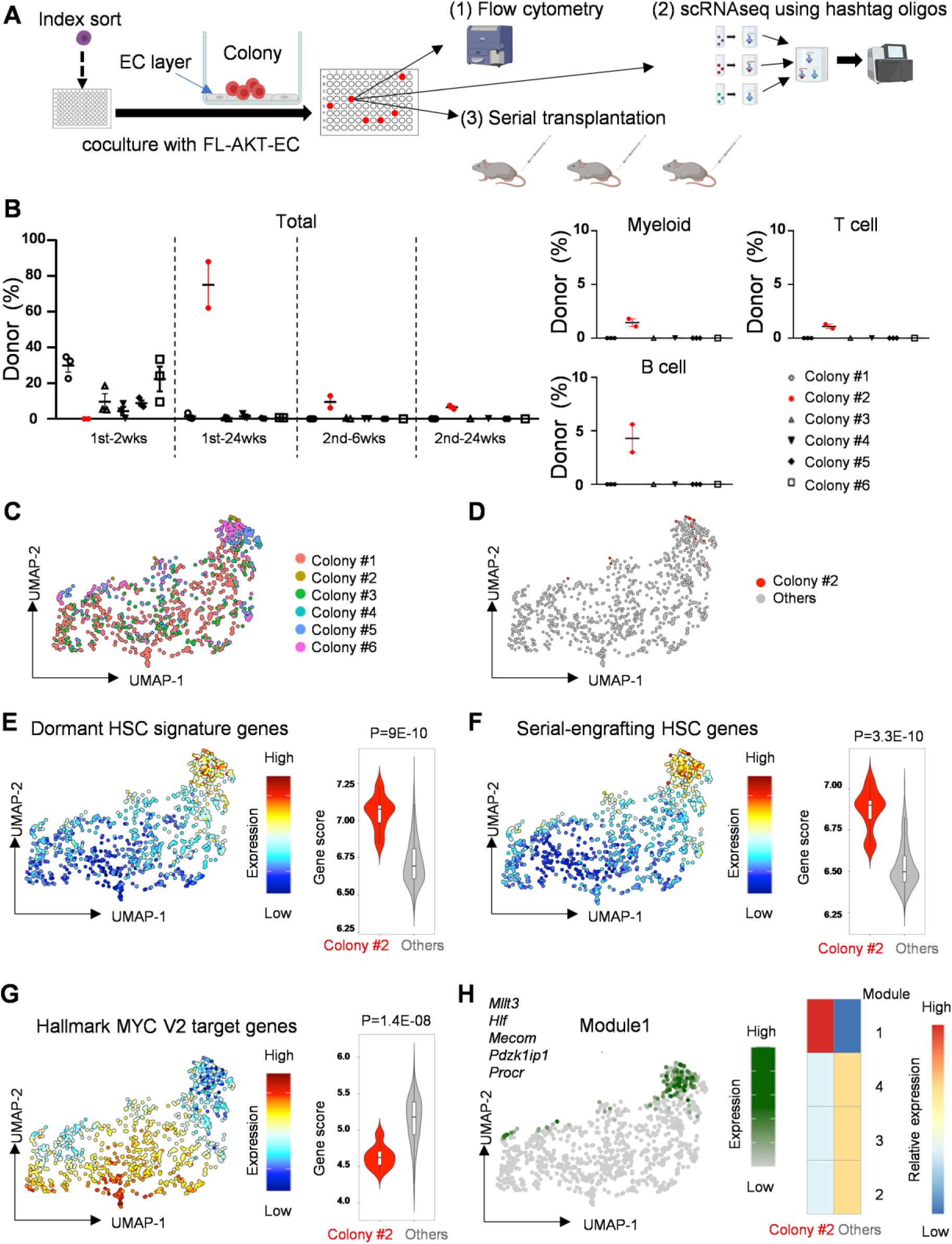
Analysis of clonal progeny of single FL-HSCs by integrating scRNAseq, flow cytometry, and transplantation reveals transcriptional and functional heterogeneity of HSC colonies. **(A)** Overview of the experiment design. 15% of each well was used for flow cytometric analysis, 55% for scRNAseq, and 30% for serial transplantation (10% to each of 3 recipient mice). **(B)** Donor chimerism in peripheral blood of primary and secondary recipients transplanted with the progeny of single E13.5 SE^hi^ cells following coculture with FL-AKT-ECs. Left: Total donor chimerism at indicated timepoints following primary and secondary transplant. Right: Donor myeloid, B cell, and T cell contribution at 24 weeks in secondary recipients. (n=3 mice transplanted for each colony). **(C)** UMAP analysis with cells labeled based on colony of origin (colony #1: 513 cells, colony #2: 16 cells, colony #3: 202 cells, colony #4: 44 cells, colony #5: 102 cells, colony #6: 153 cells). **(D)** UMAP of cells grouped by engraftment properties (serially engrafting colony #2 verses all other colonies lacking serial engraftment). **(E)** Heatmap of gene-set scores characterizing dormant adult HSCs.^27^ Violin plots of gene-set scores by colony type. p values indicate Wilcoxon rank-sum test. **(F)** Heatmap of gene-set scores characterizing serial-engrafting HSCs.^28^ Violin plots of gene-set scores by colony type. p values indicate Wilcoxon rank-sum test. **(G)** Heatmap of gene-set scores for Myc target genes (HALLMARK_MYC_TARGETS_V2). Violin plots of gene-set scores by colony type. p values indicate Wilcoxon rank-sum test. **(H)** Expression heatmaps for modules of co-regulated genes determined using Louvain community analysis. Gene modules 1 is shown in UMAP with representative genes (left). Heatmap of expression of gene modules by colony type (right).

Using this approach, we found that most identified HSC-like colonies contributed to short-term engraftment whereas colony #2 uniquely contained long-term, multilineage engrafting HSCs by serial transplantation (Figure 5B and S8B). After filtering for poor quality cells, we obtained scRNAseq data for 1,030 cells, which were distinguished by colony of origin after dimensionality reduction by UMAP using the combined scRNAseq data (Figure 5C). Expression of genes associated with HSC state, including *Hlf, Procr, and Mecom*, varied significantly in UMAP space, as did expression of *Cd48* and other genes associated with differentiation of HSCs to multipotent progenitors (Figure S8C). Cells from all HSC-like colonies generally lacked expression of genes associated with more lineage-restricted progenitors and mature blood cells, though a small population of cells expressing markers of granulocyte differentiation (*Elane*) was observed in cells coming primarily from colony #1 (Figures S8C and 5C), which was notably more proliferative than other HSC-like colonies and contained a larger proportion of cells lacking SCA1^+^EPCR^+^ co-expression (Figure S8A).

Interestingly, the majority of cells from colony #2 clustered together in regions of UMAP space with the highest expression of HSC-associated genes and absence of genes associated with differentiated progenitors (Figure 5D, S8C). Furthermore, when we examined published adult HSC-specific gene signatures associated with dormancy and serial engraftment,^26–28^ we found expression to be significantly higher in cells from colony #2 compared to cells from other types of colonies (Figures 5E, 5F and S8D). Interestingly, cells in colony #2 also exhibited significantly higher expression of genes associated with long-term, multilineage engraftment following in vitro expansion of adult HSCs,^13^ diapause,^29,31^ lipid metabolism and dormancy common to embryonic diapause and multiple adult stem cell types,^32^ and chemokine signaling (Figure S8D). In contrast, MYC target genes that promote biosynthetic and cell cycle activation, as well as gene-sets characterizing differentiated progenitors, were significantly lower in cells from colony #2 (Figures 5G and S8E). Differences in biosynthetic signatures were also confirmed by the comparison of cellular metabolism gene-sets which are reportedly higher in activated HSC/MPPs^27,47,48^ (Figure S9A). When performing LVM on cells from these colonies using metabolism-related gene-sets as Bayesian priors, colony #2 stood out from the rest of the colonies in latent embedding as shown on the factor loading dot plot (Figure S10A). Specifically, colony #2 showed a shift away from transcriptomic signature of carbohydrate metabolism and towards that of lipid metabolism (Figures S10A and S10B).

To further characterize the transcriptional heterogeneity of HSC-like colonies in an unbiased manner, we performed unsupervised clustering (Figure S9B) and gene module analysis across cell clusters to determine co-regulated genes (Figures 5H and S9C). One of the identified gene modules (module 1) which specifically marks cluster 1 (containing the majority of cells from colony #2) includes known HSC-specific genes such as *Hlf*, *Mecom*, and *Mllt3*, and genes related to autophagy, catabolic processes, epigenetic regulation, and immune/interferon responses based on Gene Ontology (GO) analysis (Table S5). In contrast, non-overlapping gene modules feature active cell cycle genes and genes related to biosynthetic activity (module 2-4), including ribosome biogenesis, translation, mitochondrial activity, and ATP synthesis based on GO analysis (Table S5).

Next, to identify genes associated with serial engraftment properties, we determined differentially expressed genes between cells in colony #2 and cells from other HSC colonies lacking serial engraftment (Table S6). Cross-referencing these genes with genes expressed in module 1 (Table S5), we identified 123 common genes representing a gene-set that we termed “serially engrafting FL-HSC genes” (Table S1). Remarkably 40% of the genes identified in our novel gene-set were shared with those in published gene-sets either associated with dormant adult BM HSCs or serially engrafting adult HSCs,^27,28^ and these gene-sets marked similar subsets of cells in both our cultured and freshly isolated FL-HSC scRNAseq data (Figures S9D and S9E). Common genes identified include a number of key regulators of adult HSC dormancy and self-renewal, including *Angpt1*, *Mecom*, *Mllt3, Cd63,* and *Ndn.*^3,51–56^ Interestingly, the *Cxcr4*-expressing subset of hemogenic endothelium (HE) in the E9-9.5 mouse embryo, which we recently demonstrated to contain the majority of HSC-competent HE,^4^ also showed significantly higher expression of this gene-set compared with HE lacking *Cxcr4* expression, which can give rise to MPP but largely lack HSC potential (Figure S9F).

When we further limited differential expression analysis to cluster #1 (which is appears to be enriched in HSC-specific genes), we identified significantly increased expression of HSC-associated genes including *Mllt3*^54^ and *Apoe,*^57^ as well as increased expression of gene-sets associated with dormancy and serial engraftment, in cells from colony #2 compared with cells from other colonies (Figure S9G, Table S6). These findings suggest potentially key roles for those genes in maintenance of HSC self-renewal in the FL-AKT-EC niche. Notably, *Esam* expression was also significantly enriched in colony #2 cells compared to cells from other colonies in cluster 1 (Figure S9H), consistent with our finding that ESAM expression in HSC-like colonies predicts serial engraftment potential.

In a complementary experiment, we compared ESAM^+^ HSC colonies to colonies lacking significant ESAM expression by scRNAseq, to determine whether functional differences in engraftment observed between these colony types correlate with differences in their transcriptional properties (Figure S11A). We observed that cells from ESAM^+^ HSC colonies (1-5) largely localized in UMAP space separately from cells from other colonies (6-7), and that cells from ESAM^+^ HSC colonies were enriched in self-renewal and dormancy-associated gene signatures (Figure S11B-H). Furthermore, gene module analysis demonstrated enriched expression of HSC-associated genes such as *Apoe, Hlf, Mllt3, Cd63,* and *Mecom* in regions of UMAP space associated with ESAM^+^ HSC colonies (Figure S11I, Table S7). These results are consistent with the transcriptional differences we observed between HSC-like colonies that differ based on their direct engraftment potential (Figure 5).

Taken together, these studies reveal that serially engrafting (ESAM^+^) HSC colonies are characterized by gene expression patterns associated with dormancy in adult BM HSCs and other stem cells, including lower biosynthetic/MYC pathway activation, a shift toward lipid metabolism, and decreased cell cycle activity.^27^ The finding that this gene expression profile also characterizes a subset of freshly isolated E13.5 SE^hi^ FL-HSCs as well as HSC-competent HE in the early embryo^4^ suggests a conserved gene expression program of biosynthetic dormancy established in early embryonic development may be essential for initiating and maintaining HSC fate.

### Single cell transcriptomic analysis identifies candidate signaling interactions supporting FL-HSC self-renewal in the FL-AKT-EC niche

FL-AKT-ECs provide a niche sufficient to support expansion of FL-HSCs, revealing that the subset of immunophenotypic SE^hi^ FL-HSCs with serial engraftment potential are intrinsically primed for differentiation latency and symmetric self-renewal behavior. To elucidate potential extrinsic signaling interactions regulating the process of FL-HSC self-renewal, we first integrated scRNAseq data from FL-AKT-ECs and serially engrafting HSCs (from HSC colony #2, Figure 5B), and broadly identified candidate ligand-receptor interactions using a comprehensive database of curated pairs of ligands and receptors^58,59^ (Table S8). Since FL-ATK-EC-conditioned media was insufficient to support the generation of ESAM^+^ HSC colonies (Figure S12), suggesting that direct contact with FL-AKT-ECs is indispensable for FL-HSC self-renewal, we included both soluble and transmembrane ligands expressed by FL-AKT-EC. This unbiased analysis identified an extensive list of potential signaling interactions, many of which have been previously shown to regulate various aspects of HSC self-renewal in the context of embryonic development and/or the adult BM, including those involving Notch, TGF-β, Wnt/β-catenin, integrins, and cytokines/chemokines (SCF, CXCL12, IGF1).

To further define the pathways that may be specifically required for HSC self-renewal in the FL-AKT-EC niche, we applied NicheNet,^60^ a computational algorithm that identifies ligand-receptor pairs from scRNAseq data based on knowledge of downstream signaling pathways and gene regulatory networks. Integrating scRNAseq data from FL-AKT-EC and HSC-like colonies (Figure 6A), NicheNet identified candidate ligand-receptor signaling interactions prioritized based on likelihood of accounting for genes significantly enriched in serially engrafting FL-HSCs (from colony #2) compared with cells from other HSC-like colonies in our scRNAseq data (Figures 6B-D). Multiple ligand-receptor pairs and downstream pathways were identified, including *tgfb1* (TGFβ1), a well-established regulator of HSC quiescence in adult BM,^61,62^ transmembrane and extracellular cell adhesion molecules (egs. *Icam1, Col4a1, Lamb2*) that bind and activate integrin receptors (*Itgb1, Itgav*) essential for colonization of HSCs to the FL^63^ and for regulating the size of the FL-HSC pool^64^, *Selp* (P-selectin), implicated in HSC expansion in the zebrafish caudal hematopoietic territory (CHT, equivalent to the mammalian FL),^65^ and *Jam3*, encoding a cell surface adhesion protein implicated in HSC maintenance in the adult BM.^66^ Numerous ligand-receptor signals not previously implicated in HSC self-renewal or fate determination were also identified. Notably, a large proportion of identified interactions were predicted to contribute to regulating downstream genes that are essential to HSC dormancy and self-renewal, particularly *Mecom*, *Prdm16*, and *Angpt1*.^3,51,52,67^ To further validate that our findings in the ex vivo FL-AKT-EC niche reflect signals endogenous to FL endothelial cells in vivo, we repeated this ligand-receptor analysis using published scRNAseq data from E14 FL-ECs,^68^ demonstrating substantial overlap in identified signaling interactions (Figure S13, Table S9). These results suggest that combinatorial ligand-receptor interactions between niche FL-ECs and FL-HSCs involving numerous, integrated signaling pathways are likely essential to the self-renewal of serially engrafting HSCs in vitro (Graphical abstract). Altogether, our ex vivo clonal FL-HSC expansion platform and correlative single cell transcriptomic data provide a valuable resource for future studies to determine how these complex receptor-ligand interactions in the FL vascular niche coordinately regulate downstream transcriptional programs essential to FL-HSC self-renewal.

**Figure 6:**
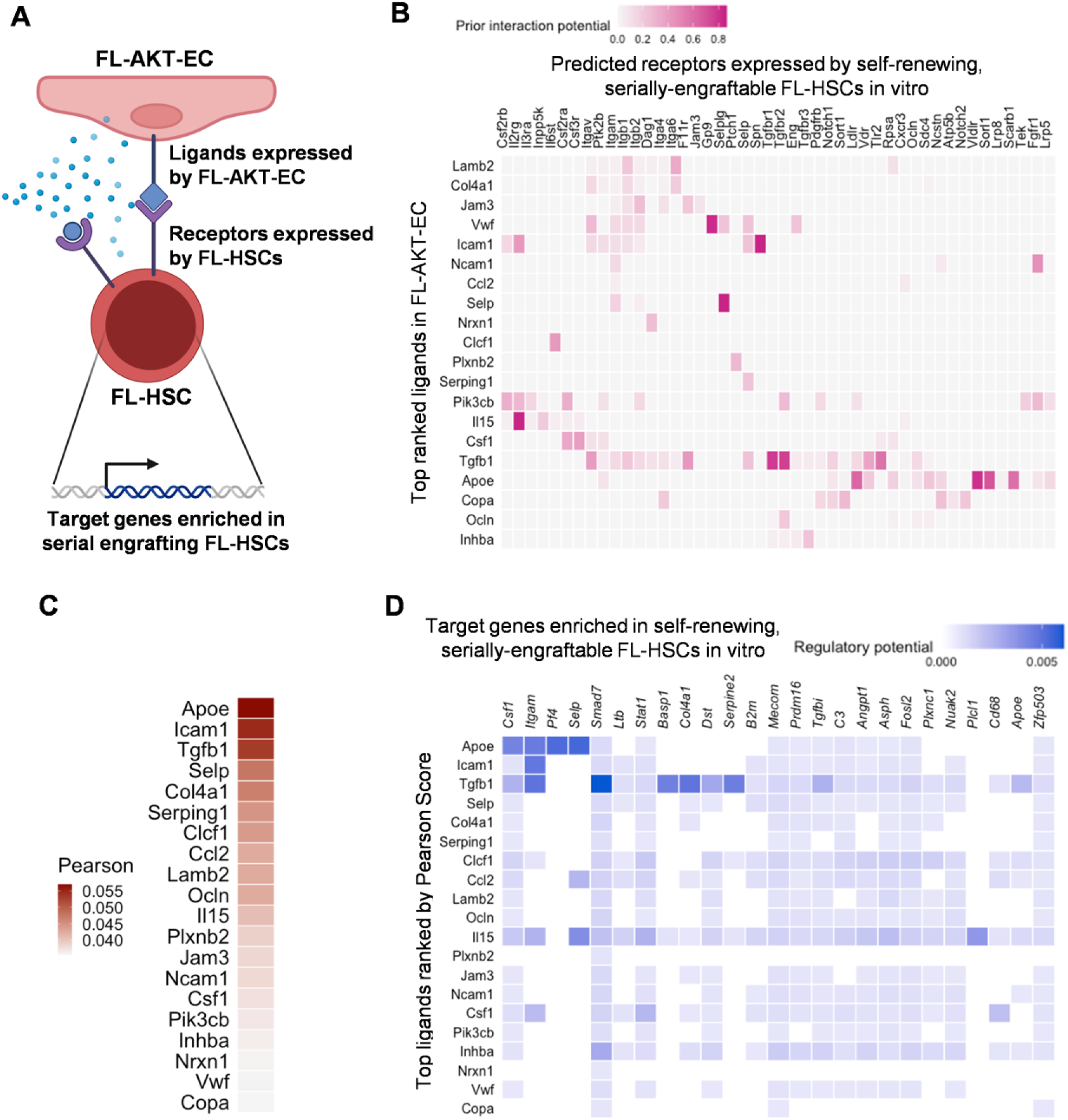
Complementary analysis of scRNAseq data identifies candidate receptor-ligand interactions regulating self-renewal of serially engrafting FL-HSCs in the FL-AKT-EC niche. **(A)** Model for analysis of receptor-ligand interactions by NicheNet.^76^ **(B)** Heatmap showing candidate ligands (expressed by FL-AKT-EC) interacting with receptors (expressed by cells in serially engrafting HSC colony #2) inferred by NicheNet. **(C)** Heatmap showing candidate ligands (expressed by FL-AKT-EC) ranked by Pearson correlation. **(D)** Heatmap showing the regulatory potential between the top ranked ligands (expressed by FL-AKT-EC) and downstream genes whose expression is enriched in cells from serially engrafting HSC colony (#2) compared to cells from other HSC-like colonies lacking serial engraftment.

### Live imaging in the FL-AKT-EC niche reveals FL-HSC undergo predominantly symmetric cell divisions

Adult murine and human HSCs can undergo asymmetric cell divisions in which the paired daughter cells inherit differential activation states, with the more dormant cell maintaining HSC fate necessary for self-renewal while the activated cell adopts a progenitor state necessary to actively contribute to hematopoiesis.^36,69^ Our immunophenotypic, functional, and transcriptomic analysis of FL-HSCs supports a distinct model in which serially engrafting HSCs predominantly undergo symmetric self-renewal and homogenous expansion in the FL-EC niche, giving rise to daughter HSCs that maintain a relatively dormant biosynthetic state and delayed differentiation to progenitors. Based on this model, we predict that FL-HSCs should be distinguishable based on symmetric cell division behavior observed during coculture with FL-AKT-EC in vitro. To test this prediction, we index sorted single SE^hi^ FL cells (co-stained with CellTrace-Far Red) to FL-AKT-EC and observed their initial cell divisions by live imaging to measure relative fluorescence intensity (total protein content) of daughter cells as a marker of division symmetry (Figure 7A). Consistent with our hypothesis, we found that ESAM^+^ HSC colonies were derived from SE^hi^ FL cells observed to almost exclusively undergo symmetric cell division (ratio of CellTrace fluorescence intensity between daughter cells ∼1) whereas other colony types were derived from cells that underwent a broader distribution of symmetric and asymmetric divisions (ratio of CellTrace fluorescence intensity between daughter cells of <1) (Figure 7B-C).

**Figure 7:**
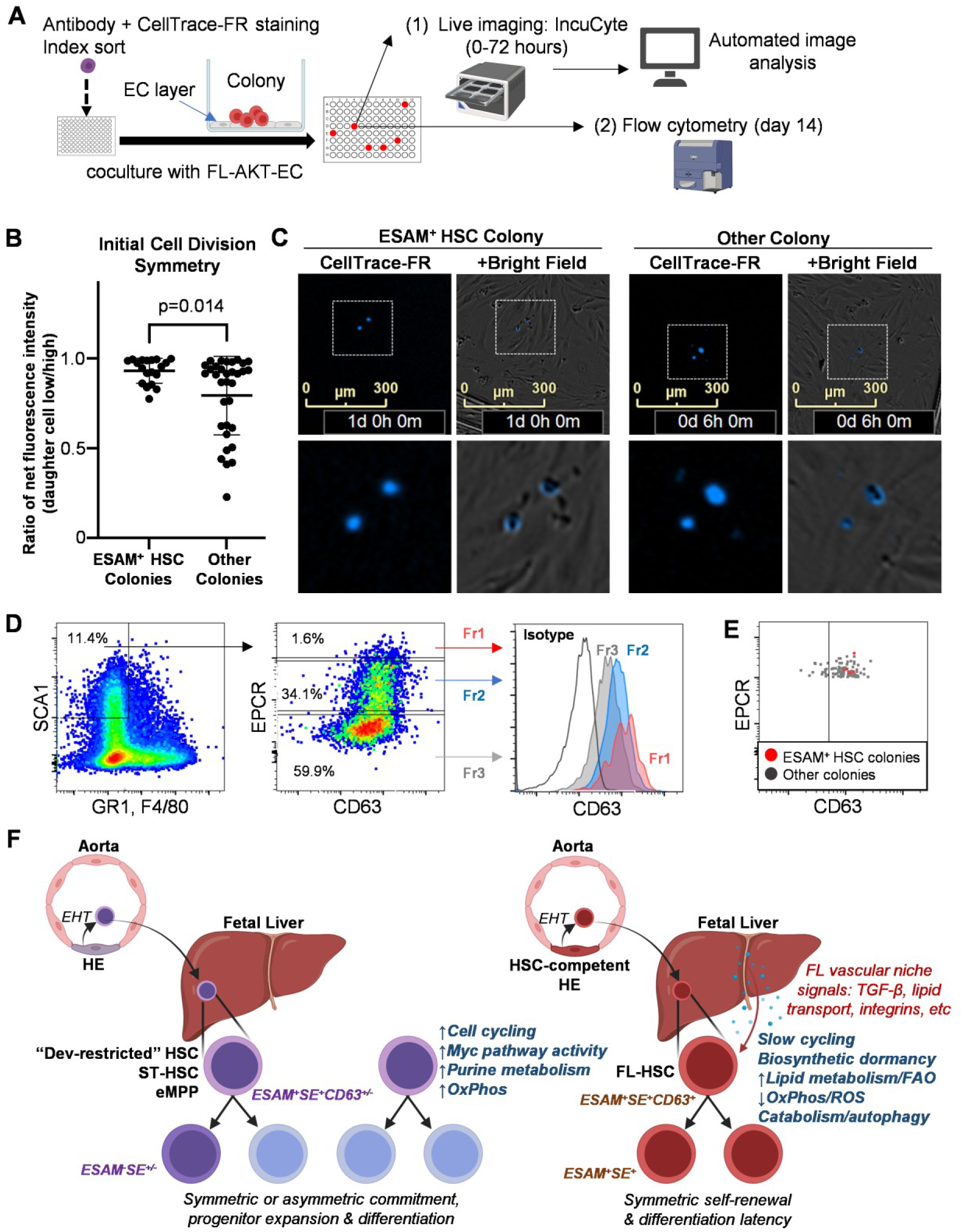
Live imaging reveals FL-HSCs are characterized by symmetric cell divisions. **(A)** Overview of the experiment design. E15.5/15.6 SE^hi^ FL cells co-stained with CellTrace-FR were individually index sorted to FL-AKT-EC and cell divisions monitored by live imaging for initial 48 hours of coculture. At day 14, colonies were harvested and analyzed by flow cytometry. **(B)** Cell division symmetry measured by ratio of fluorescence intensity of CellTrace-FR inherited by daughter cells amongst SE^hi^ FL cells after first cell division that either generated ESAM^+^ HSC colonies or other colony types (n = 118 colonies pooled from two independent experiments). Error bars represent standard deviation. **(C)** CellTrace-FR and brightfield images of initial cell division in a representative ESAM^+^ HSC colony (symmetric division) and other colony type (asymmetric division). **(D)** Relative expression of CD63 in E13.5 FL CD45^+^GR1^-^F4/80^-^SCA1^+^ HSPCs divided by EPCR expression. Fr1: EPCR high (red), Fr2: EPCR medium (blue), Fr3: EPCR low (gray). Fr: Fraction. **(E)** Expression of CD63 by index analysis of E13.5 SE^hi^ FL-HSCs giving rise to immunophenotypic ESAM^+^ HSC colonies (red) or other colony types (gray) following FL-AKT-EC coculture. **(F)** Summary model. The FL SE^hi^ compartment comprises a heterogeneous population of HSC-independent hematopoietic descendants (including embryonic multipotent progenitors, eMPP,^9^ short-term and developmentally-restricted HSCs^10^, which may actively contribute to prenatal hematopoiesis) and functionally competent (serially transplantable) FL-HSCs, which are preserved for adult hematopoiesis. FL-HSCs exhibit distinctly dormant biosynthetic and metabolic states, symmetric self-renewal behavior, reduced cell cycle kinetics, and differentiation latency, facilitating the regulated expansion of the HSC pool in the context of supportive signals from the FL vascular niche.

Lysosome-associated membrane protein 3 (Lamp3, or CD63), a tetraspanin with roles in lysosomal trafficking, cholesterol transport, and extracellular vesicle (EV) formation, was found to be highly enriched in serially engrafting FL-HSCs based on our transcriptomic analysis. CD63 has been implicated in HSC cell division symmetry^70^ and is functionally involved in HSC self-renewal and dormancy through interactions with the TGF-β pathway.^28,55^ A recent study has also suggested a role for CD63 in HSC self-renewal via autocrine signals mediated by HSC-derived EVs downstream of NADPH-driven cholesterol biosynthesis.^71^ Consistent with observations that CD63 is expressed on dormant adult HSCs with enhanced reconstitution potential, we observed that CD63 expression was enriched in primary E13.5 FL-HSPCs expressing the highest levels of EPCR (Figures 7D and S14A) and that ESAM^+^ HSC colonies were primarily derived from the CD63^+^ fraction of SE^hi^ FL-HSCs based on index flow cytometry analysis (Figure 7E and S14B). Together, these results suggest that SE^hi^ FL-HSCs with serial engraftment potential are characterized by increased CD63 expression and by their specific propensity for symmetric cell division. Collectively, our findings support a paradigm in which FL-HSCs are maintained in a relatively biosynthetically dormant, slow cycling state characterized by symmetric self-renewal and differentiation latency, which is supported by signals from the FL vascular niche (Figure 7F). Critically, this stage-specific behavior may function to expand the HSC pool during fetal development while delaying their contribution to active hematopoiesis until after birth, when HSCs have transitioned from the FL and to the bone marrow niche.

## Discussion

Here we establish a novel ex vivo platform that supports the expansion of serially engrafting HSCs from individually sorted FL-HSCs. This platform provides a powerful tool to analyze the intrinsic properties of functionally validated FL-HSCs at single cell resolution, as well as the extrinsic niche-derived signals enabling HSC self-renewal in vitro. To this end, we leveraged this platform to resolve heterogeneity in immunophenotypically defined FL-HSCs by simultaneously integrating index flow cytometry, live imaging, single cell transcriptomics, and transplantation assays. In so doing, we identified a rare subset of FL-HSCs with serial engraftment potential that are characterized by propensity for symmetric cell divisions, differentiation latency, and transcriptional signatures of lipid metabolism and biosynthetic dormancy.

Notably, the observed behavior and transcriptional profiles of serially engrafting FL-HSCs in our study contrast with previous paradigms proposing that FL-HSCs reside in a state of heightened biosynthetic and proliferative activity necessary to simultaneously expand their total numbers (providing sufficient HSCs for the adult) and differentiate to progenitors (providing functional blood cells for fetal development). Instead, our findings support a shift in this paradigm in line with several recent studies that have traced the origin and evolution of HSCs and HSC-independent progenitors from early embryonic development to adulthood by various clonal assays and lineage tracing methodologies in vitro and in vivo.^3–5,9,72^ Together, these studies suggest that functional, long-term engrafting HSCs and HSC-independent embryonic multipotent progenitors (eMPP) emerge from distinct populations of HE, that HSCs contribute minimally to differentiating progenitors and mature blood cells prenatally, and that eMPP contribute prenatally as well as postnatally to lifelong multilineage hematopoiesis natively in vivo but fail to provide long-term, multilineage hematopoietic engraftment as measured by transplantation assays. Remarkably, our platform revealed intrinsic behaviors of individual FL-HSCs in vitro that recapitulated their predicted behaviors consistent with these recent studies in vivo.^3,9,72^ Specifically, the rare subset of immunophenotypic (SE^hi^) FL-HSCs at early and mid-gestation with serial engraftment potential gave rise to colonies that consisted of fewer total cell numbers, homogenously expressed immunophenotypic markers of HSCs (i.e. EPCR, SCA1, ESAM), and supported serial engraftment in multiple mice transplanted from a single colony, suggesting symmetric self-renewal and constrained proliferative activity in vitro. Moreover, serially engrafting FL-HSC colonies demonstrated differentiation latency in vitro, failing to generate differentiated progenitors or mature blood cells based on immunophenotypic and transcriptomic analysis after extended coculture (12-15 days), which would temporally correspond to the postnatal period in vivo. Together, our data suggest the immunophenotypic (SE^hi^) FL-HSC compartment is heterogeneous, with only a minor subset representing functional, serially engrafting HSCs (Figure 7F). Based on their in vitro behavior and clonal engraftment properties, we postulate that the remaining cells in this compartment may include a spectrum of separately emerging, HSC-independent lineages with multilineage hematopoietic potential but with minimal or restricted (short-term or lineage biased) engraftment potential, including recently described eMPP (that lack engraftment potential based on transplantation assays)^9^ and “developmentally restricted” HSCs (that reconstitute multilineage hematopoiesis upon transplantation but with lymphoid biased engraftment).^10^

By integrating single cell transcriptomics into our platform, we also determined the unique transcriptional properties of serially engrafting FL-HSCs during the process of self-renewal in the ex vivo FL-AKT-EC niche. Surprisingly, this revealed a transcriptional signature of relative biosynthetic dormancy that overlaps with signatures of highly dormant and serially engrafting HSCs in the adult BM,^8^ as well as HSC-competent HE in the early embryo.^4^ Together, these studies support a novel paradigm in which HSC fate across prenatal development is linked to a state of relative dormancy that enables limited symmetric stem cell self-renewal while preventing excessive proliferation and differentiation to multilineage blood cells (Figure 7F). Crucially, this mechanism would serve to minimize metabolic, proteotoxic, and genotoxic stress during HSC development that could contribute to hematopoietic dysfunction later in adult life. Furthermore, these cell intrinsic dormancy-related programs may synergize with extrinsic mechanisms in the FL, for example niche-derived antioxidant systems that minimize oxidative stress^73^ and exogenous bile acids that alleviate endoplasmic reticulum stress,^74^ to protect self-renewing HSCs during their regulated expansion in the FL.

Studies of HSC development in the FL niche in vivo have been challenging due to the relative scarcity of functional, long-term engrafting HSCs amongst a sea of differentiating progenitors in the FL and the lack of markers that can prospectively identify FL-HSCs at single cell resolution with high specificity. By providing a culture system to functionally validate and characterize single FL-HSCs in vitro, our platform offers a complementary approach that provides several advantages and overcomes many challenges to studying FL-HSCs in vivo. We defined a combination of immunophenotypic markers in vitro (SCA1, EPCR, ESAM) sufficient to predict FL-HSCs expanded in vitro with serial engraftment potential in vivo, in line with previous studies that have identified these surface proteins as markers of HSC potential following in vitro culture.^13,22,75^ Our platform thus provides a means to screen more rapidly for functional HSC potential at the single cell level while reducing the need for resource-intensive transplantation in mouse models, which remains the gold standard to validate the engraftment potential of HSCs. Incorporating single cell index sorting into the platform, we also provide a means to retrospectively screen for immunophenotypic markers or other properties that can be captured by FACS (such as measures of metabolic activity, ROS, etc) which may predict engraftment potential. In the present study, for example, we demonstrated that functional FL-HSCs from the primary FL at E13.5 are enriched by CD63 surface expression, consistent with previous studies in adult HSCs.^55^ Our platform also facilitates direct visualization of HSCs actively self-renewing in a developmentally relevant ex vivo niche, enabling longitudinal observations of their behavior and interactions with niche cells, which is not currently possible in vivo due to technical limitations and lack of markers that can reliably identify functional FL-HSCs at single cell resolution in situ. Here we leveraged this capability, by further incorporating live fluorescence probes, to measure cell division symmetry in paired daughter cells, demonstrating that FL-HSCs are distinguished by exclusive symmetric cell division behaviors. Building on this work, our current efforts are utilizing machine learning approaches to determine whether additional aspects of cell morphology, motility, or other properties captured during live imaging in this platform can predict HSC fate, which may provide unprecedented insight into the distinct behaviors of FL-HSCs and their interaction with niche ECs.

Since the FL-AKT-EC niche was sufficient to support the in vitro self-renewal of serial engrafting FL-HSCs, we also leveraged the platform to identify potential niche signaling interactions that regulate the process of FL-HSC self-renewal, using complementary scRNAseq data from the FL-AKT-EC and FL-HSCs expanded in vitro. To this end, NicheNet^76^, a scRNAseq package which can infer and prioritize receptor-ligand pairs based on downstream gene expression changes in receiving cells, enabled identification of the signaling interactions between self-renewing HSCs and our engineered vascular environment that could account for genes identified as significantly enriched in engrafting HSC colonies verses immunophenotypically similar HSC-like colonies lacking long-term engraftment. Interestingly, TGFβ1, a key signaling molecule regulating dormancy in adult BM HSCs,^61,62,77^ was identified as one of the top 3 ligands in this analysis, with measurable regulatory potential for a large portion of differentially expressed genes in self-renewing HSCs in vitro (Figures 6C and 6D). This finding is consistent with the observation that serial engrafting FL-HSCs in our study were specifically enriched in transcriptional signatures associated with biosynthetic dormancy that are shared with adult BM HSCs. Thus, it is tempting to speculate a role for TGFβ signaling in the FL by regulating a relative state of dormancy in FL-HSCs. However, NicheNet also predicted signals involving multiple pathways that interact with TGFβ, including both positive and negative regulators and Smad7 (which can act via negative feedback on TGFβ/Activin pathways), suggesting fine tuning of TGFβ-mediated signals may be essential to achieve the appropriate balance of dormancy, allowing for FL-HSC expansion through controlled, symmetric self-renewal, while inhibiting excessive proliferation and differentiation. In addition to regulators of the TGFβ pathway, other signaling interactions and pathways identified by NicheNet have also been specifically implicated in HSC maintenance and self-renewal, including several intercellular and ECM adhesive interactions involving integrins, P-selectin, and JAM-C.^64–66^ Interestingly, the top ligand identified by NicheNet analysis, ApoE, has been postulated to regulate HSC quiescence via modulation of lipid/cholesterol efflux and oxidative stress.^78–80^ Indeed, mitochondrial fatty acid oxidation (FAO) is essential for adult HSC self-renewal^81,82^ and ex vivo maintenance of engrafting HSCs has been shown to be dependent on exogenous cholesterol/fatty acids that may serve as a source of eicosanoid signaling molecules and other lipid-derivatives essential to HSC function.^83^ Moreover, a recent study showed that FAO-generated NADPH fuels cholesterol biosynthesis in HSCs, which is essential for HSC self-renewal through both cell autonomous mechanisms and through HSC-niche interactions mediated by extracellular vesicles.^71^ Altogether, these results predict that complex, combinatorial interactions involving both EC niche-derived signals and cell intrinsic properties of FL-HSCs synergistically contribute to regulating the relative biosynthetic dormancy and cell cycle activity required for HSC self-renewal (Graphical abstract).

In summary, using a novel in vitro assay to assess for functional FL-HSCs at single cell resolution, we report here previously unrecognized heterogeneity of immunophenotypic FL-HSCs, revealing a subset of serially engraftable FL-HSCs characterized by intrinsic propensity for symmetric self-renewal, differentiation latency, and biosynthetic dormancy that is concomitantly dependent on critical extrinsic signals from the FL vascular niche. These findings support a paradigm shift in line with other recent studies that suggest prenatal hematopoiesis is largely supplied by HSC-independent progenitors whereas FL-HSCs are preserved to undergo limited self-renewal without differentiation during development to generate the adult HSC pool. Interestingly, we show that the minor subset of serially transplantable FL-HSCs exhibit transcriptional profiles overlapping with those of dormant and serially-engrafting adult HSCs, including biosynthetic dormancy, lipid metabolism/FAO, lysosomal activity, and autophagy. However, adult HSCs are largely quiescent (cell cycle inactive) at homeostasis and, when activated by physiologic stress, can undergo asymmetric cell divisions to simultaneously self-renew and contribute progenitors to active hematopoiesis. Our studies demonstrate that FL-HSCs, in contrast, predominantly undergo symmetric self-renewal without progenitor differentiation. Future studies unlocking the mechanisms by which FL-HSCs are transiently locked into symmetric expansion, including roles for intrinsic factors and extrinsic factors from the FL niche, may facilitate efforts to engineer and expand engrafting HSCs in vitro for transplantation, gene editing, and cellular therapies. Our novel platform and transcriptomic data provide a valuable resource for such future exploration of the intricate interactions between intrinsic and extrinsic signaling networks regulating FL-HSC self-renewal at unprecedented resolution.

## Acknowledgments

Studies involving mouse models in this manuscript were supported by the American Society of Hematology Scholar Award (to B.H.), the NIH under NHLBI (K08HL140143 and R01HL168110 to B.H.), and the NIH under NIDDK (RC2DK114777 to I.D.B., C.T., S.R., and B.H. and U24DK126127 sub-award to H.G.Z.). This work was also supported by the Flow Cytometry Shared Resource, the Cellular Imaging Shared Resource (RRID:SCR_022609), the Genomics & Bioinformatics Shared Resource of the Fred Hutch/University of Washington/Seattle Children’s Cancer Consortium (NCI grant P30 CA015704), and the University of Washington Birth Defects Research Laboratory (BDRL) (NICHD grant R24HD000836). T.I is supported for overseas medical research by Takeda Science Foundation, The Nakatomi Foundation and Kitasato University School of Medicine Alumni Association. The graphical abstract was created with BioRender.

## Author contributions

Study design, experiments, collection of data and interpretation: T.I, H.G.Z, J.M, A.H, B.V, S.D, C.N.-M, K.K, R.W, O.W, C.D, D.L.J, I.P, K.A.A, S.R., C.T, I.D.B., and B.H. Original writing: T.I, A.H, B.V, H.G.Z and B.H. Article revision and editing: all authors. Supervision: B.H and H.G.Z.

## STAR★Methods

### Resource availability

#### Lead contact

Further information and requests for resources and reagents should be directed to and will be fulfilled by the lead contact, Brandon Hadland.

### Materials availability

FL-AKT-EC are available upon request.

### Experimental model and subject details

#### Mice

Wild type C57Bl6/J7 (CD45.2) and congenic C57BL/6.SJL-Ly5.1-Pep3b (CD45.1) mice were bred at the Fred Hutchinson Cancer Research Center. Male and female C57Bl6/J7 CD45.2 mice at 8-12 weeks of age were used for timed matings and transplantation experiments. All animal studies were conducted in accordance with the NIH guidelines for humane treatment of animals and were approved by the Institutional Animal Care and Use Committee at the Fred Hutchinson Cancer Center.

### Methods details

#### Dissection of embryos and cell sorting

Embryos were harvested from pregnant females and washed with PBS containing 2% FBS to minimize contaminating maternal tissues. Embryo age was determined by the date of observed maternal plugging and further confirmed by the following morphologic criteria. E13.5: Four lobes in the liver and retinal pigmentation were observed, digits were not completely separated but showed indentations. E15.5: Four lobes in the liver and retinal pigmentation were observed, digits were separated clearly and appeared unparallel. E16.5: Four lobes in the liver and retinal pigmentation were observed, digits were separated clearly and appeared parallel. Fetal livers were dissected under an inverted microscope using fine forceps, pooled in 10 ml conical tubes, and dissociated to a single cell suspension in 2% or 10% FBS with PBS by vigorous pipetting followed by passage through a 70um strainer to isolate single cells. Samples were then washed in PBS and resuspended in red blood cell lysis buffer (4L water, 33.2g Ammonium chloride (Fisher #6613), 4g sodium bicarbonate (Sigma Aldrich, S5761), and EDTA (0.1488g, Acros Organics, 147855000) or 200 uL/L of 0.5 M EDTA (Invitrogen, 15575-038)) for 5 minutes at room temperature. Cells were washed once more, pre-incubated with anti-mouse CD16/32 (FcRII blocker, BD Biosciences Cat#553141), and stained with the following monoclonal anti-mouse antibodies as described in results: GR1 FITC (clone RB6-8C5, BD Biosciences, RRID: AB_394643), F4/80 FITC (clone BM8, Biolegend, RRID: AB_893502), CD201/EPCR PE (clone eBio1560, eBioscience, RRID: AB_914317), CD45 Peridinin-Chlorophyll-Protein (PerCP) Cyanine 5.5 (clone 30-F11, Invitrogen, RRID: AB_906233), SCA1 APC (clone D7, eBioscience, RRID: AB_469488), CD144 PE-cyanine7 (PE-Cy7) (clone eBioBV13, eBioscience, RRID: AB_2573402), CD150 biotin (clone TC15-12F12.2, Biolegend, RRID: AB_345278), Streptavidin APC-eFluor780(eBioscience, RRID: AB_10366688) or Streptavidin APC-cyanine7(APC-Cy7) (BD Biosciences, RRID: AB_10054651), GR1 APCeFluor780 (clone RB6-8C5, eBioscience, RRID: AB_1518804), F4/80 APCeFluor780 (clone BM8, eBioscience, RRID: AB_2735036), CD150 PerCP-Cyanine 5.5 (clone TC5-12F12.2, Biolegend, RRID: AB_2303663), CD45 PE-Cy7 (clone 30-F11, BD Biosciences, RRID: AB_394489), ESAM FITC (clone 1GB/ESAM, Biolegend, RRID: AB_2044017). CD201 PerCP-eFluor710 (clone eBio1560, Invitrogen, RRID: AB_10718383), CD63 PE (clone NVG-2, Biolegend, RRID: AB_11203532). DAPI (Millipore, Cat#268298) was used to exclude dead cells. In some experiments, FL cells were also stained with CellTrace Far Red (ThermoFisher, Cat#C34564) following the manufacturer’s protocol. Cells were sorted by either BD FACSymphony6 or Aria II equipped with BD FACS Diva Software with index sorting capability (Becton Dickinson). For index-sorted single cells, sorting was performed in single cell mode with maximum purity mask settings to minimize contaminating cells.

#### Flow cytometric analysis of fresh FL cells

For the phenotyping of freshly isolated FL, harvested FL cells were subjected to red blood cell lysis (as described above), pre-incubated with anti-mouse CD16/32 (FcRII blocker), and stained with the following monoclonal anti-mouse antibodies: GR1 APCeFluor780 (clone RB6-8C5, eBioscience, RRID: AB_1518804), F4/80 APCeFluor780 (clone BM8, eBioscience, RRID: AB_2735036), CD201 PE (clone eBio1560, eBioscience, RRID: AB_914317), SCA1 APC (clone D7, eBioscience, RRID: AB_469488), CD45 PE-Cy7 (clone 30-F11, BD Biosciences, RRID: AB_394489), TER119 FITC (clone TER119, eBiosceicne, RRID: AB_465311), CD2 FITC (clone RM2-5, eBioscience, RRID: AB_464874), CD5 FITC (clone 53-7.3, eBioscience, RRID: AB_464909), CD8a FITC (clone 53-6.7, eBioscience, RRID: AB_469897), B220 FITC (clone RA3-6B2, BD Biosciences, RRID: AB_394618), CD48 FITC (clone HM48-1, eBioscience, RRID: AB_465078). DAPI was used to exclude dead cells. Cells were analyzed by BD FACSymphony S6 equipped with BD FACS Diva Software (Becton Dickinson) and further analyzed by Flow Jo software.

#### Generation of FL-derived Akt-EC (FL-AKT-EC)

FL-AKT-EC were generated as previously described for similar EC lines derived from murine AGM,^19,20^ which is further detailed in a protocol available at Nature Protocol Exchange (https://protocolexchange.researchsquare.com/article/pex-986/v1). Briefly, FL tissues were dissected from pooled E12 embryos and VECadherin^+^CD45^-^CD41^-^ Ter119^-^ cells were isolated by FACS. Sorted cells (>50,000 cells) were cultured on 48-well tissue culture plates coated with RetroNectin (r-fibronectin CH-296; Takara Bio Inc.) in EC media; Iscove’s Modified Dulbecco’s Medium (Gibco), 20% FBS (HyClone, fisher scientific), 1%Penicillin/streptomycin (Gibco), 1%L-glutamine (STEMCELL Technologies), heparin 0.1 mg/ml (Sigma-Aldrich), endothelial mitogen 100 μg/ml (Biomedical Technologies)*, VEGF (50 ng/ml; PeproTech), CHIR009921 (5 μM; Stemgent), and SB431542 (10 μM; R&D Systems). *EC mitogen no longer available (currently substituting rmVEGF-10ng/ml, rmFGF-10ng/ml, rmIGF-1-10ng/ml, and rmEGF-10ng/ml). Following 1–2 days culture, colonies of ECs were transduced by lentivirus with constitutively active murine AKT (PGK.myr-AKT) as previously reported.^19^ Cells were serially split, expanded in EC media and then frozen down for future use.

#### FL-AKT-EC coculture

For coculture experiments, FL-AKT-ECs at passage 12 or less were plated at a density of 1 × 10^4^ cells per well on 96-well plates 1-2 days prior to use. Prior to coculture, FL-AKT-EC layers were washed with serum-free media (X-VIVO20 (Lonza). For single cell index coculture, FL-HSCs (from donor CD45.2 mice) were individually index sorted to each well of 96-well containing FL-AKT-EC in serum-free coculture media consisting of X-VIVO 20 with recombinant cytokines (PeproTech): murine stem cell factor (SCF) at 100 ng/ml and thrombopoietin (TPO) at 20 ng/ml. Formation of hematopoietic colonies in coculture was monitored visually over time by microscopy and following various periods of coculture as indicated, a portion of cells (as indicated for each experiment) were harvested by pipetting for phenotypic analysis by flow cytometry, and in some experiments, remaining cells were used for confirmatory transplantation assays and/or scRNAseq (described below). Based on initial observations of the kinetics of colony emergence (Figure S1D), we assayed the immunophenotype of colonies between day 12-15 to ensure capture of the entire array of colony types simultaneously for the remaining experiments. When the number of viable CD45^+^ cell recorded by flow cytometry was <2, the well was classified as “no colony.” Based on initial experiments correlating colony immunophenotype with engraftment properties, colonies with greater than 80% GR1^-^ F4/80^-^SCA1^+^EPCR^+^ or CD48^-^SCA1^+^EPCR^+^ cells amongst total viable CD45^+^ cells were retrospectively classified as “HSC-like colonies.” Hematopoietic colonies with all other immunophenotypes were classified as “other colonies.” For experiments using conditioned media, X-VIVO 20 media supplemented with SCF and TPO was continuously exposed to FL-AKT-ECs (conditioned media) plated in T75 flasks. Index-sorted FL-HSCs were replenished with conditioned media every 48 hours during coculture.

#### Flow cytometric analysis of colonies

Following coculture, a fraction of the generated hematopoietic progeny in each 96-well was harvested by pipetting from the EC layer for analysis of surface phenotype by flow cytometry (unless otherwise indicated, 50% of cells generated following single cell index culture on FL-AKT-EC were used for flow cytometry). Cells were spun and re-suspended in PBS with 2% FBS, pre-incubated with anti-mouse CD16/CD32 (FcRII block) and then stained with the following anti-mouse monoclonal antibodies: GR1 FITC (clone RB6-8C5, BD Biosciences, RRID: AB_394643), F4/80 FITC (clone BM8, Biolegend, RRID: AB_893502), CD201 PE (clone eBio1560, eBioscience, RRID: AB_914317), CD45 PerCP-Cyanine5.5 (clone 30-F11, Invitrogen, RRID: AB_906233), SCA1 APC (clone D7, eBioscience, RRID: AB_469488), CD144 PE-Cy7 (clone eBioBV13, eBioscience, RRID: AB_2573402), CD45 PE-Cy7 (clone 30-F11, BD Biosciences, RRID: AB_394489), ESAM FITC (clone 1GB/ESAM, Biolegend, RRID: AB_2044017), SCA1 Alexa Fluor700 (clone D7, eBioscience, RRID: AB_657836), CD201 PerCPeFluor710(clone eBio1560, Invitrogen, RRID: AB_10718383), CD48 APC (clone HM48-1, Invitrogen, RRID: AB_469408), GR1 APCeFluor780 (clone RB6-8C5, eBioscience, RRID: AB_1518804), F4/80 APCeFluor780 (clone BM8, eBioscience, RRID: AB_2735036). DAPI was used to exclude dead cells. Cells were analyzed by either BD FACS Canto II or BD FACS LSR Fortessa equipped with BD FACS Diva Software (Becton Dickinson) and further analyzed by Flow Jo ver.10 software. For index sort analysis, Flow Jo Plugin v3.0.6 and R4.2.2 were used.

#### Live-cell imaging and image data analysis

Live-cell imaging of colony formation from single index-sorted cells was carried out on the Incucyte SX5 incubator microscope system. Brightfield and fluorescent (Celltrace-Far Red) images of whole wells were captured every 3 hours during the initial incubation period (48-96 hours). For image analysis, raw image data were exported as tiff files after background reduction, which were then used as input for a custom-developed cell tracing program in Python. The code of the celltracing program has been deposited in our lab’s Github directory for open access. In brief, the program reads in the whole time series of each well at once, identifies cell objects by fluorescent signal (CellTrace-Far Red), tracks cell divisions and hierarchical orders, and extracts parameters (eg. cell volume and cell-cell distance) for output. Python packages used in the program include *skimage*, *cv2*, *numpy*, *scipy*, *os*, and *glob*. More information about the program is available in its description on Github.

#### Transplantation assays

Recipient C57BL/6.SJL-Ly5.1-Pep3b (CD45.1) mice (6-12 weeks) were lethally irradiated with 1,000cGy using a Cesium source, and transplanted via tail vein injection. Whole bone marrow 1×10^5^ cells from C57BL/6.SJL-Ly5.1-Pep3b (CD45.1) were used for hematopoietic rescue. For single cell index assays, a fraction of the colonies harvested by pipetting was used for flow cytometric analysis (described above) and the remaining cells (or a fraction indicated in each experiment) were used for transplantation. For some experiments, a portion of the cells (as indicated for each experiment) was also used for scRNAseq studies. In limit-dilution experiments, residual (50%) cells from a single colony were serially diluted and used for transplantation. ELDA software (http://bioinf.wehi.edu.au/software/elda/) was used to calculate the number and frequency of HSCs per colony based on the fraction of mice demonstrating serial-engraftment at each dilution.^85^ Secondary transplants were performed after 24 weeks using 2×10^6^ whole bone marrow cells collected from primary recipients transplanted to lethally irradiated C57BL/6.SJL-Ly5.1-Pep3b (CD45.1) secondary recipients via the tail vein. Serial, long-term multilineage engraftment was strictly defined as donor (CD45.2) contribution to the peripheral blood with detectable contribution (>0.1%) to each lineage of donor myeloid (Gr-1 and F4/80), B cells (CD19) and T-cells (CD3) at 24 weeks post-transplant in both primary and secondary recipients. Transplant data is summarized in Table S10. We observed in initial experiments that recipients without detectable multilineage donor engraftment at 24 weeks post-transplant in primary recipients failed to provide multilineage engraftment in secondary recipients; thus, for remaining experiments, mice failing to demonstrate multilineage engraftment in primary recipients at 24 weeks were not subject to secondary transplantation. Recipient mice that died before the final timepoint analyzed for engraftment were censored for analysis of long-term serial engraftment potential but peripheral blood engraftment for these mice is shown at the ultimate timepoint analyzed before death in relevant figures (as shown in Table S10 for all transplant data).

#### Analysis of donor chimerism in recipient mice

Leukocytes from peripheral blood samples collected by retro-orbital bleeding were analyzed at the indicated time points. Lineage-specific staining for donor (CD45.2) and recipient/hematopoietic rescue (CD45.1) cells from peripheral blood was performed as previously described ^20^, using anti-mouse monoclonal antibodies: CD3 FITC (clone 17A2, BD Pharmingen, RRID: AB_395698), F4/80 PE (clone BM8, Invitrogen, RRID: AB_465923), GR1 PerCP-Cyanine 5.5 (clone RB6-8C5, BD Phrmingen, RRID: AB_394334), CD45.1 PE-Cy7 (clone A20, eBioscience, RRID:AB_469629), CD19 APC (clone 1D3/CD19, Biolegend, RRID: AB_2629839), CD45.2 APCeFluor780 (clone 104, Invitrogen, RRID: AB_1272175). Bone marrow was collected from primary recipient femur, fibula and tibiae and treated with red blood cell lysis buffer for 5 minutes at room temperature. Cells were then washed and stained using anti-mouse monoclonal antibodies: CD45.1 Brilliant Violet 510 (clone A20, BD Pharmingen, RRID: AB_2739150), CD45.2 FITC (clone 104, BD Pharmingen, RRID: AB_395041), CD2 PE (clone RM2-5, BD Pharmingen, RRID: AB_2073810), CD5 PE (clone 53-7.3, Biolegend, RRID: AB_312737), CD8a PE (clone 53-6.7, BD Pharmingen, RRID: AB_394571), GR1 PE (clone RB6-8C5, BD Pharmingen, RRID: AB_394644), TER119 PE (clone TER-119, eBioscience, RRID: AB_466043), CD11b PE (clone M1/70, BD Biosciences, RRID: AB_394775), B220 PE (clone RA3-6 B2, BD Pharmingen, RRID: AB_394620), CD150 PerCP-Cyanine 5.5 (clone TC5-12F12.2, Biolegend, RRID: AB_2303663), SCA1 APC (clone D7, eBioscience, RRID: AB_469488), CD117 APCeFluor780 (clone 2B8, eBioscience, RRID: AB_1272177), CD48 PE-Cy7 (clone HM48-1, Biolegend, RRID: AB_2075049). DAPI was used to exclude dead cells. Cells were analyzed by BD FACS Canto II equipped with BD FACS Diva Software (Becton Dickinson) and further analyzed by Flow Jo software.

#### Single cell RNA sequence experiments Freshly isolated FL-HSC and FL-AKT-EC

For single cell RNA sequencing (scRNAseq) studies, freshly sorted DAPI^-^ VECadherin^-/low^CD45^+^Gr1^-^F4/80^-^Sca1^high^EPCR^high^ (SE^hi^) cells derived from E13.5 murine fetal liver samples were subject to scRNAseq experiment. Sorted cells (3050 total) were washed twice and resuspended with PBS containing 0.04% ultrapure BSA (Invitrogen) on ice. For FL-AKT-EC, confluent ECs in a 6-well plate were cultured in serum-free coculture media (described above) overnight, harvested by treatment with TrypLE Express (Gibco), then resuspended in PBS/10% FBS at 4°C, washed twice and resuspended PBS with 0.04% ultrapure BSA in on ice. A portion of cells (targeting 3,500 cells) were subject to downstream scRNAseq assay. Cell suspensions were loaded into the Chromium Single Cell Chip G (10X Genomics) and processed in the Chromium single cell controller (10X Genomics). The 10X Genomics Version 3.1 single cell 3’ kit was used to prepare single cell mRNA libraries with the Index Kit T Set A, according to manufacturer protocols. Sequencing was performed for pooled libraries from each sample on an Illumina NextSeq 500 using the 75 cycle, high output kit, targeting a minimum of 100,000 reads per cell.

#### Cocultured FL-HSC colonies

For scRNAseq of colonies emerging following coculture, a portion of each colony was used for flow cytometry and transplantation, as indicated in each experiment, and the remaining used for scRNAseq. TotalSeq™-B0301-307 anti-mouse Hashtag Antibodies (clones M1/42; 30-F11; Biolegend, RRID: AB_ 2814067-2814072) were used to label individual colonies for multiplexing,^50^ at a concentration of 125 mg/ml (determined by titration of PE-conjugated antibodies of the identical clones), per manufacturer (10X Genomics) protocols. Cells labeled with hashing antibodies were pooled, loaded into the Chromium Single Cell Chip G (10X Genomics), and processed in the Chromium single cell controller (10X Genomics). Single cell mRNA libraries were prepared using the 10X Chromium Next GEM Single Cell3’ Reagent Kits version 3.1 with Feature Barcoding technology for Cell Surface Protein, per manufacturer protocols. Sequencing was performed for pooled libraries from each sample on an Illumina NextSeq 500 using the 75 cycle, high output kit, targeting a minimum of 35,000 reads per cell.

#### Human FL-HSCs

Human fetal livers were obtained by the Birth Defects Research Laboratory at the University of Washington following ethics board approval and maternal written consent. This study was performed in accordance with ethical and legal guidelines of the University of Washington institutional review board. Specimen age is denoted in post conception days, two weeks less than gestational age, as determined by fetal foot length measurement. Liver tissue was transferred into sterile C Tube (Miltenyi Biotec, 130-093-237) with 10 mL of cold 10% FBS in PBS and dissociated using the gentleMACS Octo Dissociator (Miltenyi Biotec, 165 rpm for 36 seconds at room temperature). The cell suspension was filtered through a 70uM cell strainer and washed 3 times with 1 mL 10% FBS in PBS. Cells were centrifuged, resuspended in 5mL RBC lysis buffer (2.0L Sterile Water, 16.6g Ammonium Chloride, 2.0g Sodium Bicarbonate, 74.4mg EDTA) and incubated at room temperature for 5 minutes. The suspension was centrifuged, resuspend with 4 mL 10% FBS in PBS, and again filtered through a 70 uM cell strainer. After centrifugation, the pellet was resuspended in 4 mL MACs buffer (MACS BSA Stock Solution Miltenyi Biotec 130-091-376, in autoMACs Rinsing Solution, Miltenyi Biotec 130-091-222) for CD34 enrichment using CD34 selection beads/kit (Miltenyi Biotech, 130-100-453) according to the manufacturer protocol. CD34-enriched cells were cryopreserved at a density of 1X10^6^/ml in freezing media (90% FBS/10% DMSO) or Cryo-stor CS 10 (Sigma, C2874). Prior to FACS, frozen samples were thawed in 37C water bath, washed with PBS with 10% FBS, and incubated with the following anti-human antibodies; APC-conjugated CD34 (clone 8G12, BD Biosciences, PRID:AB_400514), BV421-conjugated CD90 (clone 5E10, BD Biosciences, RRID:AB_2737651), PE-conjugated CD201 (clone RCR-401, Biolegend, RRID:AB_10900806), PE-Cyanine7-conjugated CD38 (clone HIT2, eBioscience, RRID:AB_1724057) and APC-eFour780-conjugated CD45RA(clone HI100, eBioscience, RRID:AB_1085364). DAPI staining was used to exclude dead cells. All reagents for cell staining were diluted in PBS with 10% FBS and staining was carried out on ice for 30 minutes. Cells were sorted to enrich for HSCs as CD34^+^CD38^-^CD45RA^-^CD90^+^EPCR^+^ on a BD FACSymphonyS6 equipped with BD FACSDiva Software. Sorted cells were washed with PBS containing 0.04% ultrapure BSA (Invitrogen) and re-suspended in 0.04% ultrapure BSA in PBS on ice. Cells were processed using the Chromium Next GEM Single Cell 5’ Reagent Kits v2 (Dual Index) (10X Genomics, PN-1000265) in accordance with the manufacturers protocol. Briefly, GEMs were generated using Chromium Next GEM Chip K (10X Genomics, PN-1000182) and processed in the Chromium Controller, targeting 1,000 to 2,500 cells per lane. Following GEM-RT clean-up and cDNA amplification, 5’ genex expression dual index library construction was performed using the Dual Index plate TT Set A. Libraries were pooled and sequenced by the Northwest Genomics Center on an Illumina NovaSeq 6000 using the SP 100 cycle, high output kit.

#### Single cell transcriptome computational analysis and quality control

The Cell Ranger pipeline v6.1.2 (10X Genomics) was used to align reads to the mm10 reference genome (for mouse samples) or to the GChr38 reference genome (for human samples) to generate the feature barcode matrix, filtering low-quality cells using default parameters. Monocle3 (v.3.1.2.9) was used for downstream analysis, combining read-depth normalized data for each group of samples.^24,86^ Uniform Manifold Approximation (UMAP) was used for dimensionality reduction.^87^ The data was mapped onto the top principal components (default settings). The alignCDS() function was applied to remove batch effects between samples using a “mutual nearest neighbor” algorithm, Batchelor (v.1.2.4).^88^ Clustering was performed using the Louvain method implemented in Monocle3.^89^ Cells with high mitochondrial genes were excluded (>10% in HSC colonies and FL-AKT-EC or >5% in fresh FL-HSCs), as were cells with low genes per cell (<1,000) and cells with low UMI per cell (UMI<10,000 for scRNAseq data in fresh FL-HSCs or UMI<3,000 for scRNAseq data in HSC colonies). Outlying clusters of cells identified as non-hematopoietic populations (lacking expression of *Ptprc*, a pan-hematopoietic gene, and expressing either *Cdh5*, an EC-specific gene, or *Hbb-bt*, a globin gene expressed by red blood cells) were removed for downstream analysis. The m3addon package was used to demultiplex the hashed samples in conjunction with Monocle3. Cells unique to each hashtag-oligo were assigned to the corresponding original colony names.

The number of UMIs and genes expressed per cell are as follows. Figure 4: UMI (median 39120, mean 40290, range 10074-92208) and genes expressed (median 6088, mean 5923, range 2513-8768). Figure 5: UMI (median: 18267, mean:19908, range 3821-49485) and genes expressed (median: 4465, mean: 4435, range 1819-6694). Figure S11: UMI (median 15701, mean 15595, range 4601-29149) and genes expressed (median 3701, mean 3599, range 1473-4970).

#### Gene module analysis

The graph_test function and the find_gene_modules function in Monocle3 were used to find genes that vary across clusters and cell types, and to group genes that have similar patterns of expressions into modules. Module genes with q-value<0.05 were selected.

#### Differential gene expression analysis

A quasipoisson distribution was used to evaluate the differential expression with the fit_model() and coefficient_table() functions in Monocle3. A q-value for multiple hypothesis testing was calculated by the Benjamini and Hochberg correction method, and q<0.05 was considered as statistically significant.

#### Gene-set scores

Gene-set scores were calculated as the log-transformed sum of the size factor-normalized expression for each gene^90^ from published signature gene-sets (Table S1) as well as the Molecular Signatures Database (https://www.gsea-msigdb.org/gsea/msigdb/index.jsp) including: Dormant HSC signature genes,^27^ Serial-engrafting HSC signature genes,^28^ diapause signature genes,^29,31^ Low output signature genes,^28^ MolO genes,^26^ REPOPSIG genes,^13^ High output signature genes,^28^ Multilineage signature genes,^28^ common stem cell dormancy genes associated with lipid metabolism,^32^ Hallmark Myc target genes (https://www.gsea-msigdb.org/gsea/msigdb/mouse/geneset/HALLMARK_MYC_TARGETS_V1.html, https://www.gsea-msigdb.org/gsea/msigdb/mouse/geneset/HALLMARK_MYC_TARGETS_V2.html), Activated HSC/MPPs, ^27,84^ WP_PURINE_METABOLISM (https://www.gsea-msigdb.org/gsea/msigdb/mouse/geneset/WP_PURINE_METABOLISM.html), WP_TCA_CYCLE (https://www.gsea-msigdb.org/gsea/msigdb/mouse/geneset/WP_TCA_CYCLE.html), HALLMARK_OXIDATIVE_PHOSPHORYLATION (https://www.gsea-msigdb.org/gsea/msigdb/mouse/geneset/HALLMARK_OXIDATIVE_PHOSPHORYLATION.html), WP_MRNA_PROCESSING (https://www.gsea-msigdb.org/gsea/msigdb/mouse/geneset/WP_MRNA_PROCESSING.html), REACTOME_TRANSLATION (https://www.gsea-msigdb.org/gsea/msigdb/mouse/geneset/REACTOME_TRANSLATION.html). Wilcoxon Rank Sum Test (ggupbr package v0.4.0) was used to calculate p values.

#### Identification of genes encoding surface proteins

To identify genes encoding putative surface proteins in scRNAseq analysis, we cross-referenced to CPSA validated surface protein^91^ (http://wlab.ethz.ch/cspa/#downloads). Only those with CSPA category defined as 1-High confidence were included.

#### Gene Ontology (GO) analysis

Gene enrichment analysis was performed using an online functional annotations tool, The AmiGO v2.5.13^92–94^ (http://geneontology.org/docs/go-citation-policy/), with the ‘GO biological process complete’ algorithm.

#### NicheNet

The nichenetr package^60^ (https://github.com/saeyslab/nichenetr/blob/master/vignettes/ligand_activity_geneset.md) was applied to scRNAseq data to identify receptor-ligand interaction betweenFL-AKT-EC or primary FL-EC and cells in the serially engrafting HSC colony (#2). All genes expressed in cells from HSC colonies (colonies #1-6) were used as the background gene-set, and genes that were significantly enriched in serially-engrafting HSC colony #2 (Table S6) were used as the gene-set of interest (i.e., genes in the receiver cell population that are potentially affected by ligands).

#### Latent variable modeling with encoded domain knowledge (MuVI)

Latent variable modeling with gene ontology or pathway gene-sets as Bayesian priors were performed using the MuVI package^35^ (https://github.com/MLO-lab/MuVI), with relevant published gene-sets (Table S10) or gene-sets downloaded from Molecular Signatures Database (https://www.gsea-msigdb.org/gsea/msigdb). scRNAseq count matrix from relevant experiments were used as input for the ‘rna’ view and MuVI was run on a single view. Max training number was 2,000 epochs, batch size was 2,000, learning rate was 0.003, 3 dense factors (random priors) were included in the mask. UMAP projection, leiden clustering, and plotting were performed using incorporated analysis and plotting modules of MuVI that are based on Scanpy (https://scanpy.readthedocs.io/).

#### Quantification and Statistical analysis

Wilcoxon signed-rank testing was used for comparison of gene-set scores in scRNAseq studies. Mann-Whitney testing was used where indicated. One-way-ANOVA with Dunnett’s multiple comparisons test was used in statistical analysis for the comparison between multiple cohorts. A *p-*value < 0.05 was considered statistically significant. GraphPad Prism (GraphPad Software, La Jolla, CA) was used for all statistical analysis except for scRNAseq studies.

#### Graphics

Some figures were created using BioRender.com.

### Data and Code Availability

- Raw sequencing data and Monocle3 cell data sets have been deposited at NCBI GEO (accession number GSE233031) and are publicly available as of the date of publication.
- All original code generated during this study has been deposited at Github and is publicly available as of the date of publication. Github repository: https://github.com/FredHutch/Ishida-etal-2023.
- Any additional information required to reanalyze the data reported in this paper is available from the lead contact (Brandon Hadland) upon request.

## Supplementary Information

**Table S1:** Gene-sets used for this study

**Table S2:** Gene modules and related gene ontology analysis of scRNAseq data from freshly isolated E13.5 FL-HSCs (related to Figure 4).

**Table S3:** Gene module analysis of scRNAseq data from E12.5 FL-HSCs (published data set, GEO GSE180050) (related to Figure S6).

**Table S4:** Gene module analysis of scRNAseq data from human FL-HSCs (related to Figure S7).

**Table S5:** Gene modules and related gene ontology analysis scRNAseq data from cocultured HSC colonies (related to Figure 5).

**Table S6:** Genes differentially expressed by serially engrafting HSC colony (#2) versus HSC-like colonies lacking long-term engraftment potential (related to Figure 5).

**Table S7:** Gene module analysis of scRNAseq data from cocultured HSC-like colonies divided by ESAM expression (related to Figure S11).

**Table S8:** Receptor-ligand pairs inferred by a comprehensive database of curated pairs of ligands and receptors (using scRNAseq data from FL-AKT-EC and HSC-like colonies) (related to Figure 6).

**Table S9:** Receptor-ligand pairs inferred by a comprehensive database of curated pairs of ligands and receptors (using published scRNAseq data from FL-EC: GEO GSE174209) (related to Figure S13).

**Table S10:** Summary of transplant data

## Supplemental Information

**Figure S1:**
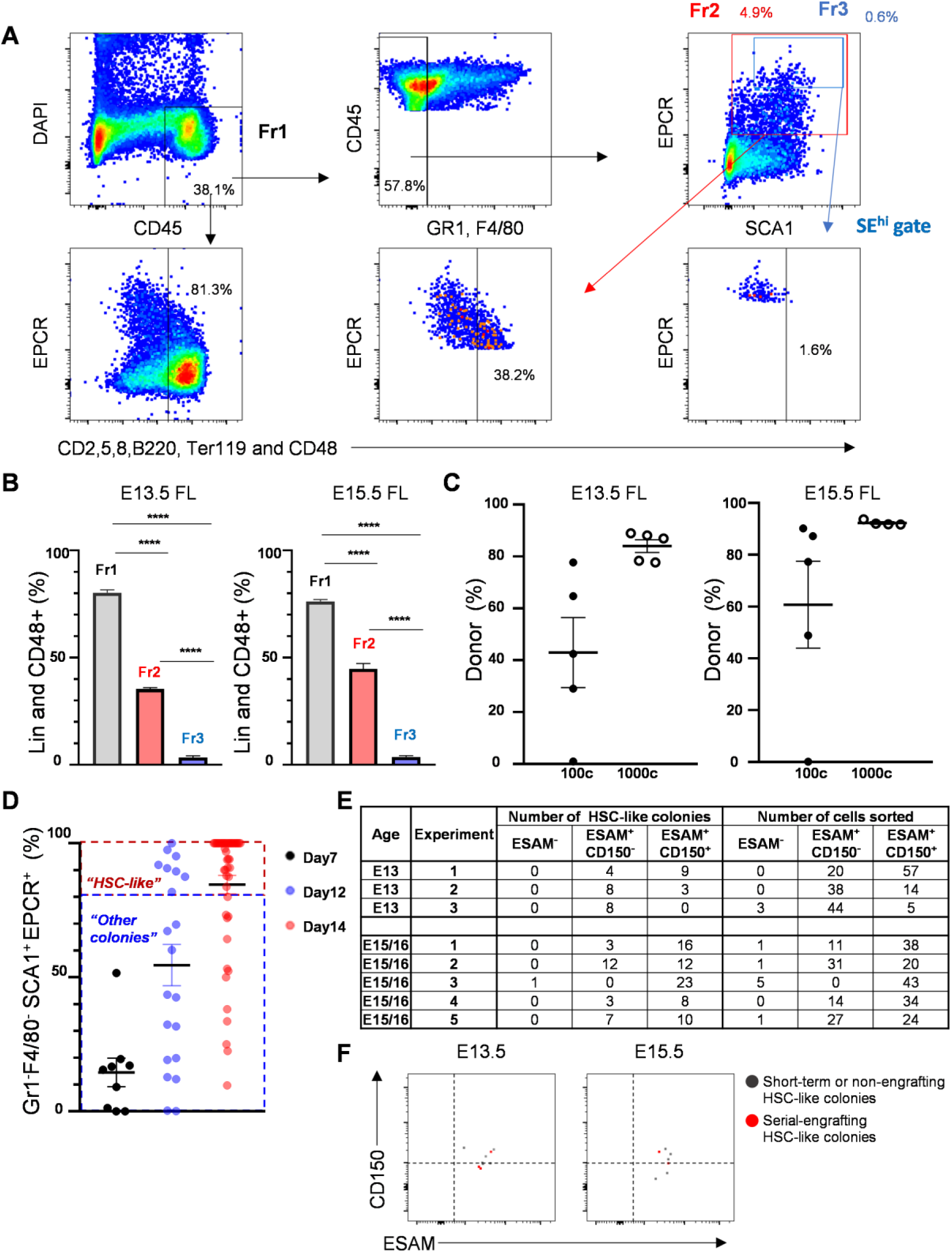
Gating strategy and index sorting of FL-HSCs. (Related to Figure 1 and Figure 2) **(A)** Expression of lineage markers (CD2, 5, 8, B220, Ter119) and CD48 in each gated fraction from freshly isolated E13.5 FL were analyzed by flow cytometry. Representative flow cytometry analysis is shown. Fr: Fraction **(B)** Frequency of cells expressing CD2, 5, 8, B220, Ter119 and CD48 in each fraction from freshly isolated E13.5 (left) and E15.5 (right) FL. N=6 (3 embryos from 2 different litters). Lin: Lineage (CD2, CD5, CD8, B220 and TER119). ****P<0.0001 (One-way ANOVA with Dunnett’s multiple comparisons test). **(C)** CD45^+^GR1^-^F4/80^-^VECadherin^-/low^SCA1^+^EPCR^+^ cells from E13.5 and E15.5 FL-HSC were sorted and transplanted into lethally irradiated mice. Donor chimerism in peripheral blood from secondary recipients at 24 weeks in E13.5 (left) and in E15.5 (right). 100 cells or 1,000 cells per recipient were transplanted. N=5 per cohort. **(D)** Frequency of GR1^-^F4/80^-^SCA1^+^EPCR^+^ cells (amongst total viable CD45^+^ cells). Single CD150^+^SE^hi^ cells from E15.5 FL were sorted and cultured with FL-AKT-EC. Colonies visually emerging at each timepoint were assessed by flow cytometry. All remaining colonies were analyzed on Day 14. n=9 (Day 7), n=21 (Day 12), n=49 (Day 14), n= 17 (no colony observed on Day14). **(E)** The distribution of HSC-like colonies and total number of cells sorted from each fraction of SE^hi^ FL-HSCs based on expression of ESAM and CD150. (n=32 HSC-like colonies from a total of 181 wells at E13.5, from 3 independent experiments; n=95 HSC-like colonies from a total of 250 wells in E15.5/16.5, from 5 independent experiments). **(F)** Expression of ESAM and CD150 by index analysis of E13.5 (left) and E15.5 (right) SE^hi^ FL-HSCs giving rise to immunophenotypic HSC-like colonies with serial multilineage engraftment (serial-engrafting HSC-like colonies, red) or lacking serial multilineage engraftment (short-term or non-engrafting HSC-like colonies, gray). E13.5: 9 HSC-like colonies were used for the transplantation. 1 colony was transplanted into 2 mice. Short-term or non-engrafting=6, serial-engrafting=3. E15.5: 7 HSC-like colonies were used for transplantation. 3 colonies were transplanted into 2 mice. Short-term or non-engrafting=5, Serial-engrafting=2.

**Figure S2:**
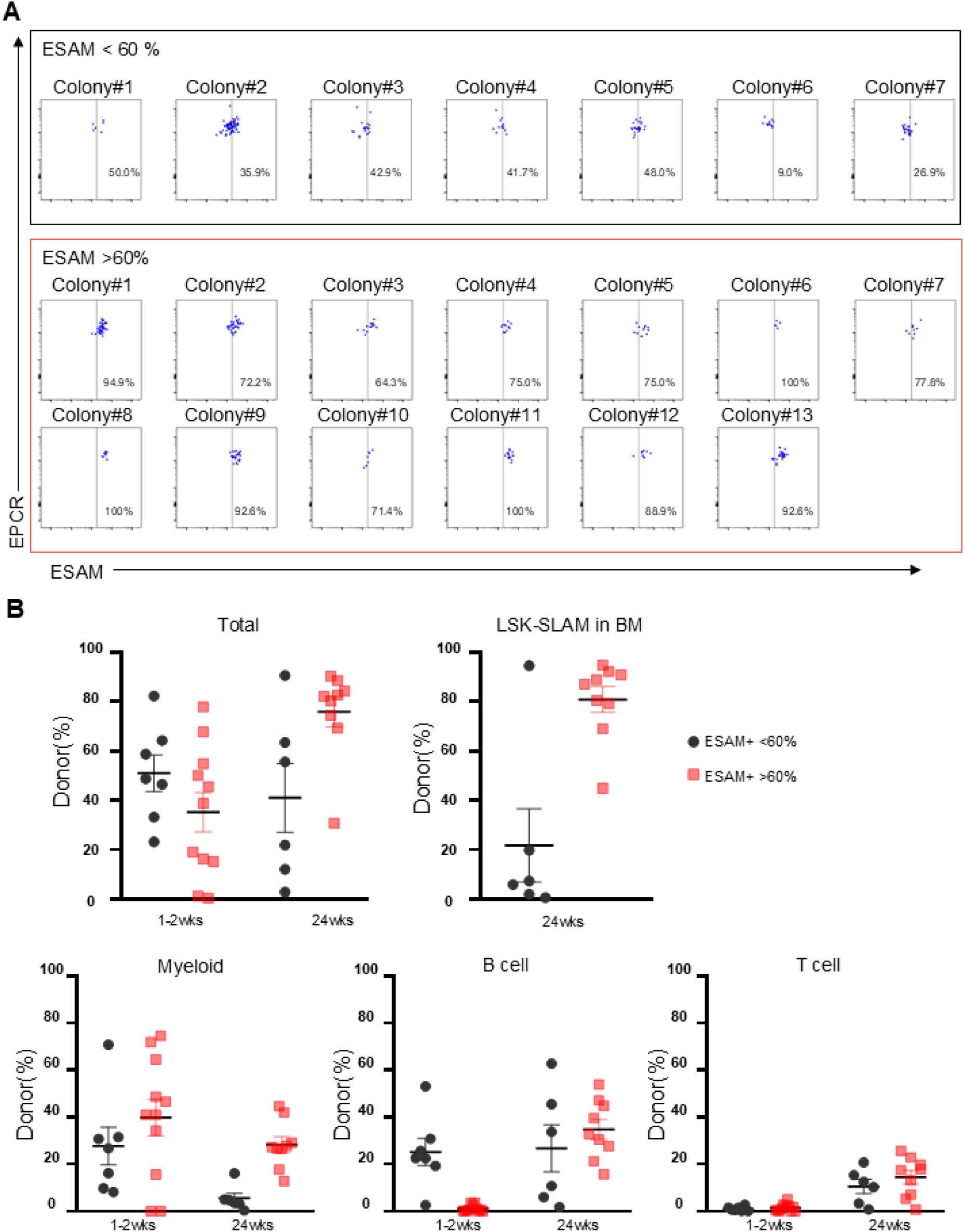
ESAM-expressing HSC-like colonies exhibit high-level contribution to immunophenotypic HSCs following transplantation. (Related to Figure 3) **(A)** Expression of ESAM in HSC-like colonies analyzed by flow cytometry following coculture (after gating cells as DAPI^-^CD45^+^CD48^-^SCA1^+^EPCR^+^). 50% of the cells from each colony were used for flow cytometry and the remaining cells from each of the 20 HSC-like colonies shown were used for transplantation. HSC-like colonies were divided based on ESAM expression (top: <60% ESAM^+^ cells, n=7 or bottom: >60% ESAM^+^ cells, n=13, in cells gated as DAPI^-^CD45^+^CD48^-^SCA1^+^EPCR^+^). **(B)** Donor chimerism (total, myeloid, B cell, and T cell in peripheral blood and LSK-SLAM: Lin^-^SCA1^+^Kit^+^CD48^-^CD150^+^ HSCs in bone marrow) of primary recipients transplanted with cells from each of the HSC-like colonies defined based on ESAM expression as shown in A (black: <60% ESAM^+^ cells, n=7; red: >60% ESAM^+^ cells, n=13).

**Figure S3:**
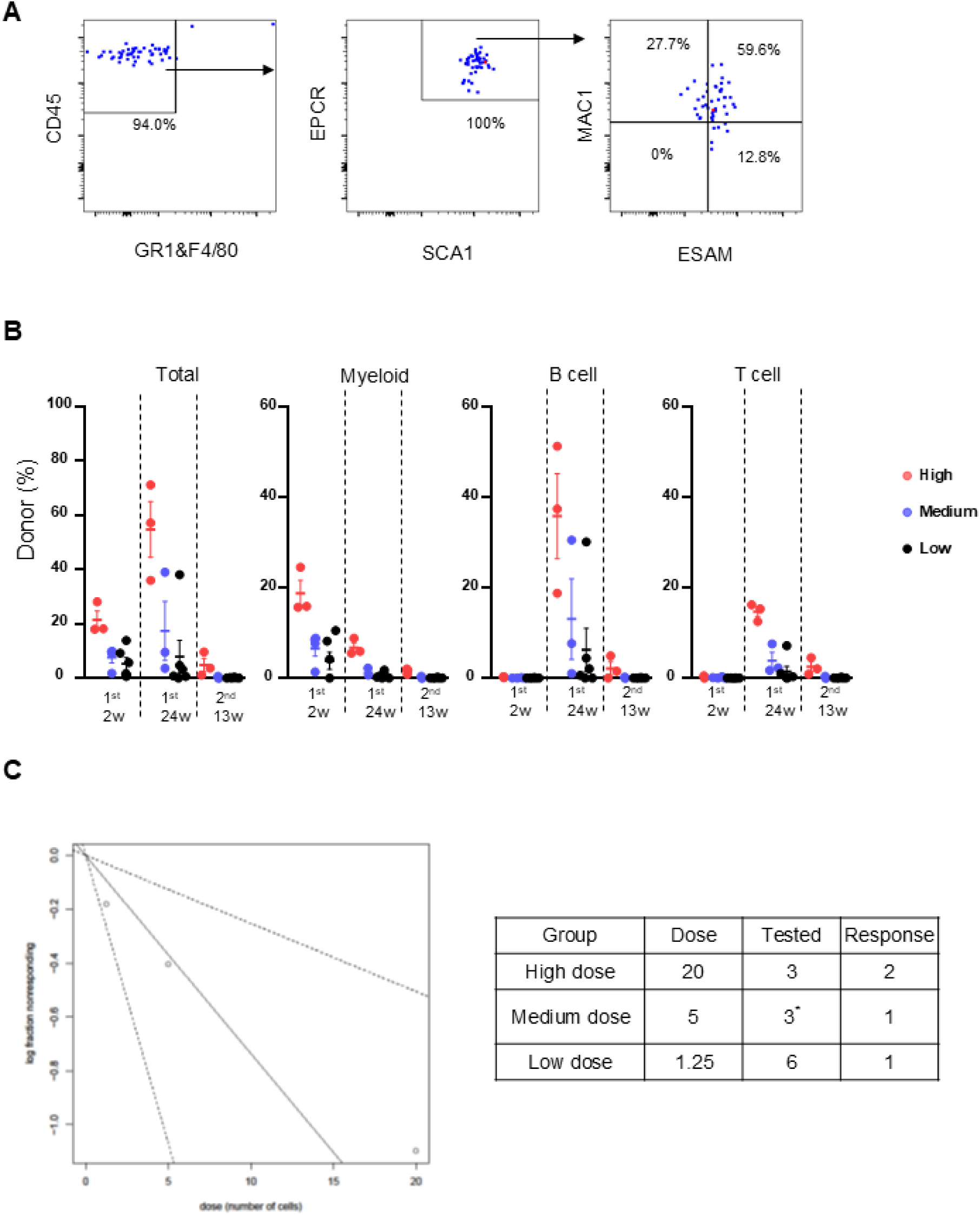
Limiting dilution transplantation demonstrates amplification of serial engrafting HSCs in an ESAM^+^ HSC colony. (Related to figure 3). **(A)** Immunophenotype of representative ESAM^+^ HSC colony tested in limiting dilution transplantation (50% of cells were analyzed by flow cytometry on Day 15). **(B)** Donor chimerism (total, myeloid, B cell, and T cell) in peripheral blood of primary and secondary recipients transplanted with limiting dilutions of cells from a single ESAM^+^ HSC colony shown in (A) (High dose = 20 cells equivalent, Medium dose = 5 cells equivalent, Low dose = 1.25 cells equivalent). **(C)** Limiting dilution calculation by ELDA (http://bioinf.wehi.edu.au/software/elda/) using the fraction of mice demonstrating serial engraftment of donor cells at each dilution^1^. Estimated HSC frequency: 1 in 13.6 cells (95% confidence interval 1 in 4.7 to 1 in 40). Estimated HSC generated: 15 (range 5-43). Engraftment was defined as >0.1% donor (CD45.2) contribution to the peripheral blood, with each of donor myeloid (Gr-1 and/or F4/80), B cell (CD19), and T cell (CD3) engraftment detected at >0.1% of peripheral blood at 13 weeks after secondary transplant. *3 out of 6 mice in medium dose cohort died before 2nd-13wks.

**Figure S4:**
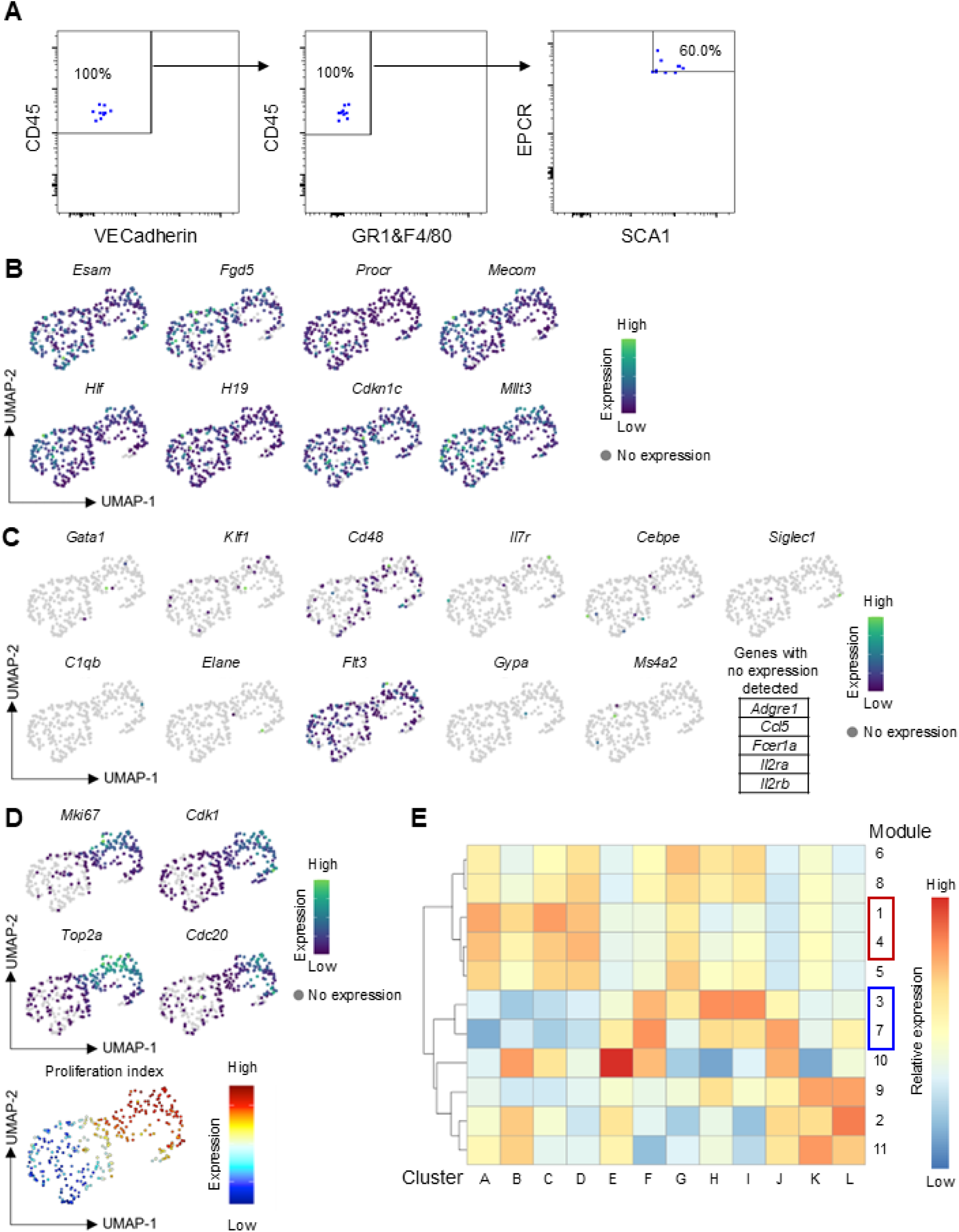
Transcriptional heterogeneity of freshly isolated FL-HSCs by scRNAseq. (Related to Figure 4) **(A)** Post-sort purity of E13.5 SE^hi^ FL-HSCs used for scRNAseq. **(B)** Gene expression heatmap for HSC-specific genes. **(C)** Gene expression heatmap for genes specific to differentiating hematopoietic progenitors and mature hematopoietic lineages. **(D)** Gene expression heatmap for cell cycle genes and proliferation index ^2–4^. **(E)** Heatmap of gene-module expression by cluster. Gene modules shown in figure 4E are highlighted in red (containing HSC-associated genes, modules 1 and 4) and blue (containing genes associated with biosynthetic/cell cycle activation, modules 3 and 7).

**Figure S5.**
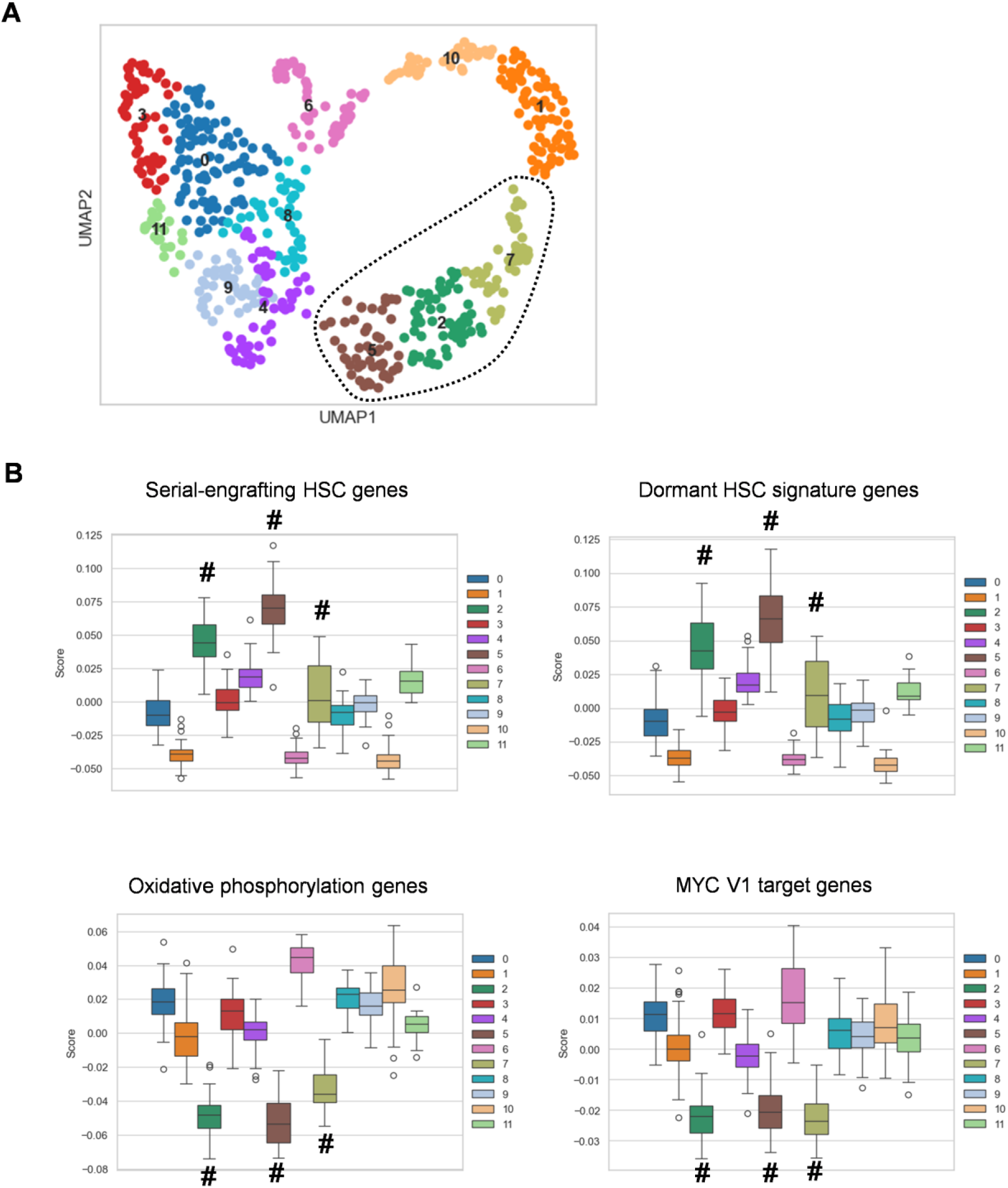
LVM of sorted SE^hi^ primary FL-HSCs using select gene sets as Bayesian priors. (Related to Figure 4) **(A)** UMAP plot of the cell clustering (leiden) in the inferred latent space representation. Numbers denote clusters, and dotted contour denotes the isolated island of cells composed of cluster 2, 5, and 7. **(B)** Quantitation of latent factor loading among clusters for each gene set, comparing how each gene set (latent factor) explains the transcriptomic features of each cluster. Cluster 2, 5, 7 in A are marked by #.

**Figure S6:**
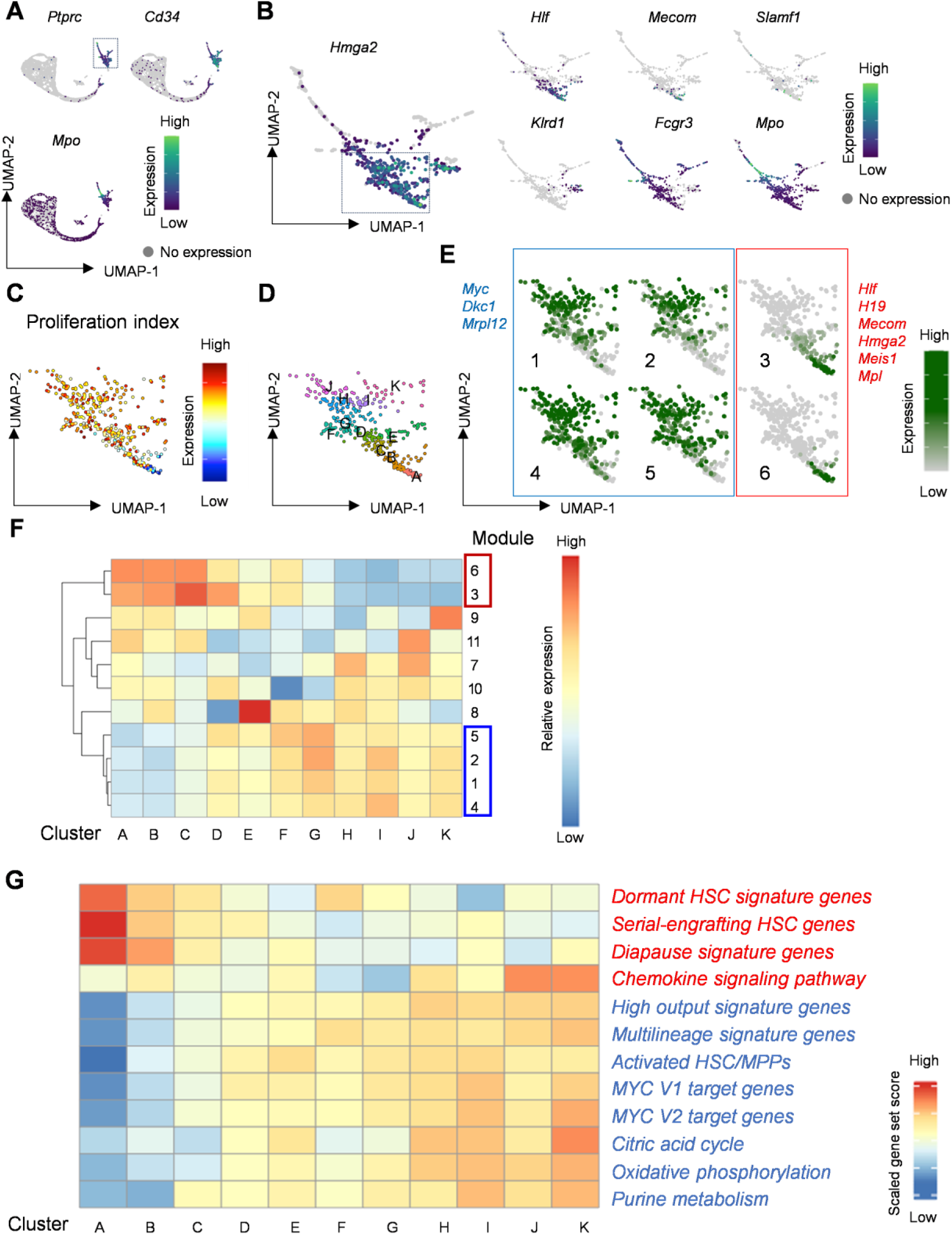
Transcriptional heterogeneity in freshly isolated E12.5 FL-HSCs from published scRNAseq data. (Related to Figure 4) **(A)** Expression of *Ptprc* (CD45), *Mpo* and *Cd34* in scRNAseq data from GSE180050.^5^ FL-HSPC expressing both *Ptprc* and *Cd34* (dashed square) were extracted for subsequent analysis (809 of 7,440 cells). **(B)** Expression of HSC-associated and lineage-specific genes in extracted FL-HSPC fraction. FL-HSCs expressing *Hmga2* (dashed square) were extracted for subsequent analysis (372 of 809 cells). **(C)** Gene expression heatmap for proliferation index. **(D)** Unsupervised clustering in UMAP. **(E)** Heatmaps for expression scores of gene modules. Gene modules 1-6 are shown with representative genes identified in each module. **(F)** Heatmap of gene-module expression by cluster. Gene modules shown in (E) are highlighted in red (containing HSC-associated genes, modules 3 and 6) and blue (containing genes associated with metabolic/cell cycle activation, modules 1,2,4 and 5). **(G)** Heatmap of gene-set scores by cluster for genes associated with HSC dormancy,^6^ serial engrafting HSCs,^7,8^ diapause,^9^ and chemokine signaling (WP_CHEMOKINE_SIGNALING_PATHWAY) or genes associated with activated HSC/MPP states including high output and multilineage signatures,^6,8,10^ Myc pathway activation (Hallmark Myc Target Genes V1, V2), and metabolic activity (WP_TCA_CYCLE, HALLMARK_OXIDATIVE_PHOSPHORY,LATION, WP_PURINE_METABOLISM). (See Table S1 for gene-sets)

**Figure S7:**
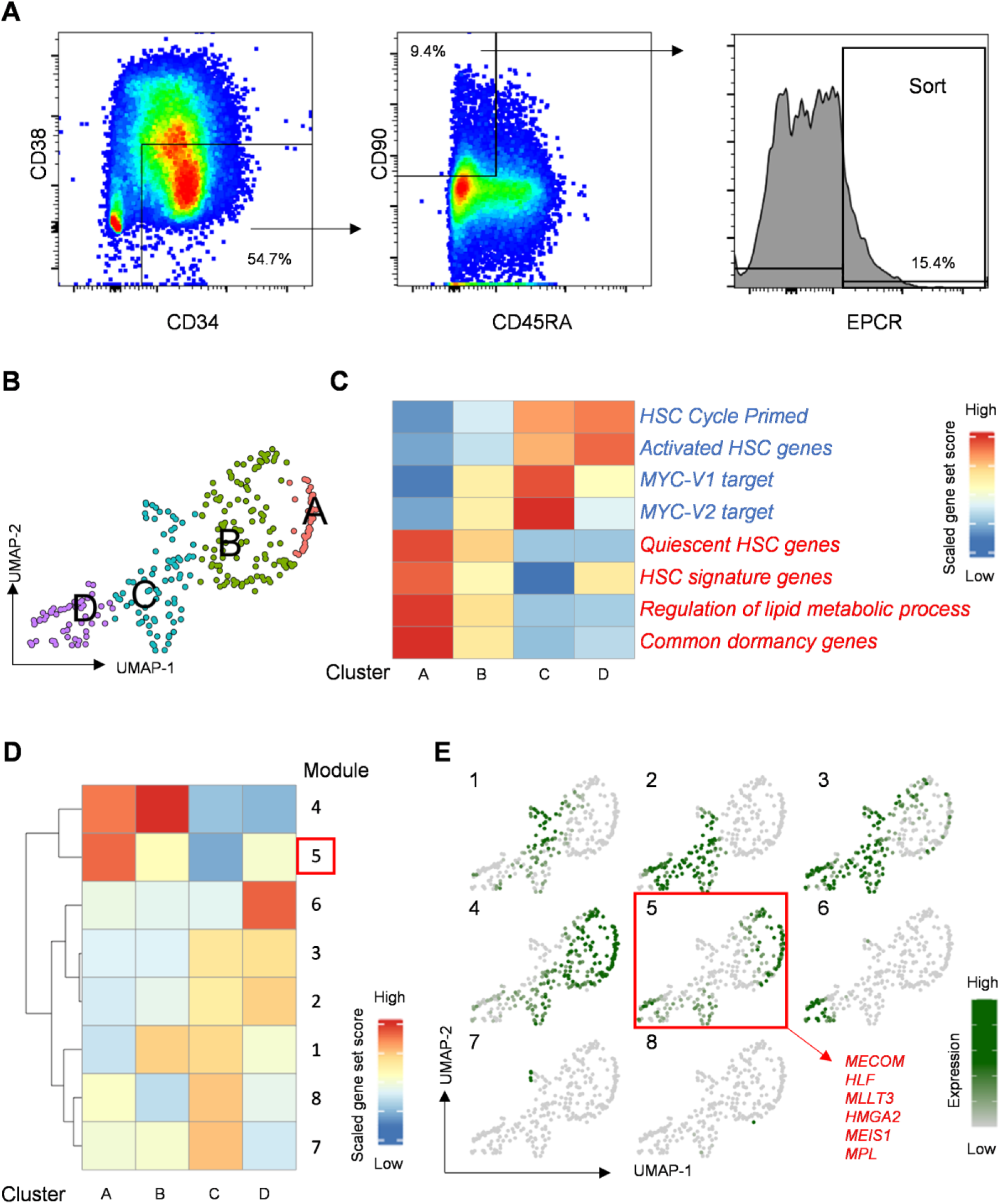
Transcriptional heterogeneity of phenotypically purified human FL-HSCs. (Related to Figure 4) **(A)** Gating strategy to sort CD34^+^38^-^45RA^-^90^+^EPCR^+^ human FL-HSCs. Freshly isolated FL samples were subject to CD34 positive cell selection and cryopreserved. Thawed samples were pooled for FACS and scRNAseq. **(B)** Unsupervised clustering in UMAP. (The number of cells in each cluster; A:116, B:80, C:55, D:41) **(C)** Heatmap of gene-set scores by cluster for genes associated with HSC activation (HSC Cycle Primed,^11^ Activated HSC^11^, HALLMARK_MYC_TARGETS_V1 and V2), HSC dormancy/quiescence^11^, HSC signature^12^, lipid metabolism (GOBP_REGULATION_OF_LIPID_METABOLIC_PROCESS), and common stem cell dormancy genes^13^ (See Table S1 for gene-sets). **(D)** Heatmap of gene-module expression by cluster. **(E)** Heatmaps for expression scores of gene modules. Gene modules 1-8 are shown. Representative HSC-associated genes are shown for module 5.

**Figure S8:**
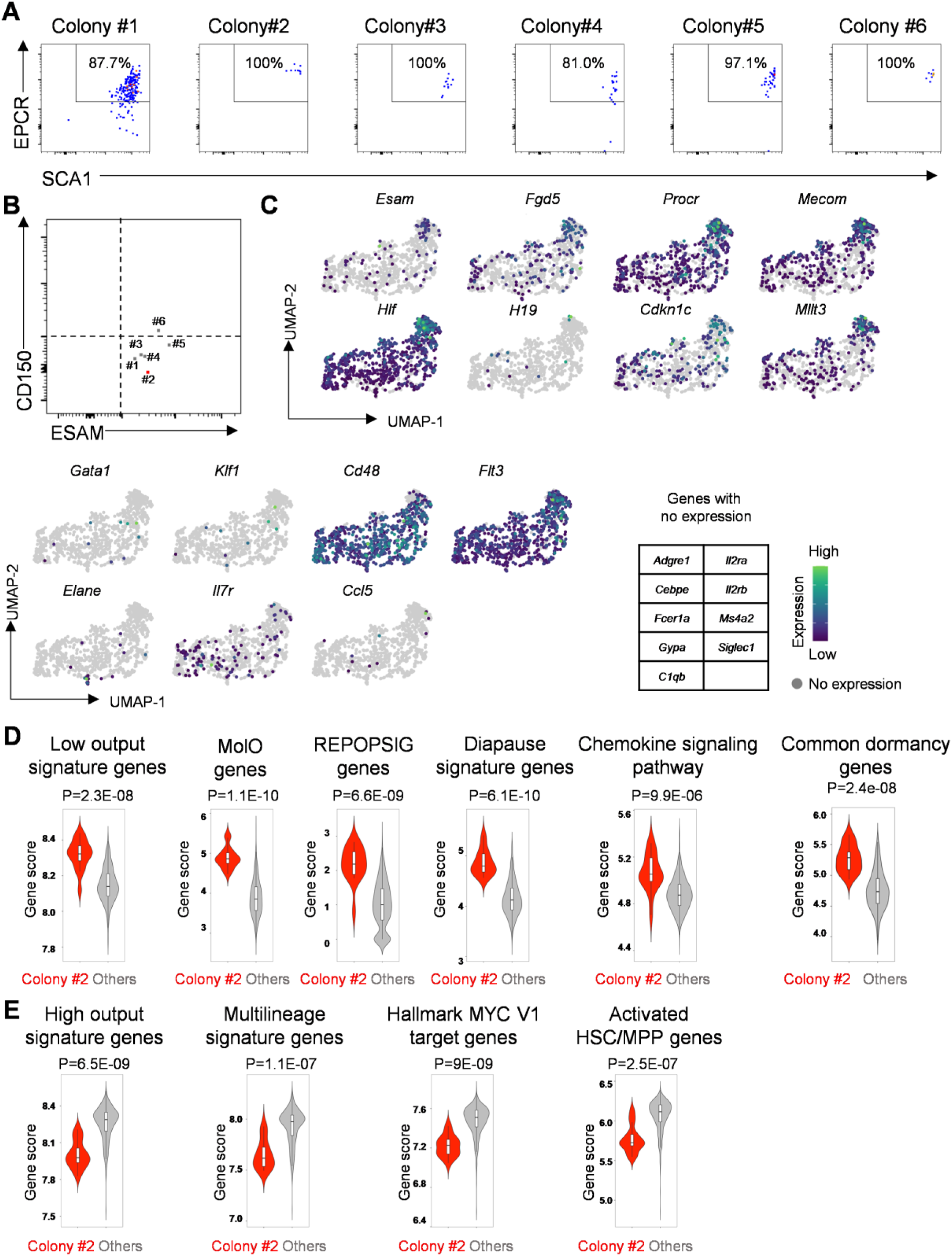
Immunophenotype and scRNAseq analysis of HSC-like colonies. (Related to Figure 5) **(A)** Flow cytometric analysis of EPCR and SCA1 expression in 6 HSC colonies used for simultaneous scRNAseq and transplantation. Cells are initially gated as viable (DAPI^-^) CD45^+^Gr1^-^F4/80^-^ after exclusion of VECadherin^+^ FL-AKT-ECs. 15% of each colony was used for phenotyping. **(B)** Index-sort profile showing surface expression of CD150 and ESAM in each of the originating SE^hi^ FL-HSCs giving rise to the indicated HSC-like colonies shown in (A). **(C)** Gene expression heatmap for genes marking HSCs (top) and differentiating progenitors/mature hematopoietic lineages (bottom). **(D)** Violin plots of gene-set scores by colony type. p values indicate Wilcoxon rank-sum test. Low output signature genes,^7^ HSC Molecular overlap population (MolO) genes,^14^ REPOPSIG genes,^15^ diapause signature genes,^9,16^ Chemokine signaling pathway (WP_CHEMOKINE_SIGNALING_PATHWAY) and common stem cell dormancy genes.^13^ (See Table S1 for gene-sets). **(E)** Violin plots of gene-set scores by colony type. p values indicate Wilcoxon rank-sum test. High output signature genes,^7^ multilineage signature genes,^7^ Myc target genes (HALLMARK_MYC_TARGETS_V1), and Activated HSC/MPP genes.^6,17^ (See Table S1 for gene-sets).

**Figure S9:**
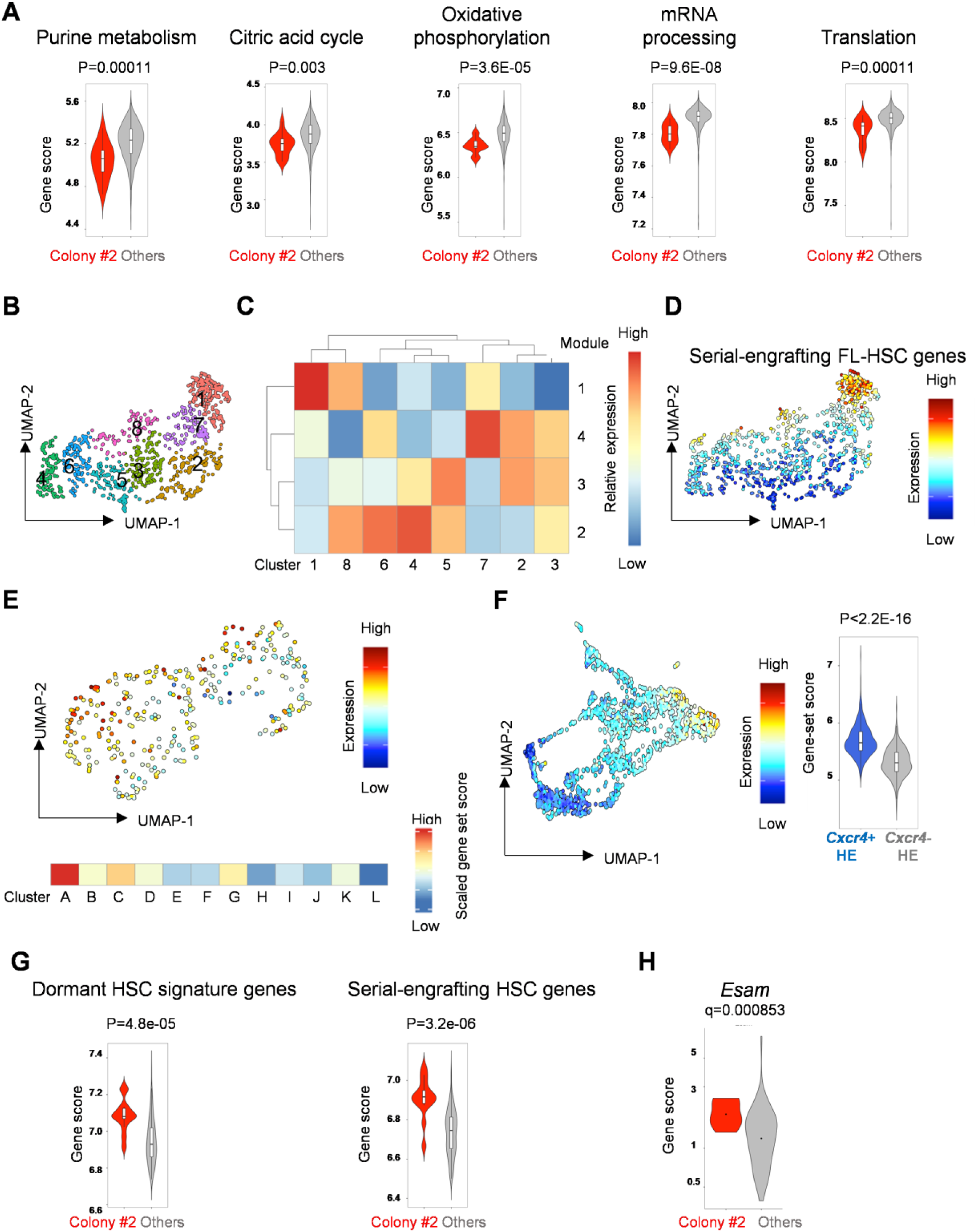
scRNAseq analysis of HSC-like colonies and correlation with freshly isolated FL-HSCs and HSC-competent hemogenic endothelium. (Related to Figure 5) **(A)** Violin plots of gene-set scores by colony type for genes associated with activated HSC/MPP metabolic states,^6^ including purine metabolism (WP_PURINE_METABOLISM), citric acid cycle (WP_TCA_CYCLE), oxidative phosphorylation (HALLMARK_OXIDATIVE_PHOSPHORYLATION), mRNA processing (WP_MRNA_PROCESSING), and translation (REACTOME_TRANSLATION). p values indicate Wilcoxon rank-sum test. (See Table S1 for gene-sets). **(B)** Unsupervised cluster analysis in UMAP. **(C)** Heatmap of gene-module expression by cluster. **(D)** Heatmap showing gene-set score of “Serially engrafting FL-HSC genes” in UMAP, identified by cross-referencing genes identified in module 1 to those identified by differential gene expression analysis (comparing colony #2 to other HSC-like colonies) (Table S1). **(E)** Heatmap showing gene-set score of “Serially engrafting FL-HSC genes” applied to scRNAseq data from freshly isolated FL-HSCs at E13.5 (Figure 4) in UMAP (top) and scaled-gene set score in each cluster (bottom). **(F)** Heatmap showing gene-set score of “Serially engrafting FL-HSC genes” applied to murine embryonic AGM hemogenic endothelium (HE) scRNAseq data^2^ in UMAP. Violin plots of gene-set scores in *Cxcr4*-expressing HE (containing HSC-competent HE) versus *Cxcr4*-negative HE (lacking HSC potential). p values indicate Wilcoxon rank-sum test. **(G)** Violin plots of gene-set scores within cluster 1 by colony type for genes associated with HSC dormancy^6^ and serial-engraftment.^7^ (See Table S1 for gene-sets). **(H)** Violin plots of *Esam* gene expression by colony type. q value was calculated using the Benjamini and Hochberg correction method.

**Figure S10:**
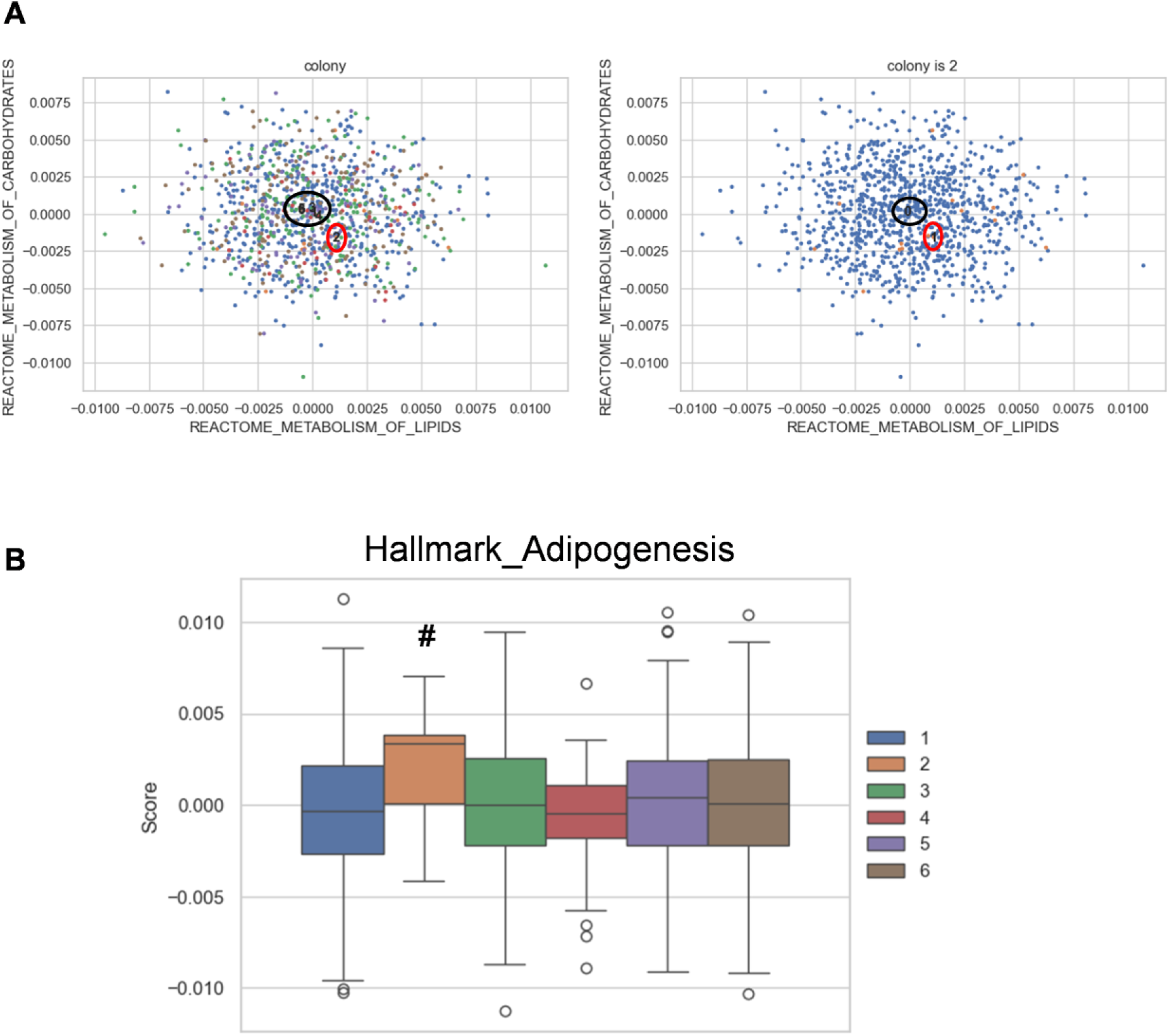
LVM analysis of scRNAseq data from HSC-like colonies using metabolism-related gene sets as Bayesian priors. (Related to Figure 5) **(A)** Scatter plot of factor scores for gene signatures “metabolism of lipids” verses “metabolism of carbohydrates” amongst cells classified by colony. A positive factor score indicates positive association with the indicated gene signature. In the left panel, the colony numbers marked on the plots represent the centroids (density centers) among the cells of each colony. In the right panel, centroid of colony 2 is marked as “1”, and the centroid of all other colonies is marked as “0”. Colony 2 (red circle) separates from the remaining colonies (black circle), suggesting distinct metabolic features with a shift toward lipid metabolism. **(B)** Quantitation of latent factor loading among colonies for the gene-set “Hallmark_Adipogenesis”. Colony 2 is marked by #.

**Figure S11:**
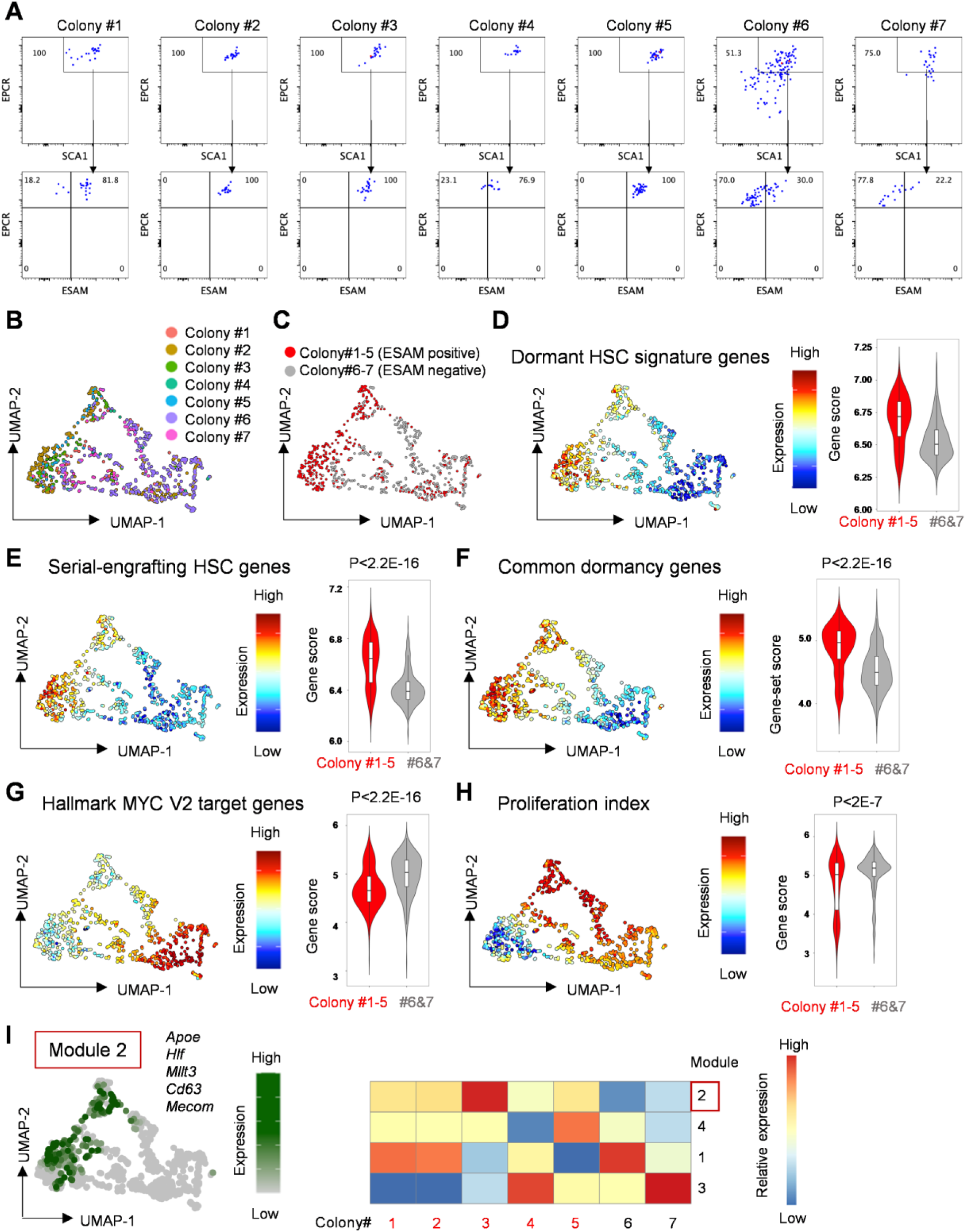
scRNAseq analysis of ESAM^+^ HSC colonies (Related to Figure 5) **(A)** Single E15.5 FL SE^hi^ cells were index sorted and cocultured with FL-AKT-EC. Flow cytometric analysis of colonies used for scRNAseq following coculture (n=7, day 13). Cells were initially gated as DAPI^-^CD45^+^Gr1^-^F4/80^-^. 25% of each colony was used for phenotyping and the remaining 75% were used for scRNAseq. **(B)** UMAP of cells labeled by colony of origin (colony #1: 56 cells, colony #2: 212 cells, colony #3: 73 cells, colony #4: 16 cells, colony #5: 40 cells, colony #6: 319 cells, colony #7: 59 cells). **(C)** UMAP of cells classified as ESAM^+^ HSC colonies (#1-5, 397 cells) verses ESAM^-^ colonies (#6-7, 378 cells). **(D)** Heatmap of gene-set scores for dormant HSC signature genes.^6^ Violin plots of gene-set scores by colony type. p values indicate Wilcoxon rank-sum test. (See Table S1 for gene-sets). **(E)** Heatmap of gene-set scores for serial-engrafting HSC genes.^7^ Violin plots of gene-set scores by colony type. p values indicate Wilcoxon rank-sum test. **(F)** Heatmap of gene-set scores for common stem cell dormancy genes.^13^ Violin plots of gene-set scores by colony type. p values indicate Wilcoxon rank-sum test. **(G)** Heatmap of gene-set scores for Myc target genes (HALLMARK_MYC_TARGETS_V2). Violin plots of gene-set scores by colony type. p values indicate Wilcoxon rank-sum test. **(H)** Heatmap of proliferation index. Violin plots of proliferation index by colony type. p values indicate Wilcoxon rank-sum test. **(I)** Expression heatmaps for modules of co-regulated genes determined using Louvain community analysis. Gene modules 2 is shown in UMAP with representative genes (left). Heatmap of expression of gene modules by colony type (right).

**Figure S12:**
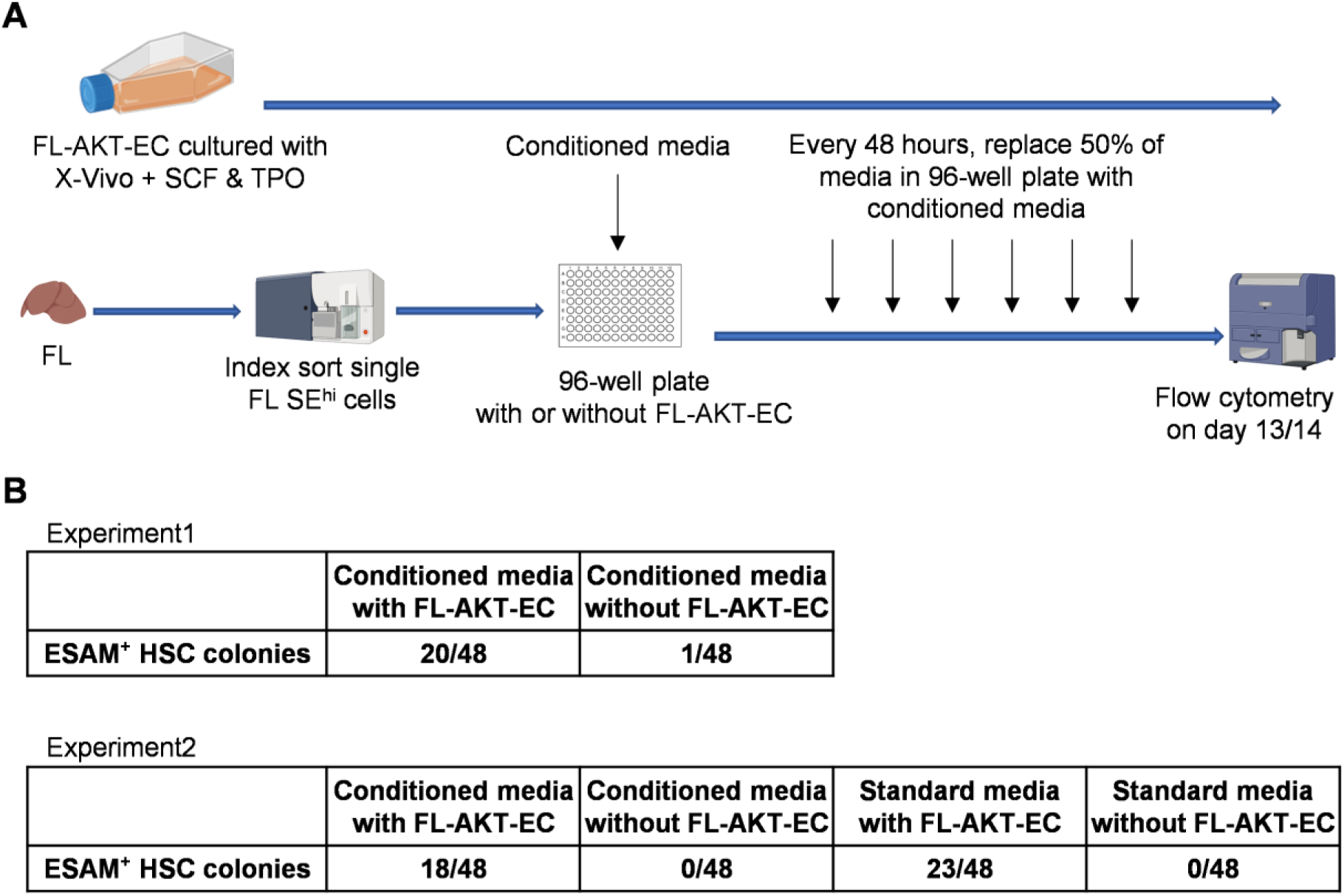
Conditioned media fails to support ESAM^+^ HSC colonies. **(A)** Overview of the experimental design. Single index sorted E15.5/16.5 FL SE^hi^ cells were cultured in the presence or absence of FL-AKT-EC. FL-AKT-EC-conditioned media (or fresh standard media) was replenished every 48 hours during culture. **(B)** The number of ESAM^+^ HSC colonies identified by flow cytometry in each experiment at end of culture (n=48 wells per cohort for each experiment).

**Figure S13:**
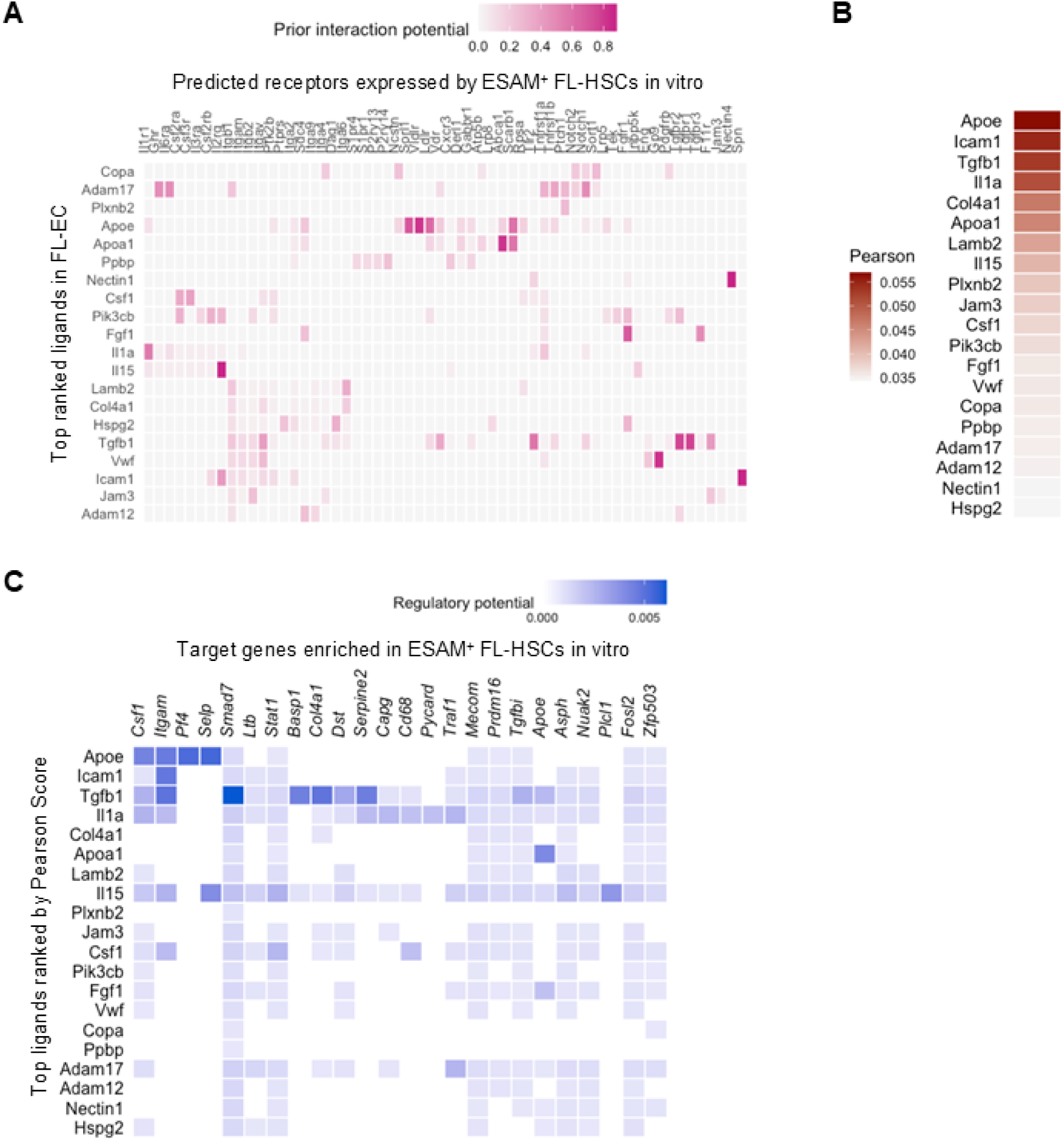
Analysis of scRNAseq data identifies candidate receptor-ligand interactions regulating self-renewal of serially engrafting FL-HSCs in the endogenous FL-EC niche. (Related to Figure 6). scRNAseq data from E14 FL-EC were obtained from GEO (GSE174209).^18^ **(A)** Heatmap showing candidate ligands (expressed by FL-EC) interacting with receptors (expressed by cells in the serially engrafting HSC colony #2) inferred by NicheNet. **(B)** Heatmap showing candidate ligands (expressed by FL-AKT-EC) ranked by Pearson correlation. **(C)** Heatmap showing the regulatory potential between the top ranked ligands (expressed by FL-EC) and downstream genes whose expression is enriched in cells from the serially engrafting HSC colony (#2) compared to cells from other HSC colonies lacking serial engraftment.

**Figure S14:**
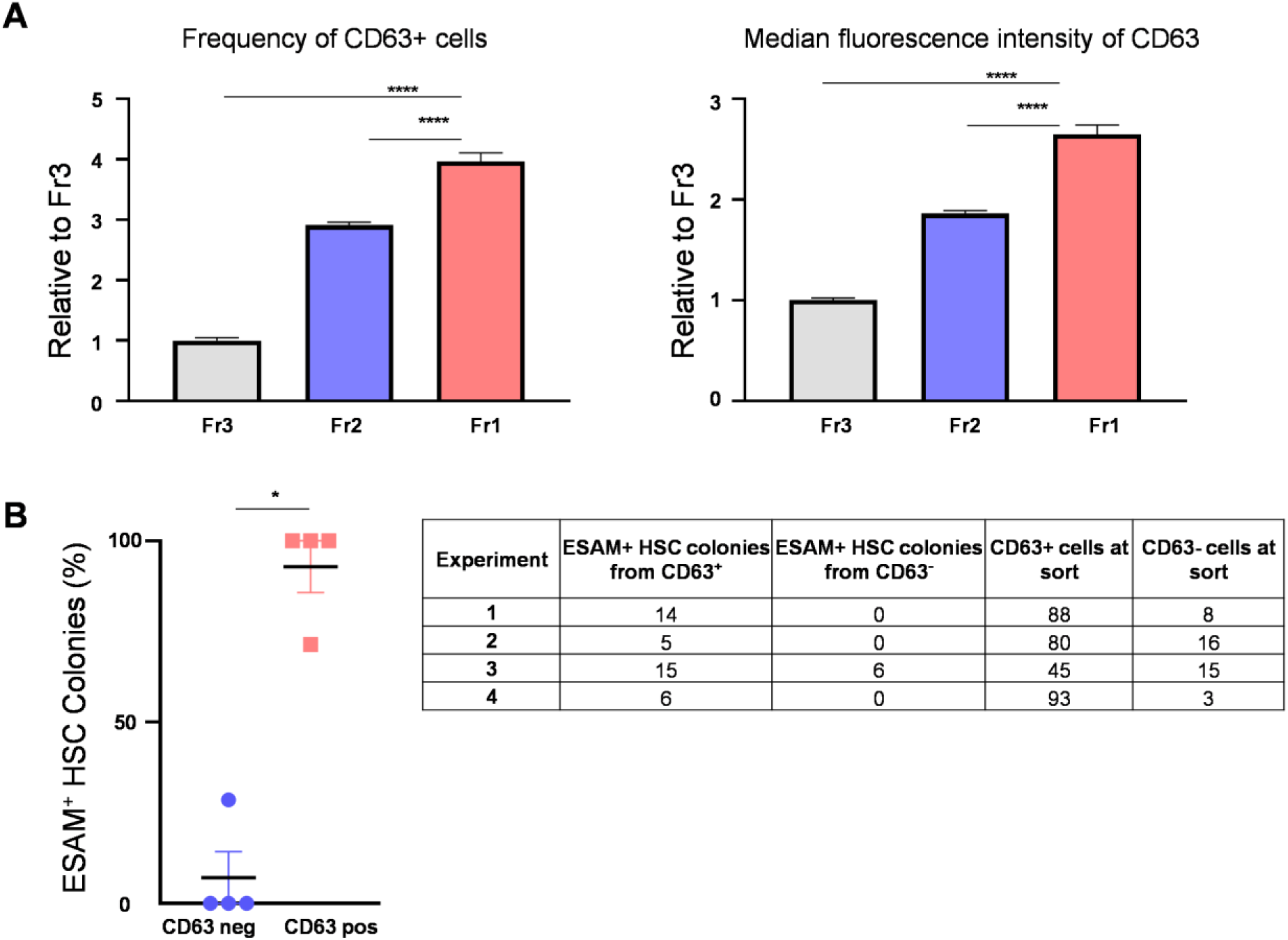
CD63 expression on FL-HSC colonies. (Related to Figure 7). **(A)** Frequency of CD63 positive cells (left) and median fluorescence intensity of CD63 (right) in each fraction of cells from Figure 7D. 3 pooled FL samples from 4 different litters were used for analysis (total n=12) in 2 independent experiments. Values are normalized to Fr3 in each experiment. One-Way ANNOVA with Dunnett’s multiple comparisons test was used for statistical comparison. **** P<0.0001 **(B)** Frequency of ESAM^+^ HSC colonies arising from CD63 negative vs CD63 positive fraction of index sorted SE^hi^ FL cells, and summary of distribution of ESAM^+^ HSC colonies based on CD63 expression at the time of index sorting. Total n=348 colonies from 4 independent experiments. Mann-Whitney testing was used for statistical analysis. *P<0.05.

## References

1. Bowie, M.B., McKnight, K.D., Kent, D.G., McCaffrey, L., Hoodless, P.A., and Eaves, C.J. (2006). Hematopoietic stem cells proliferate until after birth and show a reversible phase-specific engraftment defect. The Journal of clinical investigation 116, 2808–2816. 10.1172/jci28310.

2. Bowie, M.B., Kent, D.G., Dykstra, B., McKnight, K.D., McCaffrey, L., Hoodless, P.A., and Eaves, C.J. (2007). Identification of a new intrinsically timed developmental checkpoint that reprograms key hematopoietic stem cell properties. Proceedings of the National Academy of Sciences of the United States of America 104, 5878–5882. 10.1073/pnas.0700460104.

3. Yokomizo, T., Ideue, T., Morino-Koga, S., Tham, C.Y., Sato, T., Takeda, N., Kubota, Y., Kurokawa, M., Komatsu, N., Ogawa, M., et al. (2022). Independent origins of fetal liver haematopoietic stem and progenitor cells. Nature 609, 779–784. 10.1038/s41586-022-05203-0.

4. Dignum, T., Varnum-Finney, B., Srivatsan, S.R., Dozono, S., Waltner, O., Heck, A.M., Ishida, T., Nourigat-McKay, C., Jackson, D.L., Rafii, S., et al. (2021). Multipotent progenitors and hematopoietic stem cells arise independently from hemogenic endothelium in the mouse embryo. Cell Rep 36, 109675. 10.1016/j.celrep.2021.109675.

5. Ulloa, B.A., Habbsa, S.S., Potts, K.S., Lewis, A., McKinstry, M., Payne, S.G., Flores, J.C., Nizhnik, A., Feliz Norberto, M., Mosimann, C., and Bowman, T.V. (2021). Definitive hematopoietic stem cells minimally contribute to embryonic hematopoiesis. Cell Reports 36, 109703. 10.1016/j.celrep.2021.109703.

6. Ganuza, M., Hall, T., Finkelstein, D., Chabot, A., Kang, G., and McKinney-Freeman, S. (2017). Lifelong haematopoiesis is established by hundreds of precursors throughout mammalian ontogeny. Nature cell biology 19, 1153–1163. 10.1038/ncb3607.

7. Pei, W., Feyerabend, T.B., Rössler, J., Wang, X., Postrach, D., Busch, K., Rode, I., Klapproth, K., Dietlein, N., Quedenau, C., et al. (2017). Polylox barcoding reveals haematopoietic stem cell fates realized in vivo. Nature 548, 456–460. 10.1038/nature23653.

8. Ganuza, M., Hall, T., Myers, J., Nevitt, C., Sánchez-Lanzas, R., Chabot, A., Ding, J., Kooienga, E., Caprio, C., Finkelstein, D., et al. (2022). Murine foetal liver supports limited detectable expansion of life-long haematopoietic progenitors. Nature cell biology 24, 1475–1486. 10.1038/s41556-022-00999-5.

9. Patel, S.H., Christodoulou, C., Weinreb, C., Yu, Q., da Rocha, E.L., Pepe-Mooney, B.J., Bowling, S., Li, L., Osorio, F.G., Daley, G.Q., and Camargo, F.D. (2022). Lifelong multilineage contribution by embryonic-born blood progenitors. Nature 606, 747–753. 10.1038/s41586-022-04804-z.

10. Beaudin, A.E., Boyer, S.W., Perez-Cunningham, J., Hernandez, G.E., Derderian, S.C., Jujjavarapu, C., Aaserude, E., MacKenzie, T., and Forsberg, E.C. (2016). A Transient Developmental Hematopoietic Stem Cell Gives Rise to Innate-like B and T Cells. Cell stem cell 19, 768–783. 10.1016/j.stem.2016.08.013.

11. Benz, C., Copley, M.R., Kent, D.G., Wohrer, S., Cortes, A., Aghaeepour, N., Ma, E., Mader, H., Rowe, K., Day, C., et al. (2012). Hematopoietic stem cell subtypes expand differentially during development and display distinct lymphopoietic programs. Cell stem cell 10, 273–283. 10.1016/j.stem.2012.02.007.

12. Iwasaki, H., Arai, F., Kubota, Y., Dahl, M., and Suda, T. (2010). Endothelial protein C receptor-expressing hematopoietic stem cells reside in the perisinusoidal niche in fetal liver. Blood 116, 544–553. 10.1182/blood-2009-08-240903.

13. Che, J.L.C., Bode, D., Kucinski, I., Cull, A.H., Bain, F., Becker, H.J., Jassinskaja, M., Barile, M., Boyd, G., Belmonte, M., et al. (2022). Identification and characterization of in vitro expanded hematopoietic stem cells. 23, e55502. 10.15252/embr.202255502.

14. Hadland, B.K., Varnum-Finney, B., Mandal, P.K., Rossi, D.J., Poulos, M.G., Butler, J.M., Rafii, S., Yoder, M.C., Yoshimoto, M., and Bernstein, I.D. (2017). A Common Origin for B-1a and B-2 Lymphocytes in Clonal Pre-Hematopoietic Stem Cells. Stem Cell Reports 8, 1563–1572. 10.1016/j.stemcr.2017.04.007.

15. McKinney-Freeman, S.L., Naveiras, O., Yates, F., Loewer, S., Philitas, M., Curran, M., Park, P.J., and Daley, G.Q. (2009). Surface antigen phenotypes of hematopoietic stem cells from embryos and murine embryonic stem cells. Blood 114, 268–278. 10.1182/blood-2008-12-193888 %J Blood.

16. Kim, I., He, S., Yilmaz, O.H., Kiel, M.J., and Morrison, S.J. (2006). Enhanced purification of fetal liver hematopoietic stem cells using SLAM family receptors. Blood 108, 737–744. 10.1182/blood-2005-10-4135.

17. Papathanasiou, P., Attema, J.L., Karsunky, H., Xu, J., Smale, S.T., and Weissman, I.L. (2009). Evaluation of the long-term reconstituting subset of hematopoietic stem cells with CD150. Stem Cells 27, 2498–2508. 10.1002/stem.170.

18. McKinney-Freeman, S.L., Naveiras, O., Yates, F., Loewer, S., Philitas, M., Curran, M., Park, P.J., and Daley, G.Q. (2009). Surface antigen phenotypes of hematopoietic stem cells from embryos and murine embryonic stem cells. Blood 114, 268–278. 10.1182/blood-2008-12-193888.

19. Kobayashi, H., Butler, J.M., O’Donnell, R., Kobayashi, M., Ding, B.S., Bonner, B., Chiu, V.K., Nolan, D.J., Shido, K., Benjamin, L., and Rafii, S. (2010). Angiocrine factors from Akt-activated endothelial cells balance self-renewal and differentiation of haematopoietic stem cells. Nature cell biology 12, 1046–1056. 10.1038/ncb2108.

20. Hadland, B.K., Varnum-Finney, B., Poulos, M.G., Moon, R.T., Butler, J.M., Rafii, S., and Bernstein, I.D. (2015). Endothelium and NOTCH specify and amplify aorta-gonad-mesonephros-derived hematopoietic stem cells. J Clin Invest 125, 2032–2045. 10.1172/jci80137.

21. Ema, H., and Nakauchi, H. (2000). Expansion of hematopoietic stem cells in the developing liver of a mouse embryo. Blood 95, 2284–2288.

22. Hadland, B.K., Varnum-Finney, B., Nourigat-Mckay, C., Flowers, D., and Bernstein, I.D. (2018). Clonal Analysis of Embryonic Hematopoietic Stem Cell Precursors Using Single Cell Index Sorting Combined with Endothelial Cell Niche Co-culture. JoVE, e56973. doi:10.3791/56973.

23. Yokota, T., Oritani, K., Butz, S., Kokame, K., Kincade, P.W., Miyata, T., Vestweber, D., and Kanakura, Y. (2009). The endothelial antigen ESAM marks primitive hematopoietic progenitors throughout life in mice. Blood 113, 2914–2923. 10.1182/blood-2008-07-167106.

24. Cao, J., Spielmann, M., Qiu, X., Huang, X., Ibrahim, D.M., Hill, A.J., Zhang, F., Mundlos, S., Christiansen, L., Steemers, F.J., et al. (2019). The single-cell transcriptional landscape of mammalian organogenesis. Nature 566, 496–502. 10.1038/s41586-019-0969-x.

25. Nakamura-Ishizu, A., Ito, K., and Suda, T. (2020). Hematopoietic Stem Cell Metabolism during Development and Aging. Developmental Cell 54, 239–255. 10.1016/j.devcel.2020.06.029.

26. Wilson, N.K., Kent, D.G., Buettner, F., Shehata, M., Macaulay, I.C., Calero-Nieto, F.J., Sánchez Castillo, M., Oedekoven, C.A., Diamanti, E., Schulte, R., et al. (2015). Combined Single-Cell Functional and Gene Expression Analysis Resolves Heterogeneity within Stem Cell Populations. Cell stem cell 16, 712–724. 10.1016/j.stem.2015.04.004.

27. Cabezas-Wallscheid, N., Buettner, F., Sommerkamp, P., Klimmeck, D., Ladel, L., Thalheimer, F.B., Pastor-Flores, D., Roma, L.P., Renders, S., Zeisberger, P., et al. (2017). Vitamin A-Retinoic Acid Signaling Regulates Hematopoietic Stem Cell Dormancy. Cell 169, 807–823.e819. 10.1016/j.cell.2017.04.018.

28. Rodriguez-Fraticelli, A.E., Weinreb, C., Wang, S.W., Migueles, R.P., Jankovic, M., Usart, M., Klein, A.M., Lowell, S., and Camargo, F.D. (2020). Single-cell lineage tracing unveils a role for TCF15 in haematopoiesis. Nature 583, 585–589. 10.1038/s41586-020-2503-6.

29. Boroviak, T., Loos, R., Lombard, P., Okahara, J., Behr, R., Sasaki, E., Nichols, J., Smith, A., and Bertone, P. (2015). Lineage-Specific Profiling Delineates the Emergence and Progression of Naive Pluripotency in Mammalian Embryogenesis. Dev Cell 35, 366–382. 10.1016/j.devcel.2015.10.011.

30. Liberzon, A., Birger, C., Thorvaldsdóttir, H., Ghandi, M., Mesirov, J.P., and Tamayo, P. (2015). The Molecular Signatures Database (MSigDB) hallmark gene set collection. Cell systems 1, 417–425. 10.1016/j.cels.2015.12.004.

31. Duy, C., Li, M., Teater, M., Meydan, C., Garrett-Bakelman, F.E., Lee, T.C., Chin, C.R., Durmaz, C., Kawabata, K.C., Dhimolea, E., et al. (2021). Chemotherapy Induces Senescence-Like Resilient Cells Capable of Initiating AML Recurrence. Cancer Discovery 11, 1542–1561. 10.1158/2159-8290.Cd-20-1375.

32. van der Weijden, V.A., Stötzel, M., Iyer, D.P., Fauler, B., Gralinska, E., Shahraz, M., Meierhofer, D., Vingron, M., Rulands, S., Alexandrov, T., et al. (2024). FOXO1-mediated lipid metabolism maintains mammalian embryos in dormancy. Nat Cell Biol 26, 181–193. 10.1038/s41556-023-01325-3.

33. Argelaguet, R., Velten, B., Arnol, D., Dietrich, S., Zenz, T., Marioni, J.C., Buettner, F., Huber, W., and Stegle, O. (2018). Multi-Omics Factor Analysis-a framework for unsupervised integration of multi-omics data sets. Mol Syst Biol 14, e8124. 10.15252/msb.20178124.

34. Qoku, A., Katsaouni, N., Flinner, N., Buettner, F., and Schulz, M.H. (2023). Multimodal analysis methods in predictive biomedicine. Comput Struct Biotechnol J 21, 5829–5838. 10.1016/j.csbj.2023.11.011.

35. Qoku, A., and Buettner, F. (2023). Encoding Domain Knowledge in Multi-view Latent Variable Models: A Bayesian Approach with Structured Sparsity. In 206. (Proceedings of Machine Learning Research), pp. 11545–11562.

36. Loeffler, D., Wehling, A., Schneiter, F., Zhang, Y., Müller-Bötticher, N., Hoppe, P.S., Hilsenbeck, O., Kokkaliaris, K.D., Endele, M., and Schroeder, T. (2019). Asymmetric lysosome inheritance predicts activation of haematopoietic stem cells. Nature 573, 426–429. 10.1038/s41586-019-1531-6.

37. García-Prat, L., Kaufmann, K.B., Schneiter, F., Voisin, V., Murison, A., Chen, J., Chan-Seng-Yue, M., Gan, O.I., McLeod, J.L., Smith, S.A., et al. (2021). TFEB-mediated endolysosomal activity controls human hematopoietic stem cell fate. Cell Stem Cell 28, 1838–1850.e1810. 10.1016/j.stem.2021.07.003.

38. Alvarez, S., Díaz, M., Flach, J., Rodriguez-Acebes, S., López-Contreras, A.J., Martínez, D., Cañamero, M., Fernández-Capetillo, O., Isern, J., Passegué, E., and Méndez, J. (2015). Replication stress caused by low MCM expression limits fetal erythropoiesis and hematopoietic stem cell functionality. Nature Communications 6, 8548. 10.1038/ncomms9548.

39. Flach, J., Bakker, S.T., Mohrin, M., Conroy, P.C., Pietras, E.M., Reynaud, D., Alvarez, S., Diolaiti, M.E., Ugarte, F., Forsberg, E.C., et al. (2014). Replication stress is a potent driver of functional decline in ageing haematopoietic stem cells. Nature 512, 198–202. 10.1038/nature13619.

40. Inoue, S.-I., Noda, S., Kashima, K., Nakada, K., Hayashi, J.-I., and Miyoshi, H. (2010). Mitochondrial respiration defects modulate differentiation but not proliferation of hematopoietic stem and progenitor cells. FEBS Letters 584, 3402–3409. 10.1016/j.febslet.2010.06.036.

41. Simsek, T., Kocabas, F., Zheng, J., DeBerardinis, R.J., Mahmoud, A.I., Olson, E.N., Schneider, J.W., Zhang, C.C., and Sadek, H.A. (2010). The Distinct Metabolic Profile of Hematopoietic Stem Cells Reflects Their Location in a Hypoxic Niche. Cell stem cell 7, 380–390. 10.1016/j.stem.2010.07.011.

42. Tothova, Z., Kollipara, R., Huntly, B.J., Lee, B.H., Castrillon, D.H., Cullen, D.E., McDowell, E.P., Lazo-Kallanian, S., Williams, I.R., Sears, C., et al. (2007). FoxOs are critical mediators of hematopoietic stem cell resistance to physiologic oxidative stress. Cell 128, 325–339. 10.1016/j.cell.2007.01.003.

43. Abbas, H.A., Maccio, D.R., Coskun, S., Jackson, J.G., Hazen, A.L., Sills, T.M., You, M.J., Hirschi, K.K., and Lozano, G. (2010). Mdm2 is required for survival of hematopoietic stem cells/progenitors via dampening of ROS-induced p53 activity. Cell stem cell 7, 606–617. 10.1016/j.stem.2010.09.013.

44. Maryanovich, M., Zaltsman, Y., Ruggiero, A., Goldman, A., Shachnai, L., Zaidman, S.L., Porat, Z., Golan, K., Lapidot, T., and Gross, A. (2015). An MTCH2 pathway repressing mitochondria metabolism regulates haematopoietic stem cell fate. Nature Communications 6, 7901. 10.1038/ncomms8901.

45. Keyvani Chahi, A., Belew, M.S., Xu, J., Chen, H.T.T., Rentas, S., Voisin, V., Krivdova, G., Lechman, E., Marhon, S.A., De Carvalho, D.D., et al. (2022). PLAG1 dampens protein synthesis to promote human hematopoietic stem cell self-renewal. Blood 140, 992–1008. 10.1182/blood.2021014698 %J Blood.

46. Kruta, M., Sunshine, M.J., Chua, B.A., Fu, Y., Chawla, A., Dillingham, C.H., Hidalgo San Jose, L., De Jong, B., Zhou, F.J., and Signer, R.A.J. (2021). Hsf1 promotes hematopoietic stem cell fitness and proteostasis in response to ex vivo culture stress and aging. Cell stem cell 28, 1950–1965.e1956. 10.1016/j.stem.2021.07.009.

47. Karigane, D., Kobayashi, H., Morikawa, T., Ootomo, Y., Sakai, M., Nagamatsu, G., Kubota, Y., Goda, N., Matsumoto, M., Nishimura, Emi K., et al. (2016). p38α Activates Purine Metabolism to Initiate Hematopoietic Stem/Progenitor Cell Cycling in Response to Stress. Cell stem cell 19, 192–204. 10.1016/j.stem.2016.05.013.

48. Takubo, K., Nagamatsu, G., Kobayashi, Chiharu I., Nakamura-Ishizu, A., Kobayashi, H., Ikeda, E., Goda, N., Rahimi, Y., Johnson, Randall S., Soga, T., et al. (2013). Regulation of Glycolysis by Pdk Functions as a Metabolic Checkpoint for Cell Cycle Quiescence in Hematopoietic Stem Cells. Cell stem cell 12, 49–61. 10.1016/j.stem.2012.10.011.

49. Ceccacci, E., Villa, E., Santoro, F., Minucci, S., Ruhrberg, C., and Fantin, A. (2023). A Refined Single Cell Landscape of Haematopoiesis in the Mouse Foetal Liver. Journal of Developmental Biology 11, 15.

50. Stoeckius, M., Zheng, S., Houck-Loomis, B., Hao, S., Yeung, B.Z., Mauck, W.M., Smibert, P., and Satija, R. (2018). Cell Hashing with barcoded antibodies enables multiplexing and doublet detection for single cell genomics. Genome Biology 19, 224. 10.1186/s13059-018-1603-1.

51. Arai, F., Hirao, A., Ohmura, M., Sato, H., Matsuoka, S., Takubo, K., Ito, K., Koh, G.Y., and Suda, T. (2004). Tie2/angiopoietin-1 signaling regulates hematopoietic stem cell quiescence in the bone marrow niche. Cell 118, 149–161. 10.1016/j.cell.2004.07.004.

52. Kataoka, K., Sato, T., Yoshimi, A., Goyama, S., Tsuruta, T., Kobayashi, H., Shimabe, M., Arai, S., Nakagawa, M., Imai, Y., et al. (2011). Evi1 is essential for hematopoietic stem cell self-renewal, and its expression marks hematopoietic cells with long-term multilineage repopulating activity. J Exp Med 208, 2403–2416. 10.1084/jem.20110447.

53. Kustikova, O.S., Schwarzer, A., Stahlhut, M., Brugman, M.H., Neumann, T., Yang, M., Li, Z., Schambach, A., Heinz, N., Gerdes, S., et al. (2013). Activation of Evi1 inhibits cell cycle progression and differentiation of hematopoietic progenitor cells. Leukemia 27, 1127–1138. 10.1038/leu.2012.355.

54. Calvanese, V., Nguyen, A.T., Bolan, T.J., Vavilina, A., Su, T., Lee, L.K., Wang, Y., Lay, F.D., Magnusson, M., Crooks, G.M., et al. (2019). MLLT3 governs human haematopoietic stem-cell self-renewal and engraftment. Nature 576, 281–286. 10.1038/s41586-019-1790-2.

55. Hu, M., Lu, Y., Wang, S., Zhang, Z., Qi, Y., Chen, N., Shen, M., Chen, F., Chen, M., Yang, L., et al. (2022). CD63 acts as a functional marker in maintaining hematopoietic stem cell quiescence through supporting TGFβ signaling in mice. Cell Death & Differentiation 29, 178–191. 10.1038/s41418-021-00848-2.

56. Asai, T., Liu, Y., Di Giandomenico, S., Bae, N., Ndiaye-Lobry, D., Deblasio, A., Menendez, S., Antipin, Y., Reva, B., Wevrick, R., and Nimer, S.D. (2012). Necdin, a p53 target gene, regulates the quiescence and response to genotoxic stress of hematopoietic stem/progenitor cells. Blood 120, 1601–1612. 10.1182/blood-2011-11-393983.

57. Murphy, A.J., Akhtari, M., Tolani, S., Pagler, T., Bijl, N., Kuo, C.-L., Wang, M., Sanson, M., Abramowicz, S., Welch, C., et al. (2011). ApoE regulates hematopoietic stem cell proliferation, monocytosis, and monocyte accumulation in atherosclerotic lesions in mice. The Journal of Clinical Investigation 121, 4138–4149. 10.1172/JCI57559.

58. Hadland, B., Varnum-Finney, B., Dozono, S., Dignum, T., Nourigat-McKay, C., Heck, A.M., Ishida, T., Jackson, D.L., Itkin, T., Butler, J.M., et al. (2022). Engineering a niche supporting hematopoietic stem cell development using integrated single-cell transcriptomics. Nat Commun 13, 1584. 10.1038/s41467-022-28781-z.

59. Ramilowski, J.A., Goldberg, T., Harshbarger, J., Kloppmann, E., Lizio, M., Satagopam, V.P., Itoh, M., Kawaji, H., Carninci, P., Rost, B., and Forrest, A.R. (2015). A draft network of ligand-receptor-mediated multicellular signalling in human. Nat Commun 6, 7866. 10.1038/ncomms8866.

60. Browaeys, R., Saelens, W., and Saeys, Y. (2020). NicheNet: modeling intercellular communication by linking ligands to target genes. Nature Methods 17, 159–162. 10.1038/s41592-019-0667-5.

61. Yamazaki, S., Iwama, A., Takayanagi, S., Eto, K., Ema, H., and Nakauchi, H. (2009). TGF-beta as a candidate bone marrow niche signal to induce hematopoietic stem cell hibernation. Blood 113, 1250–1256. 10.1182/blood-2008-04-146480.

62. Yamazaki, S., Ema, H., Karlsson, G., Yamaguchi, T., Miyoshi, H., Shioda, S., Taketo, Makoto M., Karlsson, S., Iwama, A., and Nakauchi, H. (2011). Nonmyelinating Schwann Cells Maintain Hematopoietic Stem Cell Hibernation in the Bone Marrow Niche. Cell 147, 1146–1158. 10.1016/j.cell.2011.09.053.

63. Potocnik, A.J., Brakebusch, C., and Fässler, R. (2000). Fetal and adult hematopoietic stem cells require beta1 integrin function for colonizing fetal liver, spleen, and bone marrow. Immunity 12, 653–663. 10.1016/s1074-7613(00)80216-2.

64. Biswas, A., Roy, I.M., Babu, P.C., Manesia, J., Schouteden, S., Vijayakurup, V., Anto, R.J., Huelsken, J., Lacy-Hulbert, A., Verfaillie, C.M., and Khurana, S. (2020). The Periostin/Integrin-αv Axis Regulates the Size of Hematopoietic Stem Cell Pool in the Fetal Liver. Stem Cell Reports 15, 340–357. 10.1016/j.stemcr.2020.06.022.

65. Esain, V., Kwan, W., Carroll, K.J., Cortes, M., Liu, S.Y., Frechette, G.M., Sheward, L.M., Nissim, S., Goessling, W., and North, T.E. (2015). Cannabinoid Receptor-2 Regulates Embryonic Hematopoietic Stem Cell Development via Prostaglandin E2 and P-Selectin Activity. Stem Cells 33, 2596–2612. 10.1002/stem.2044.

66. Arcangeli, M.L., Frontera, V., Bardin, F., Obrados, E., Adams, S., Chabannon, C., Schiff, C., Mancini, S.J., Adams, R.H., and Aurrand-Lions, M. (2011). JAM-B regulates maintenance of hematopoietic stem cells in the bone marrow. Blood 118, 4609–4619. 10.1182/blood-2010-12-323972.

67. Gudmundsson, K.O., Nguyen, N., Oakley, K., Han, Y., Gudmundsdottir, B., Liu, P., Tessarollo, L., Jenkins, N.A., Copeland, N.G., and Du, Y. (2020). Prdm16 is a critical regulator of adult long-term hematopoietic stem cell quiescence. Proc Natl Acad Sci U S A 117, 31945–31953. 10.1073/pnas.2017626117.

68. Gómez-Salinero, J.M., Izzo, F., Lin, Y., Houghton, S., Itkin, T., Geng, F., Bram, Y., Adelson, R.P., Lu, T.M., Inghirami, G., et al. (2022). Specification of fetal liver endothelial progenitors to functional zonated adult sinusoids requires c-Maf induction. Cell Stem Cell 29, 593–609.e597. 10.1016/j.stem.2022.03.002.

69. Loeffler, D., Schneiter, F., Wang, W., Wehling, A., Kull, T., Lengerke, C., Manz, M.G., and Schroeder, T. (2022). Asymmetric organelle inheritance predicts human blood stem cell fate. Blood 139, 2011–2023. 10.1182/blood.2020009778.

70. Giebel, B., and Beckmann, J. (2007). Asymmetric cell divisions of human hematopoietic stem and progenitor cells meet endosomes. Cell Cycle 6, 2201–2204. 10.4161/cc.6.18.4658.

71. Bonora, M., Morganti, C., van Gastel, N., Ito, K., Calura, E., Zanolla, I., Ferroni, L., Zhang, Y., Jung, Y., Sales, G., et al. (2024). A mitochondrial NADPH-cholesterol axis regulates extracellular vesicle biogenesis to support hematopoietic stem cell fate. Cell Stem Cell 31, 359–377.e310. 10.1016/j.stem.2024.02.004.

72. Kobayashi, M., Wei, H., Yamanashi, T., Azevedo Portilho, N., Cornelius, S., Valiente, N., Nishida, C., Cheng, H., Latorre, A., Zheng, W.J., et al. (2023). HSC-independent definitive hematopoiesis persists into adult life. Cell Rep 42, 112239. 10.1016/j.celrep.2023.112239.

73. Cacialli, P., Mahony, C.B., Petzold, T., Bordignon, P., Rougemont, A.-L., and Bertrand, J.Y. (2021). A connexin/ifi30 pathway bridges HSCs with their niche to dampen oxidative stress. Nature Communications 12, 4484. 10.1038/s41467-021-24831-0.

74. Sigurdsson, V., Takei, H., Soboleva, S., Radulovic, V., Galeev, R., Siva, K., Leeb-Lundberg, L.M.F., Iida, T., Nittono, H., and Miharada, K. (2016). Bile Acids Protect Expanding Hematopoietic Stem Cells from Unfolded Protein Stress in Fetal Liver. Cell Stem Cell 18, 522–532. 10.1016/j.stem.2016.01.002.

75. Fares, I., Chagraoui, J., Lehnertz, B., MacRae, T., Mayotte, N., Tomellini, E., Aubert, L., Roux, P.P., and Sauvageau, G. (2017). EPCR expression marks UM171-expanded CD34(+) cord blood stem cells. Blood 129, 3344–3351. 10.1182/blood-2016-11-750729.

76. Browaeys, R., Saelens, W., and Saeys, Y. (2020). NicheNet: modeling intercellular communication by linking ligands to target genes. Nat Methods 17, 159–162. 10.1038/s41592-019-0667-5.

77. Termini, C.M., Pang, A., Li, M., Fang, T., Chang, V.Y., and Chute, J.P. (2022). Syndecan-2 enriches for hematopoietic stem cells and regulates stem cell repopulating capacity. Blood 139, 188–204. 10.1182/blood.2020010447 %J Blood.

78. Murphy, A.J., Akhtari, M., Tolani, S., Pagler, T., Bijl, N., Kuo, C.L., Wang, M., Sanson, M., Abramowicz, S., Welch, C., et al. (2011). ApoE regulates hematopoietic stem cell proliferation, monocytosis, and monocyte accumulation in atherosclerotic lesions in mice. J Clin Invest 121, 4138–4149. 10.1172/jci57559.

79. Aires, R., Porto, M.L., de Assis, L.M., Pereira, P.A.N., Carvalho, G.R., Côco, L.Z., Vasquez, E.C., Pereira, T.M.C., Campagnaro, B.P., and Meyrelles, S.S. (2021). DNA damage and aging on hematopoietic stem cells: Impact of oxidative stress in ApoE. Exp Gerontol 156, 111607. 10.1016/j.exger.2021.111607.

80. Tie, G., Messina, K.E., Yan, J., Messina, J.A., and Messina, L.M. (2014). Hypercholesterolemia induces oxidant stress that accelerates the ageing of hematopoietic stem cells. J Am Heart Assoc 3, e000241. 10.1161/JAHA.113.000241.

81. Ito, K., Carracedo, A., Weiss, D., Arai, F., Ala, U., Avigan, D.E., Schafer, Z.T., Evans, R.M., Suda, T., Lee, C.H., and Pandolfi, P.P. (2012). A PML–PPAR-δ pathway for fatty acid oxidation regulates hematopoietic stem cell maintenance. Nat Med 18, 1350–1358. 10.1038/nm.2882.

82. Ito, K., Turcotte, R., Cui, J., Zimmerman, S.E., Pinho, S., Mizoguchi, T., Arai, F., Runnels, J.M., Alt, C., Teruya-Feldstein, J., et al. (2016). Self-renewal of a purified Tie2+ hematopoietic stem cell population relies on mitochondrial clearance. Science 354, 1156–1160. 10.1126/science.aaf5530.

83. Kobayashi, H., Morikawa, T., Okinaga, A., Hamano, F., Hashidate-Yoshida, T., Watanuki, S., Hishikawa, D., Shindou, H., Arai, F., Kabe, Y., et al. (2019). Environmental Optimization Enables Maintenance of Quiescent Hematopoietic Stem Cells Ex Vivo. Cell Rep 28, 145–158.e149. 10.1016/j.celrep.2019.06.008.

84. Schönberger, K., Obier, N., Romero-Mulero, M.C., Cauchy, P., Mess, J., Pavlovich, P.V., Zhang, Y.W., Mitterer, M., Rettkowski, J., Lalioti, M.E., et al. (2021). Multilayer omics analysis reveals a non-classical retinoic acid signaling axis that regulates hematopoietic stem cell identity. Cell Stem Cell. 10.1016/j.stem.2021.10.002.

85. Hu, Y., and Smyth, G.K. (2009). ELDA: extreme limiting dilution analysis for comparing depleted and enriched populations in stem cell and other assays. J Immunol Methods 347, 70–78. 10.1016/j.jim.2009.06.008.

86. Trapnell, C., Cacchiarelli, D., Grimsby, J., Pokharel, P., Li, S., Morse, M., Lennon, N.J., Livak, K.J., Mikkelsen, T.S., and Rinn, J.L. (2014). The dynamics and regulators of cell fate decisions are revealed by pseudotemporal ordering of single cells. Nature Biotechnology 32, 381–386. 10.1038/nbt.2859.

87. Armstrong, G., Martino, C., Rahman, G., Gonzalez, A., Vázquez-Baeza, Y., Mishne, G., and Knight, R. (2021). Uniform Manifold Approximation and Projection (UMAP) Reveals Composite Patterns and Resolves Visualization Artifacts in Microbiome Data. mSystems, e0069121. 10.1128/mSystems.00691-21.

88. Haghverdi, L., Lun, A.T.L., Morgan, M.D., and Marioni, J.C. (2018). Batch effects in single-cell RNA-sequencing data are corrected by matching mutual nearest neighbors. Nat Biotechnol 36, 421–427. 10.1038/nbt.4091.

89. Levine, J.H., Simonds, E.F., Bendall, S.C., Davis, K.L., Amir el, A.D., Tadmor, M.D., Litvin, O., Fienberg, H.G., Jager, A., Zunder, E.R., et al. (2015). Data-Driven Phenotypic Dissection of AML Reveals Progenitor-like Cells that Correlate with Prognosis. Cell 162, 184–197. 10.1016/j.cell.2015.05.047.

90. Saunders, L.M., Mishra, A.K., Aman, A.J., Lewis, V.M., Toomey, M.B., Packer, J.S., Qiu, X., McFaline-Figueroa, J.L., Corbo, J.C., Trapnell, C., and Parichy, D.M. (2019). Thyroid hormone regulates distinct paths to maturation in pigment cell lineages. Elife 8. 10.7554/eLife.45181.

91. Bausch-Fluck, D., Hofmann, A., Bock, T., Frei, A.P., Cerciello, F., Jacobs, A., Moest, H., Omasits, U., Gundry, R.L., Yoon, C., et al. (2015). A mass spectrometric-derived cell surface protein atlas. PLoS One 10, e0121314. 10.1371/journal.pone.0121314.

92. Carbon, S., Ireland, A., Mungall, C.J., Shu, S., Marshall, B., and Lewis, S. (2009). AmiGO: online access to ontology and annotation data. Bioinformatics 25, 288–289. 10.1093/bioinformatics/btn615.

93. Ashburner, M., Ball, C.A., Blake, J.A., Botstein, D., Butler, H., Cherry, J.M., Davis, A.P., Dolinski, K., Dwight, S.S., Eppig, J.T., et al. (2000). Gene ontology: tool for the unification of biology. The Gene Ontology Consortium. Nat Genet 25, 25–29. 10.1038/75556.

94. Gene Ontology, C. (2021). The Gene Ontology resource: enriching a GOld mine. Nucleic Acids Res 49, D325–D334. 10.1093/nar/gkaa1113.

## Supplemental References

1. Hu, Y., and Smyth, G.K. (2009). ELDA: extreme limiting dilution analysis for comparing depleted and enriched populations in stem cell and other assays. J Immunol Methods 347, 70–78. 10.1016/j.jim.2009.06.008.

2. Dignum, T., Varnum-Finney, B., Srivatsan, S.R., Dozono, S., Waltner, O., Heck, A.M., Ishida, T., Nourigat-McKay, C., Jackson, D.L., Rafii, S., et al. (2021). Multipotent progenitors and hematopoietic stem cells arise independently from hemogenic endothelium in the mouse embryo. Cell Reports 36, 109675. 10.1016/j.celrep.2021.109675.

3. Srivatsan, S.R., McFaline-Figueroa, J.L., Ramani, V., Saunders, L., Cao, J., Packer, J., Pliner, H.A., Jackson, D.L., Daza, R.M., Christiansen, L., et al. (2020). Massively multiplex chemical transcriptomics at single-cell resolution. Science 367, 45–51. 10.1126/science.aax6234.

4. Tirosh, I., Izar, B., Prakadan, S.M., Wadsworth, M.H., 2nd, Treacy, D., Trombetta, J.J., Rotem, A., Rodman, C., Lian, C., Murphy, G., et al. (2016). Dissecting the multicellular ecosystem of metastatic melanoma by single-cell RNA-seq. Science 352, 189–196. 10.1126/science.aad0501.

5. Ceccacci, E., Villa, E., Santoro, F., Minucci, S., Ruhrberg, C., and Fantin, A. (2023). A Refined Single Cell Landscape of Haematopoiesis in the Mouse Foetal Liver. Journal of Developmental Biology 11, 15.

6. Cabezas-Wallscheid, N., Buettner, F., Sommerkamp, P., Klimmeck, D., Ladel, L., Thalheimer, F.B., Pastor-Flores, D., Roma, L.P., Renders, S., Zeisberger, P., et al. (2017). Vitamin A-Retinoic Acid Signaling Regulates Hematopoietic Stem Cell Dormancy. Cell 169, 807–823.e819. 10.1016/j.cell.2017.04.018.

7. Rodriguez-Fraticelli, A.E., Weinreb, C., Wang, S.W., Migueles, R.P., Jankovic, M., Usart, M., Klein, A.M., Lowell, S., and Camargo, F.D. (2020). Single-cell lineage tracing unveils a role for TCF15 in haematopoiesis. Nature 583, 585–589. 10.1038/s41586-020-2503-6.

8. Weinreb, C., Rodriguez-Fraticelli, A., Camargo, F.D., and Klein, A.M. (2020). Lineage tracing on transcriptional landscapes links state to fate during differentiation. Science 367. 10.1126/science.aaw3381.

9. Boroviak, T., Loos, R., Lombard, P., Okahara, J., Behr, R., Sasaki, E., Nichols, J., Smith, A., and Bertone, P. (2015). Lineage-Specific Profiling Delineates the Emergence and Progression of Naive Pluripotency in Mammalian Embryogenesis. Dev Cell 35, 366–382. 10.1016/j.devcel.2015.10.011.

10. Schönberger, K., Obier, N., Romero-Mulero, M.C., Cauchy, P., Mess, J., Pavlovich, P.V., Zhang, Y.W., Mitterer, M., Rettkowski, J., Lalioti, M.E., et al. (2022). Multilayer omics analysis reveals a non-classical retinoic acid signaling axis that regulates hematopoietic stem cell identity. Cell Stem Cell 29, 131–148.e110. 10.1016/j.stem.2021.10.002.

11. García-Prat, L., Kaufmann, K.B., Schneiter, F., Voisin, V., Murison, A., Chen, J., Chan-Seng-Yue, M., Gan, O.I., McLeod, J.L., Smith, S.A., et al. (2021). TFEB-mediated endolysosomal activity controls human hematopoietic stem cell fate. Cell Stem Cell 28, 1838–1850.e1810. 10.1016/j.stem.2021.07.003.

12. Calvanese, V., Capellera-Garcia, S., Ma, F., Fares, I., Liebscher, S., Ng, E.S., Ekstrand, S., Aguadé-Gorgorió, J., Vavilina, A., Lefaudeux, D., et al. (2022). Mapping human haematopoietic stem cells from haemogenic endothelium to birth. Nature. 10.1038/s41586-022-04571-x.

13. van der Weijden, V.A., Stötzel, M., Iyer, D.P., Fauler, B., Gralinska, E., Shahraz, M., Meierhofer, D., Vingron, M., Rulands, S., Alexandrov, T., et al. (2024). FOXO1-mediated lipid metabolism maintains mammalian embryos in dormancy. Nat Cell Biol 26, 181–193. 10.1038/s41556-023-01325-3.

14. Wilson, N.K., Kent, D.G., Buettner, F., Shehata, M., Macaulay, I.C., Calero-Nieto, F.J., Sánchez Castillo, M., Oedekoven, C.A., Diamanti, E., Schulte, R., et al. (2015). Combined Single-Cell Functional and Gene Expression Analysis Resolves Heterogeneity within Stem Cell Populations. Cell Stem Cell 16, 712–724. 10.1016/j.stem.2015.04.004.

15. Che, J.L.C., Bode, D., Kucinski, I., Cull, A.H., Bain, F., Becker, H.J., Jassinskaja, M., Barile, M., Boyd, G., Belmonte, M., et al. (2022). Identification and characterization of in vitro expanded hematopoietic stem cells. 23, e55502. 10.15252/embr.202255502.

16. Duy, C., Li, M., Teater, M., Meydan, C., Garrett-Bakelman, F.E., Lee, T.C., Chin, C.R., Durmaz, C., Kawabata, K.C., Dhimolea, E., et al. (2021). Chemotherapy Induces Senescence-Like Resilient Cells Capable of Initiating AML Recurrence. Cancer Discovery 11, 1542–1561. 10.1158/2159-8290.Cd-20-1375.

17. Schönberger, K., Obier, N., Romero-Mulero, M.C., Cauchy, P., Mess, J., Pavlovich, P.V., Zhang, Y.W., Mitterer, M., Rettkowski, J., Lalioti, M.E., et al. (2021). Multilayer omics analysis reveals a non-classical retinoic acid signaling axis that regulates hematopoietic stem cell identity. Cell Stem Cell. 10.1016/j.stem.2021.10.002.

18. Gómez-Salinero, J.M., Izzo, F., Lin, Y., Houghton, S., Itkin, T., Geng, F., Bram, Y., Adelson, R.P., Lu, T.M., Inghirami, G., et al. (2022). Specification of fetal liver endothelial progenitors to functional zonated adult sinusoids requires c-Maf induction. Cell Stem Cell 29, 593–609.e597. 10.1016/j.stem.2022.03.002.

